# Nerve and vascular abnormalities precede loss of corneal transparency in *Pax6*-haploinsufficient mice

**DOI:** 10.64898/2026.11.18.767597

**Authors:** Sneha K. Mohan, James D. Lauderdale

## Abstract

Aniridia-associated keratopathy (AAK), caused by *PAX6* haploinsufficiency, progresses from a clinically clear cornea to opacity, neovascularization, and abnormal innervation, but when disease-associated changes first emerge remains unclear. We combined corneal nerve imaging, CD31 immunolabeling, transmission electron microscopy, and RNA sequencing of whole cornea from *Pax6^Sey-Neu/+^*mice at postnatal day 3-4, day 15, and adulthood. Corneal transcripts were analyzed using curated axon-guidance, myelination, and vascular/lymphatic gene panels. Mutant corneas showed reduced nerve density and persistent stromal fascicles beginning during the first postnatal week. Peripheral vascular structures were present in both genotypes at birth; these resolved during normal maturation but persisted and extended centrally in mutant corneas by two weeks, before loss of transparency. Transcriptomic analysis identified widespread differences between genotypes while both corneas remained transparent, with the strongest and most coherent changes involving vascular and lymphatic genes. Wild-type corneas silenced an endothelial and lymphatic expression program between postnatal day 15 and adulthood, whereas mutant corneas retained expression of this program. Individual axon-guidance genes differed between genotypes, but axon-guidance did not show program-level expansion in transparent corneas. Compact stromal myelin was restricted to opaque regions. These findings identify early vascular and neural abnormalities that precede corneal opacity and establish that a transparent *Pax6*-haploinsufficient cornea is already pathologically altered.

## Introduction

Pax6 is a transcription factor required for eye development and for the formation and maintenance of the corneal and limbal epithelia, where its expression persists into adult life (Davis et al., 2003; Koroma et al., 1997; Li et al., 2015). In humans, heterozygous loss-of-function mutations in *PAX6* cause aniridia, in which aniridia-associated keratopathy (AAK) progresses over years from a clinically clear cornea to opacity, neovascularization, inflammation and epithelial barrier failure (Lagali et al., 2020; Latta et al., 2021). Mouse models of *Pax6* haploinsufficiency recapitulate key features of AAK (Collinson et al., 2004; Collinson et al., 2002; Javidjam et al., 2024; Javidjam et al., 2026; Ramaesh et al., 2003; Ramaesh et al., 2005), but it remains unclear how reduced *Pax6* dosage alters the corneal microenvironment beyond the epithelium, or when in that progression the cornea first departs from normal.

The cornea is among the most densely innervated tissues in the body (Al-Aqaba et al., 2019; Shaheen et al., 2014; Yang et al., 2018). Sensory axons from the trigeminal ganglion enter the stroma as fascicles, defasciculate into the subbasal plexus, and terminate as free nerve endings in the epithelium (Al-Aqaba et al., 2019; He and Bazan, 2016; Müller et al., 2003). During development, stromal projections extend radially across the cornea and are subsequently refined into the centripetal organization of the adult subbasal plexus, with contributions from circumferential limbal nerve bundles and the limbal plexus (Bouheraoua et al., 2019; He and Bazan, 2016; Marfurt et al., 1989; McKenna and Lwigale, 2011). In the mature cornea, sensory nerves are predominantly unmyelinated, a feature required to maintain transparency (Müller et al., 2003; Whitear, 1960). The establishment and maintenance of this architecture depend not only on neuron-intrinsic programs but also on signals provided by the surrounding tissue (Bouheraoua et al., 2019; Stepp et al., 2017; Wu et al., 2022). Experimental studies, including analyses of neuropilin-dependent guidance, demonstrate that precise corneal nerve patterning requires tightly regulated interactions between sensory axons and the corneal microenvironment (McKenna et al., 2012; McKenna et al., 2014). Guidance cues presented by the epithelium and stroma, Schwann-cell programs that regulate myelination, and mechanisms responsible for maintaining corneal avascularity therefore represent candidate pathways through which reduced *Pax6* dosage could alter corneal structure and function (Guerrero-Moreno et al., 2020; Jessen and Mirsky, 2019; Lasagni Vitar et al., 2022). Whether these candidate pathways contribute equally to disease progression, or whether one predominates, remains unknown.

In *Pax6* haploinsufficient corneas, this developmental program is disrupted from the earliest stages: early innervation dynamics are altered, with evidence of precocious trigeminal axon extension, and stromal projections are disorganized, failing to adopt the stereotyped radial alignment observed in controls (Leiper et al., 2009). The epithelium is nonetheless innervated in both genotypes, but reduced density, loss of centripetal patterning and disruption of the subbasal plexus persist into adulthood (Javidjam et al., 2024; Leiper et al., 2009). *Pax6* is therefore required not for innervation itself but for establishing and maintaining organized nerve architecture. Whether these phenotypes arise primarily from altered neuronal properties, disrupted Schwann cell interactions, or *Pax6*-dependent changes in the epithelial and stromal signaling landscape is unresolved, and whether axon–glial changes contribute to ectopic segmental myelination within the cornea has not been established.

Here we combined structural and transcriptomic analyses to define when the *Pax6*-haploinsufficient cornea first departs from normal and to identify the molecular programs associated with that transition. Because abnormalities in nerves, myelination, and vascularization have all been implicated in aniridia-associated keratopathy, we examined these processes jointly across development and adulthood using imaging, ultrastructural analysis, and RNA sequencing. By separating the effects of genotype from those associated with loss of transparency, we sought to determine whether the clinically clear mutant cornea represents an unaffected state or an already altered tissue on the path to opacity.

## Results

### Postnatal corneal nerve patterning is altered in *Pax6* haploinsufficient mice

Corneal nerve fiber patterning and density were examined during postnatal development in *Pax6*^+/+^ and *Pax6^Sey-Neu^*^/+^ mice using whole-mount TuJ1 immunolabeling (Figure 1, Supplementary Table 1). In wild-type corneas, stromal fascicles were distributed throughout the cornea at birth and progressively became restricted to the peripheral cornea and limbus during maturation, while epithelial and subbasal fibers increased in density and organization. By 4 weeks of age, a well-defined subbasal plexus had formed and early development of the characteristic corneal whorl was evident (Figure 1B-I). In contrast, *Pax6^Sey-Neu^*^/+^ corneas showed persistent stromal fascicles within central corneal regions throughout postnatal development and delayed maturation of the epithelial nerve plexus (Figure 1J-Q). Although epithelial fiber density increased between 2 and 4 weeks, subbasal fibers remained less organized and lacked the directional patterning observed in age-matched controls. Quantification demonstrated no genotype difference in nerve density at birth, but significant reductions were detected in *Pax6^Sey-Neu^*^/+^ corneas at 1, 2, and 4 weeks of age (Figure 1R-T). Similar differences were observed when central and peripheral regions were analyzed separately. Analysis of epithelial and stromal compartments independently likewise demonstrated delayed epithelial innervation and persistence of stromal fibers in mutant corneas (Supplementary Figure 1D-G). Together, these results demonstrate that reduced *Pax*6 dosage disrupts postnatal corneal nerve maturation, resulting in persistent stromal innervation, impaired subbasal organization, and reduced nerve density before the appearance of corneal opacity.

**Figure 1.**
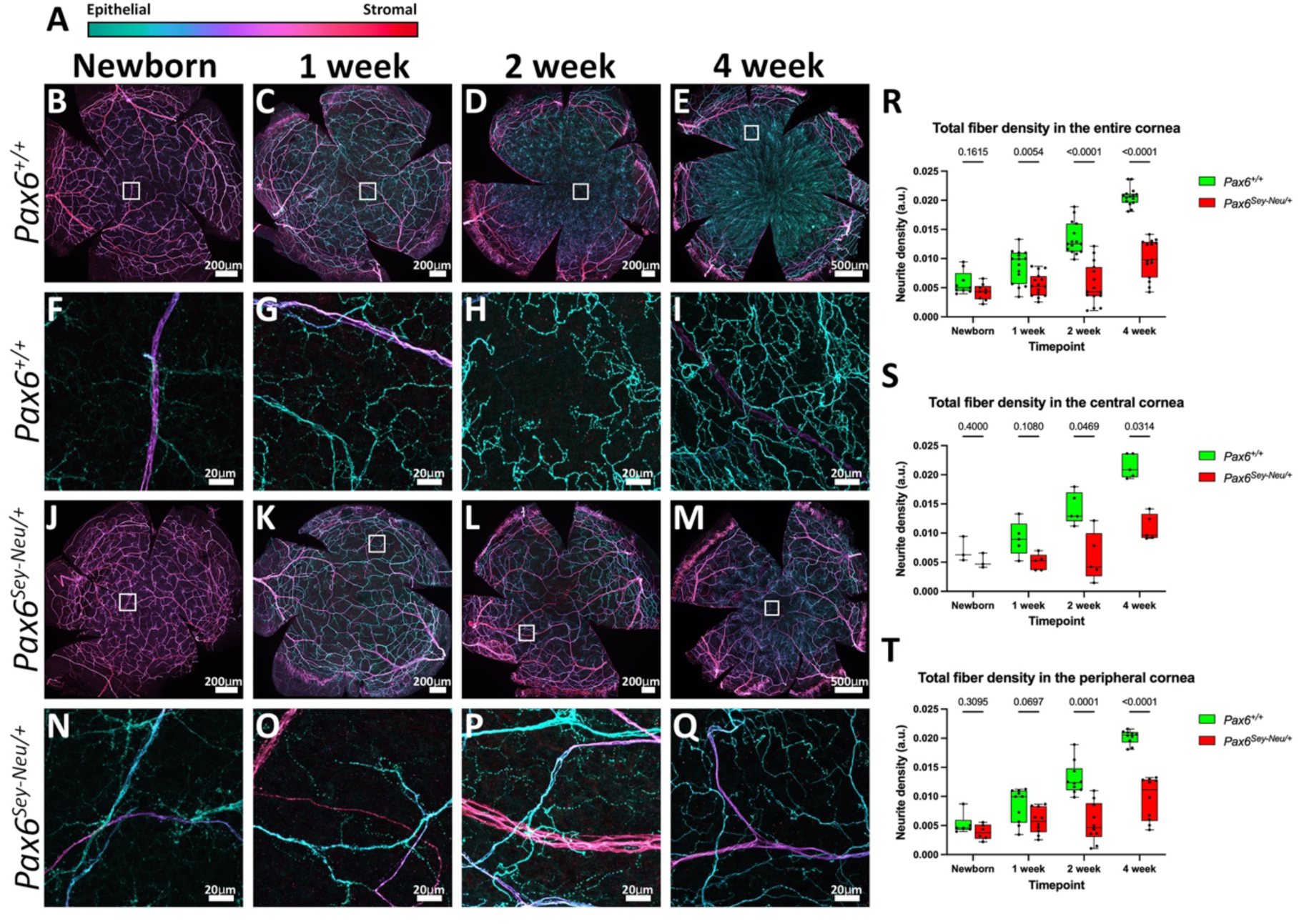
Corneal nerve development is reduced and delayed in *Pax6^Sey-Neu^*^/+^ mice. (A) Depth-encoded color scale applied to all images in B–Q. TuJ1-labeled fibers are pseudocolored by Z position within the confocal stack, from superficial/epithelial (green) to deep/stromal (red). (B–E) Corneal whole mounts from *Pax6*^+/+^ mice immunolabeled for βIII-tubulin (TuJ1) at newborn (B), 1 week (C), 2 weeks (D), and 4 weeks (E). (F–I) Higher-magnification views of the boxed regions in B–E, respectively. (J–M) Corresponding whole mounts from *Pax6^Sey-Neu^*^/+^ littermates at newborn (J), 1 week (K), 2 weeks (L), and 4 weeks (M). (N–Q) Higher-magnification views of the boxed regions in J–M, respectively. White boxes indicate the location of the higher-magnification fields. All panels are maximum-intensity projections of confocal Z-stacks spanning the full corneal thickness. The early formation of a subbasal whorl is evident in the 4-week *Pax6*^+/+^ cornea (E) but not in the age-matched mutant (M). Scale bars: 200 μm (B–D, J–L), 500 μm (E, M), and 20 μm (F–I, N–Q). Images are representative of the group; see Results for n values. (R–T) Quantification of total TuJ1^+^ nerve fiber density across the entire cornea (R), within a central ROI positioned inside the central 50% of corneal area (S), and within two equal-area peripheral ROIs positioned outside the central zone (T). Peripheral ROIs were permitted up to 20% overlap with the central zone to avoid inclusion of limbal fibers. Fiber density was quantified as thresholded particle counts normalized to ROI area and is expressed in arbitrary units (a.u.; see Methods). Each point represents one ROI measurement (three non-overlapping ROIs per cornea: one central and two peripheral; see Methods and Supplementary Table 1). Boxes indicate the median and interquartile range; whiskers indicate the minimum and maximum values. Green, *Pax6*^+/+^; red, *Pax6^Sey-Neu^*^/+^. Genotypes were compared at each age using multiple two-tailed Mann-Whitney tests with Holm-Šídák correction for multiple comparisons. Adjusted P values are shown above comparisons.

### Adult *Pax6* haploinsufficient corneas exhibit heterogeneous nerve patterning and reduced density

Corneal nerve patterning and density were examined in adult mice (Supplementary Methods) using whole-mount TuJ1 immunolabeling (Figure 2). In *Pax6*^+/+^ corneas, stromal fascicles were largely confined to the limbal and peripheral regions, while long, organized subbasal fibers formed a characteristic central whorl (Figure 2B, C). Higher-magnification imaging revealed a dense and uniformly oriented subbasal plexus. In contrast, *Pax6^Sey-Neu^*^/+^ corneas displayed marked heterogeneity in nerve architecture, both between corneas and within individual corneas (Figure 2D-I). Mutant corneas ranged from sparse clusters of subbasal fibers to extensive stromal innervation, with many exhibiting mixed phenotypes in which distinct regions of the same cornea differed substantially in nerve organization. In some corneas, stromal fascicles extended into central regions, whereas in others they were restricted to peripheral or sector-specific domains. Higher-magnification imaging revealed reduced subbasal organization and persistence of fibers occupying deeper stromal planes (Supplementary Figure 2), although localized whorl-like organization persisted in a subset of mutant corneas (Supplementary Figure 3). Quantification demonstrated a significant reduction in total nerve density in adult *Pax6^Sey-Neu^*^/+^ corneas relative to *Pax6*^+/+^ controls (Figure 2J). This reduction was most pronounced in the peripheral cornea (Figure 2L), whereas central corneal density did not differ significantly between genotypes (Figure 2K). Additional analyses of total and epithelial fiber density are shown in Supplementary Figure 1H–K. Together, these findings demonstrate that abnormal corneal innervation persists into adulthood in *Pax6^Sey-Neu^*^/+^ mice and is characterized by both reduced nerve density and substantial spatial heterogeneity in nerve architecture. Given this heterogeneity, we next asked whether differences in nerve architecture were associated with corneal transparency.

**Figure 2.**
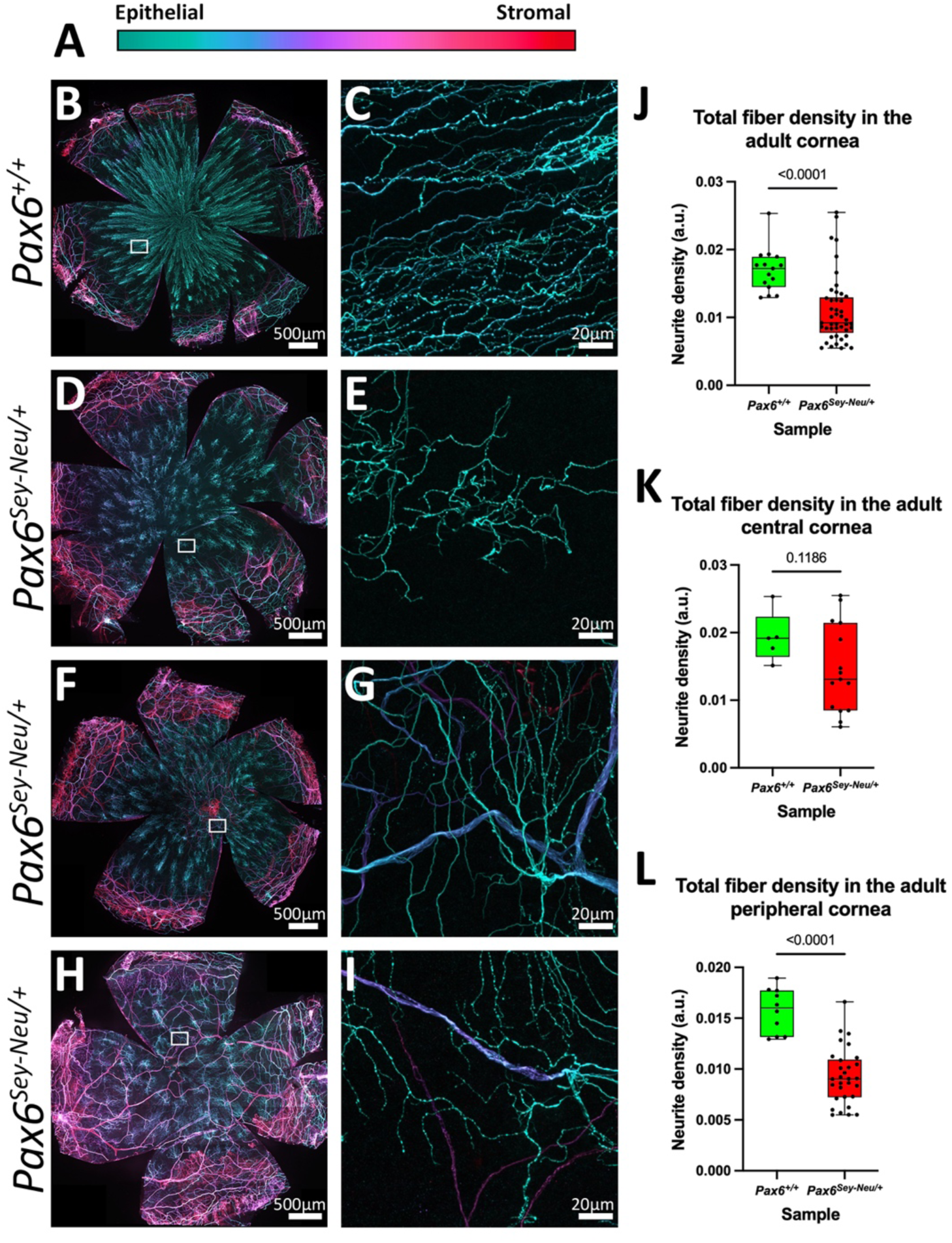
Adult *Pax6^Sey-Neu^*^/+^ corneas exhibit abnormal nerve architecture and reduced nerve density. (A) Depth-encoded color scale applied to all images in B–I. TuJ1-labeled fibers are pseudocolored by Z position within the confocal stack, from superficial/epithelial (green) to deep/stromal (red). (B, C) Adult *Pax6*^+/+^ cornea. The whole mount (B) shows the organized subbasal nerve plexus converging into a central whorl, whereas the higher-magnification field (C) shows densely packed and uniformly oriented subbasal fibers. (D–I) Three individual adult *Pax6^Sey-Neu^*^/+^ corneas illustrating the range of nerve phenotypes observed. In each case, the whorl is absent or poorly formed, and the subbasal plexus is sparse and irregularly organized. Large-caliber fibers occupying deeper stromal planes (blue-magenta) remain prominent in regions where subbasal fibers are reduced (G, I). (B, D, F, H) Whole-mount views of the entire cornea. White boxes indicate the location of the corresponding higher-magnification field. (C, E, G, I) Higher-magnification views of the boxed regions. All panels are maximum-intensity projections of confocal Z-stacks spanning the full corneal thickness. Scale bars: 500 μm (B, D, F, H) and 20 μm (C, E, G, I). Mouse ages are given in the Ages table (Supplementary Methods). Images are representative of the group; see Results for n values. (J–L) Quantification of total TuJ1^+^ nerve fiber density across the entire cornea (J), within a central ROI positioned inside the central 50% of corneal area (K), and within two equal-area peripheral ROIs positioned outside the central zone (L). Peripheral ROIs were permitted up to 20% overlap with the central zone to avoid inclusion of limbal fibers. Fiber density was quantified as thresholded particle counts normalized to ROI area and is expressed in arbitrary units (a.u.; see Methods). Each point represents one ROI measurement (three non-overlapping ROIs per cornea: one central and two peripheral; see Methods and Supplementary Table 1). Boxes indicate the median and interquartile range; whiskers indicate the minimum and maximum values. Groups were compared using two-tailed Mann-Whitney tests. Exact P values are shown. Density was reduced in *Pax6^Sey-Neu^*^/+^ corneas across the whole cornea (J) and within peripheral ROIs (L), whereas central ROI density did not differ significantly between genotypes (K).

### Corneal transparency is associated with region-specific nerve patterning in *Pax6* haploinsufficient corneas

Given the heterogeneity of adult *Pax6^Sey-Neu^*^/+^ corneas, we asked whether differences in nerve architecture were associated with corneal transparency. Corneas were classified as transparent (*Pax6^Sey-Neu^*^/+^-T), mixed (*Pax6^Sey-Neu^*^/+^-M), or opaque (*Pax6^Sey-Neu^*^/+^-O) based on gross appearance and examined by whole-mount TuJ1 immunolabeling (Figure 3).

**Figure 3.**
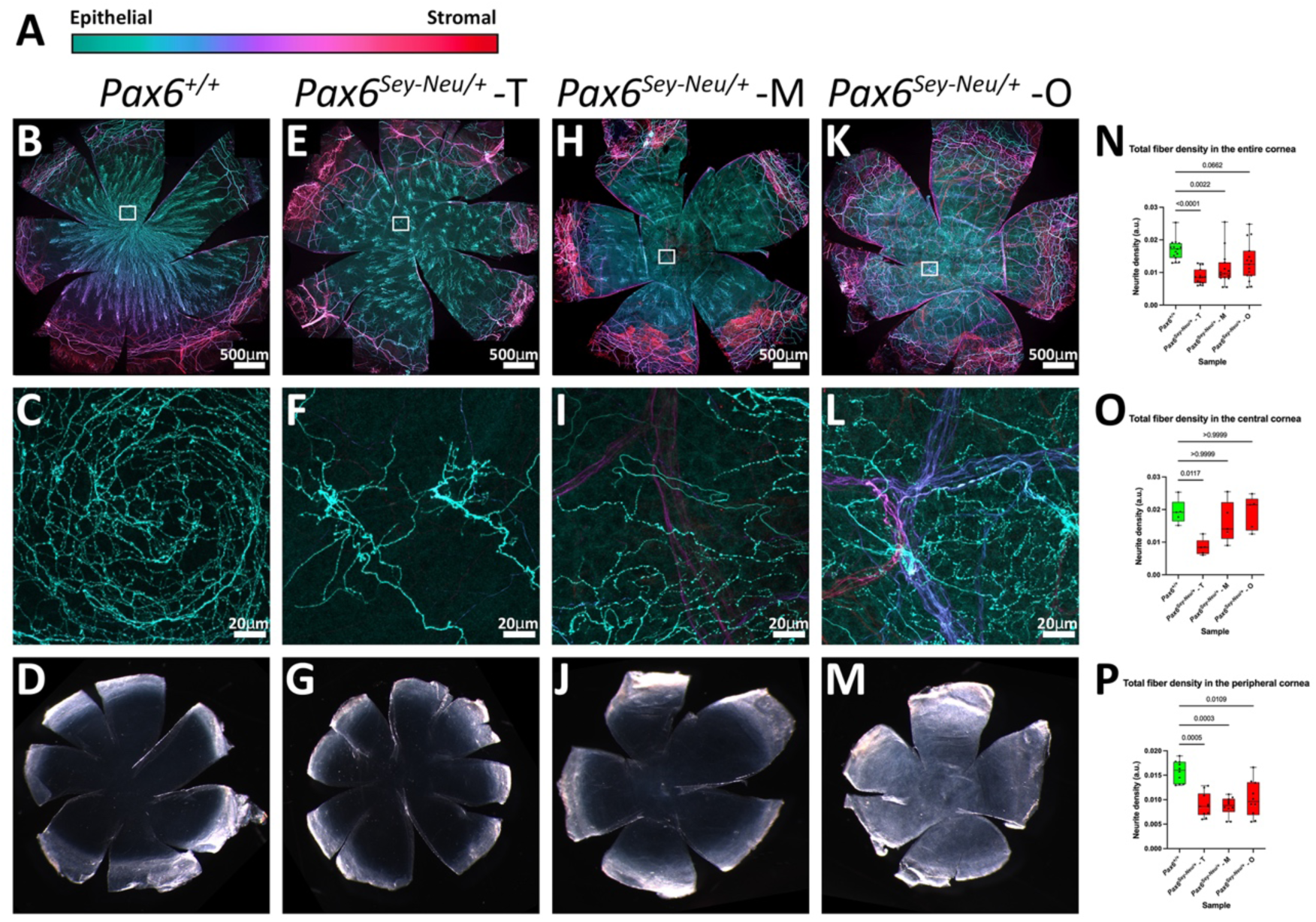
Corneal transparency is associated with region-specific nerve patterning in adult *Pax6^Sey-Neu^*^/+^ corneas. (A) Depth-encoded color scale applied to all fluorescence images. TuJ1-labeled fibers are pseudocolored by Z position within the confocal stack, from superficial/epithelial (green) to deep/stromal (red). Columns show *Pax6*^+/+^ (B–D) and *Pax6^Sey-Neu^*^/+^ corneas classified by transparency phenotype as transparent (–T; E–G), mixed (–M; H–J), or opaque (–O; K–M). These categories represent distinct presentations and are not intended as a severity series. Corneas were classified from images acquired under incident illumination against a black background following immunolabeling and confocal imaging (see Methods). (B, E, H, K) Whole-mount corneas immunolabeled for TuJ1. White boxes indicate the location of the corresponding higher-magnification field. (C, F, I, L) Higher-magnification views of the boxed regions, obtained from the central cornea in each case. *Pax6*^+/+^ corneas (C) contain densely packed superficial subbasal fibers. In many *Pax6^Sey-Neu^*^/+^ corneas, subbasal fibers are sparse and irregularly organized (F, I, L). Corneas containing opaque regions additionally exhibit large-caliber fibers occupying deeper stromal planes within the central cornea (I, L). (D, G, J, M) Backscatter images of the same corneas shown in B, E, H, and K. Transparent corneas appear dark centrally, whereas opaque regions appear brighter and more reflective. Mixed corneas contain discrete opaque regions within an otherwise transparent cornea (J), whereas opaque corneas exhibit opacity across most of the corneal surface (M). All fluorescence panels are maximum-intensity projections of confocal Z-stacks spanning the full corneal thickness. Scale bars: 500 μm (B, E, H, K) and 20 μm (C, F, I, L). Panels are representative of the group; see Results for n values. Corneas shown are distinct from those illustrated in Figure 2. (N–P) Quantification of total TuJ1^+^ nerve fiber density across the entire cornea (N), within a central ROI positioned inside the central 50% of corneal area (O), and within two equal-area peripheral ROIs positioned outside the central zone (P). Peripheral ROIs were permitted up to 20% overlap with the central zone to avoid inclusion of limbal fibers. Fiber density was quantified as thresholded particle counts normalized to ROI area and is expressed in arbitrary units (a.u.; see Methods). Each point represents one ROI measurement (three non-overlapping ROIs per cornea: one central and two peripheral; see Methods and Supplementary Table 1). Boxes indicate the median and interquartile range; whiskers indicate the minimum and maximum values. Groups were compared using Kruskal-Wallis tests followed by Dunn’s multiple-comparisons procedure. Adjusted P values for each *Pax6^Sey-Neu^*^/+^ group versus *Pax6*^+/+^ are shown. Total density was reduced in transparent and mixed mutant corneas (N). Peripheral density was reduced in all mutant groups (P), whereas central density was reduced only in transparent mutant corneas (O).

*Pax6*^+/+^ corneas displayed a dense subbasal plexus and characteristic central whorl (Figure 3B,C), together with a uniformly transparent surface (Figure 3D). Transparent mutant corneas contained sparse clusters of subbasal fibers but lacked central stromal fascicles and appeared grossly clear (Figure 3E-G). In contrast, mixed and opaque mutant corneas exhibited stromal fascicles extending into central corneal regions (Figure 3H-L), corresponding to localized or widespread loss of transparency (Figure 3J, M). These observations reveal a spatial association between reduced transparency and the persistence of stromal nerve fibers within the cornea. Quantification demonstrated that total nerve density was reduced in transparent and mixed mutant corneas relative to wild-type controls (Figure 3N). Analysis of regional density revealed that peripheral nerve density was reduced in all three mutant groups (Figure 3P), whereas a significant reduction in central nerve density was detected only in transparent mutant corneas (Figure 3O). Independent analyses of epithelial compartments are shown in Supplementary Figure 1I-K. Together, these findings indicate that reduced nerve density is a general feature of the *Pax6^Sey-Neu^*^/+^ cornea, whereas differences in transparency are associated primarily with distinct patterns of nerve organization, particularly the persistence of stromal fascicles within affected regions. Because stromal fibers are normally unmyelinated within the cornea, we next asked whether the altered nerve architecture in mutant corneas was accompanied by changes in myelination status.

### Segmented stromal myelination is detected in opaque Pax6 haploinsufficient corneas

We next asked whether altered nerve architecture in *Pax6^Sey-Neu^*^/+^ corneas was accompanied by changes in myelination. Adult corneas were co-labeled for TuJ1 and myelin basic protein (MBP) and classified according to transparency state (Figure 4). In *Pax6*^+/+^ corneas, MBP labeling was largely restricted to limbal fibers and was absent from the central cornea (Figure 4D,E). A similar pattern was observed in transparent *Pax6^Sey-Neu^*^/+^ corneas, in which MBP labeling was confined primarily to limbal and peripheral fibers (Figure 4H,I). In contrast, MBP-positive stromal fibers were detected in all corneas containing opaque regions, whether focal or widespread (Figure 4L-Q). These MBP-positive fibers were restricted to affected regions and displayed discontinuous labeling along individual nerve bundles. To determine whether MBP immunoreactivity reflected compact myelination, stromal nerve fibers were examined by transmission electron microscopy (Figure 4S-Z, Supplementary Table 2). Stromal axons in *Pax6*^+/+^ and transparent *Pax6^Sey-Neu^*^/+^ corneas were organized as unmyelinated Remak-type bundles (Figure 4S-V). In contrast, compact multilamellar myelin was observed in nerve fibers associated with opaque regions of mutant corneas (Figure 4X-Z), although unmyelinated fibers remained present within the same tissue. Together, these findings demonstrate that stromal myelination is not a general consequence of Pax6 haploinsufficiency but is specifically associated with corneal regions that have lost transparency.

**Figure 4.**
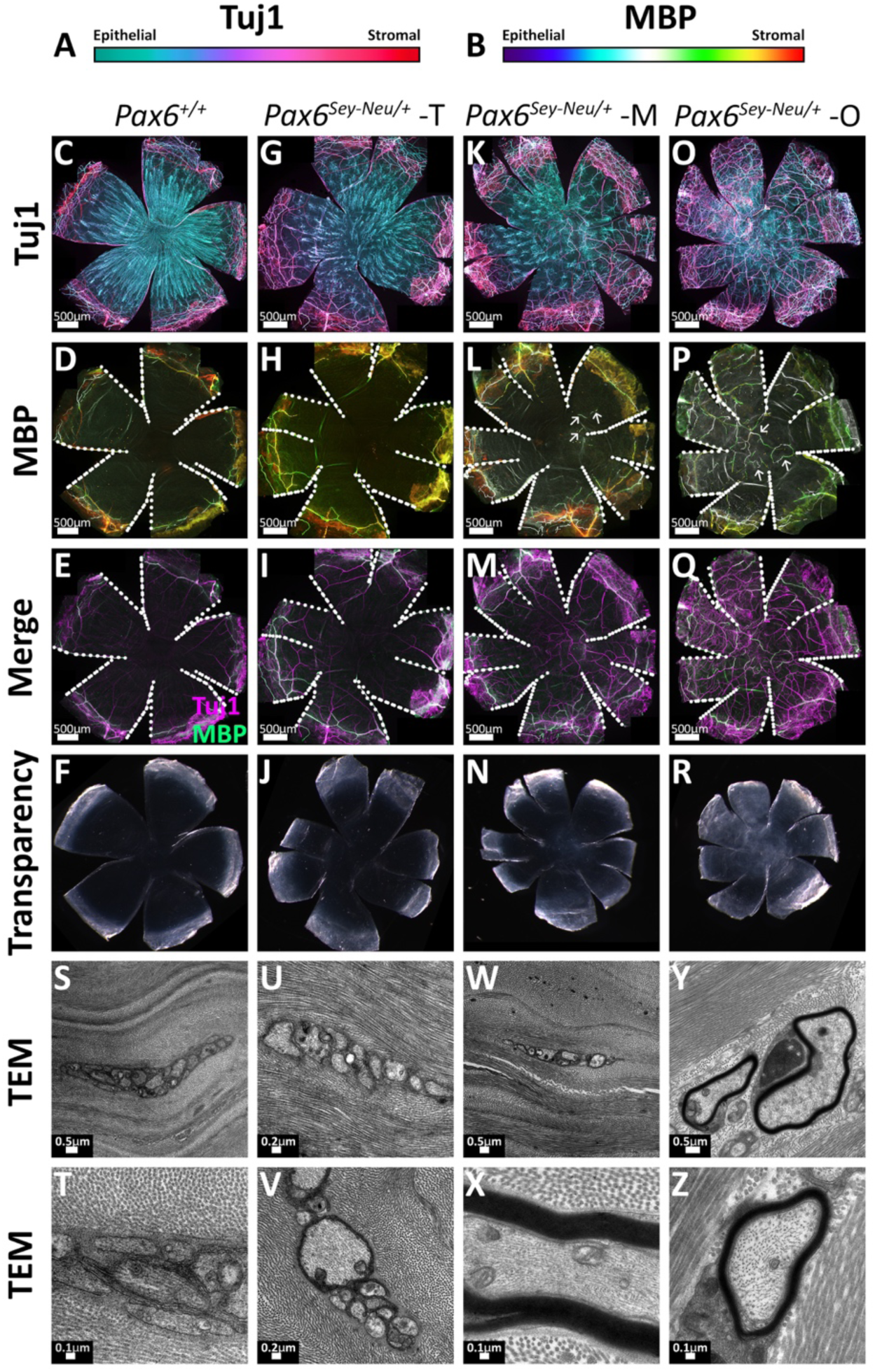
Compact stromal myelination is associated with opaque regions in *Pax6^Sey-Neu^*^/+^ corneas. (A) Depth-encoded color scale for TuJ1-labeled fibers, from superficial/epithelial (green) to deep/stromal (red). (B) Depth-encoded color scale for MBP immunolabeling, using the same depth convention. Columns show *Pax6*^+/+^ (C–F, S–T) and *Pax6^Sey-Neu^*^/+^ corneas classified as transparent (–T; G–J, U–V), mixed (–M; K–N, W–X), and opaque (–O; O–R, Y–Z). (C, G, K, O) Maximum-intensity projections of TuJ1-labeled nerve fibers across the entire cornea. (D, H, L, P) Maximum-intensity projections of selected optical sections containing MBP immunolabeling (see Methods). MBP labeling was not detected in the central cornea of *Pax6*^+/+^ (D) or transparent *Pax6^Sey-Neu^*^/+^ corneas (H). MBP-positive fibers are present in mixed and opaque mutant corneas (L, P; arrows). White dotted lines indicate the relaxing incisions used to flatten the cornea. (E, I, M, Q) Merged images showing TuJ1 (magenta) and MBP (green) over the same Z range. Channels are displayed in conventional two-color format rather than depth-encoded format. (F, J, N, R) Backscatter images of the same corneas, illustrating transparency phenotypes as defined in Figure 3. Relative scales apply. All fluorescence panels are maximum-intensity projections of confocal Z-stacks. Mouse ages are given in the Ages table (Supplementary Methods). Panels are representative of the group; see Results for n values. Corneas shown are distinct from those illustrated in Figures 2 and 3. (S–Z) Transmission electron micrographs of stromal nerve profiles from *Pax6*^+/+^ (S, T), transparent *Pax6^Sey-Neu^*^/+^ (U, V), mixed *Pax6^Sey-Neu^*^/+^ (W, X), and opaque *Pax6^Sey-Neu^*^/+^ (Y, Z) corneas. Stromal axons in *Pax6*^+/+^, transparent mutant, and transparent regions of mixed corneas are organized as unmyelinated Remak-type bundles ensheathed by Schwann cell cytoplasm (S–W). Compact multilamellar myelin is present in stromal nerve fibers sampled from opaque regions of mixed corneas (X) and opaque corneas (Y, Z), although unmyelinated fibers remain present within the same tissues. Electron microscopy was performed on separate animals from those used for immunolabeling. Panels S, T, U, V, and Y were obtained from the central cornea; panel Z was obtained from the peripheral cornea. Scale bars as indicated.

### MBP labeling in mutant corneas is spatially restricted and discontinuous

Higher-magnification imaging was used to examine the organization of MBP labeling within individual nerve fibers (Figure 5). In both *Pax6*^+/+^ and *Pax6^Sey-Neu^*^/+^ corneas, MBP labeling was detected on a subset of limbal fibers (Figure 5A,C). In the peripheral cornea of *Pax6*^+/+^, TuJ1-positive fibers lacked detectable MBP labeling (Figure 5B). In contrast, mutant corneas contained MBP-positive stromal fibers interspersed with MBP-negative fibers within the same field (Figure 5D-F). MBP labeling was discontinuous and restricted to subsets of nerve fascicles and individual fibers rather than uniformly distributed throughout the nerve population. XZ reslices confirmed that MBP-positive fibers were confined to the stromal compartment and were not detected within epithelial or subbasal nerve layers (Figure 5G,H). Across all corneas examined, MBP immunoreactivity was observed only in stromal fibers and only in mutant corneas containing opaque regions. Together, these observations indicate that the myelination-associated phenotype in *Pax6^Sey-Neu^*^/+^ corneas is spatially restricted and heterogeneous, involving only a subset of stromal nerve fibers rather than the corneal nerve population as a whole.

**Figure 5.**
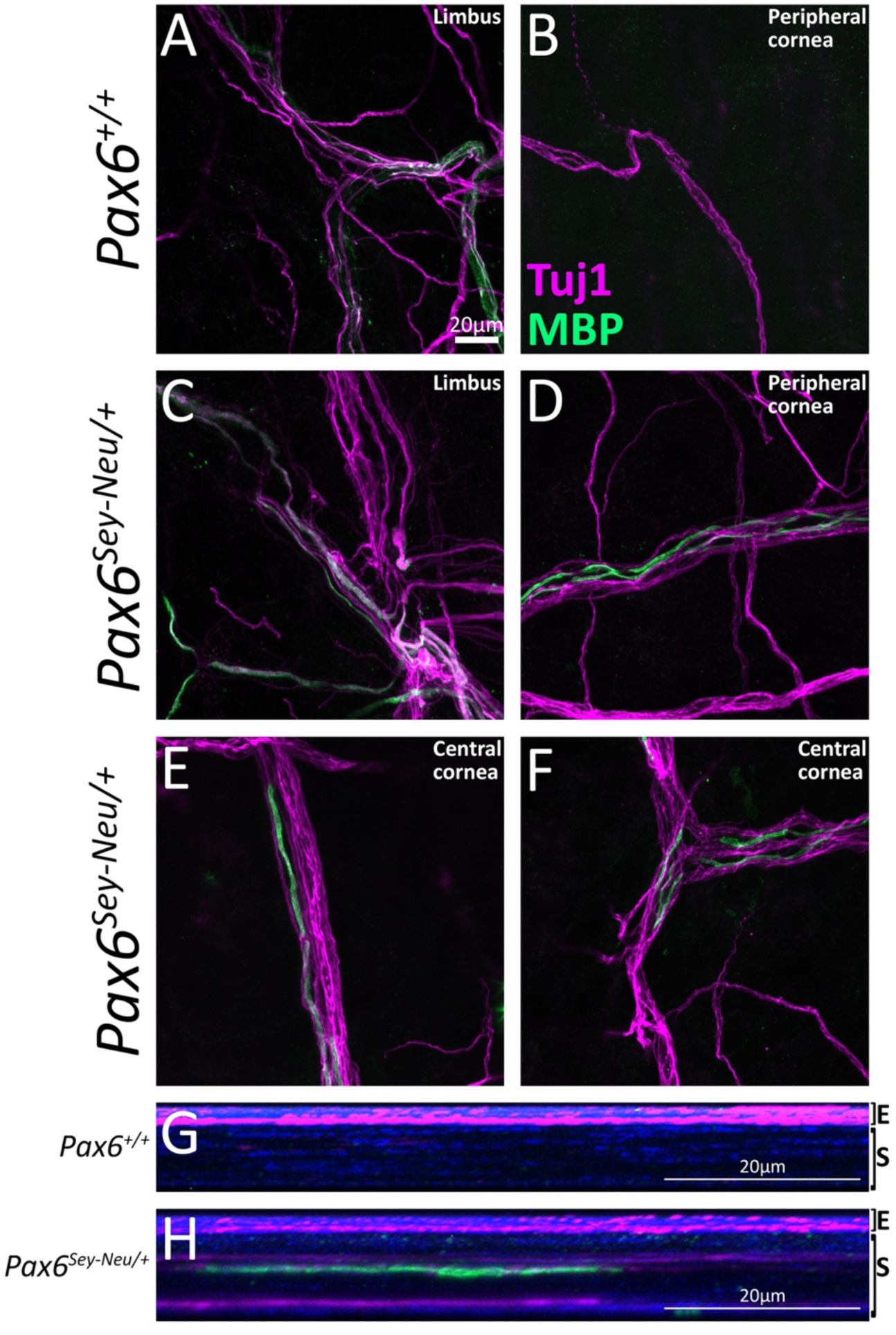
MBP labeling is spatially restricted and discontinuous in *Pax6^Sey-Neu/+^* corneas. (A–F) TuJ1 (magenta) and MBP (green) immunolabeling; regions of overlap appear white. (A) Limbal nerve fibers in *Pax6*^+/+^ cornea. MBP labeling is present on a subset of fibers, consistent with normal myelination of limbal nerves. (B) Peripheral corneal nerve fiber in *Pax6*^+/+^ cornea. TuJ1-positive fibers lack detectable MBP labeling. (C) Limbal nerve fibers in *Pax6^Sey-Neu^*^/+^ cornea. As in *Pax6*^+/+^, MBP labeling is restricted to a subset of limbal nerve fibers. (D) Peripheral corneal nerve fibers in a *Pax6^Sey-Neu^*^/+^ cornea. (E, F) Central corneal nerve fibers from the same *Pax6^Sey-Neu^*^/+^ cornea shown in D. MBP labeling is present on only a subset of TuJ1-positive fibers and appears discontinuous along individual nerve fibers. Both MBP-positive and MBP-negative TuJ1-positive fibers are present within the same field. No MBP labeling was detected in the peripheral cornea of *Pax6*^+/+^ mice (B). All panels are maximum-intensity projections of selected optical sections (1 μm per section) spanning the nerve fibers of interest. Scale bar in A applies to A–F (20 μm). (G, H) XZ reslices corresponding to the fields shown in B and D, respectively. DAPI, blue; TuJ1, magenta; MBP, green. E, epithelium; S, stroma. TuJ1-positive fibers occupy epithelial/subbasal layers in both genotypes. An MBP-positive stromal fiber is present in the mutant cornea (H) and absent from the *Pax6*^+/+^ cornea (G). Scale bars, 20 μm.

### Corneal neovascularization in Pax6 haploinsufficient corneas

To determine whether vascular abnormalities accompany changes in nerve organization and corneal transparency, corneas were immunolabeled for the endothelial marker CD31 and examined by confocal microscopy (Figure 6). In *Pax6*^+/+^ corneas, CD31-positive structures were largely restricted to the limbal region, with little or no labeling detected within the cornea proper (Figure 6B,F). In contrast, *Pax6^Sey-Neu^*^/+^ corneas displayed variable patterns of intracorneal CD31 labeling (Figure 6C-H). A minority of mutant corneas retained a limbus-restricted pattern resembling wild type (4 of 23 corneas), whereas others exhibited CD31-positive structures extending into peripheral and central corneal regions (Figure 6D,E). XZ projections confirmed the presence of CD31-positive structures within the anterior corneal stroma of affected mutant corneas (Figure 6H). To determine when these differences first emerge, CD31 labeling was examined across postnatal development (Supplementary Table 3). Peripheral labeling was present at birth in both genotypes (3 of 3 wild-type and 2 of 3 mutant corneas). In wild-type corneas, this labeling was subsequently largely lost and was not detected in the central cornea at any age examined. In contrast, mutant corneas showed persistent peripheral labeling and extension into central corneal regions by 2 weeks of age. Thus, vascular abnormalities emerge during the first two postnatal weeks, at ages when all corneas remained transparent.

**Figure 6.**
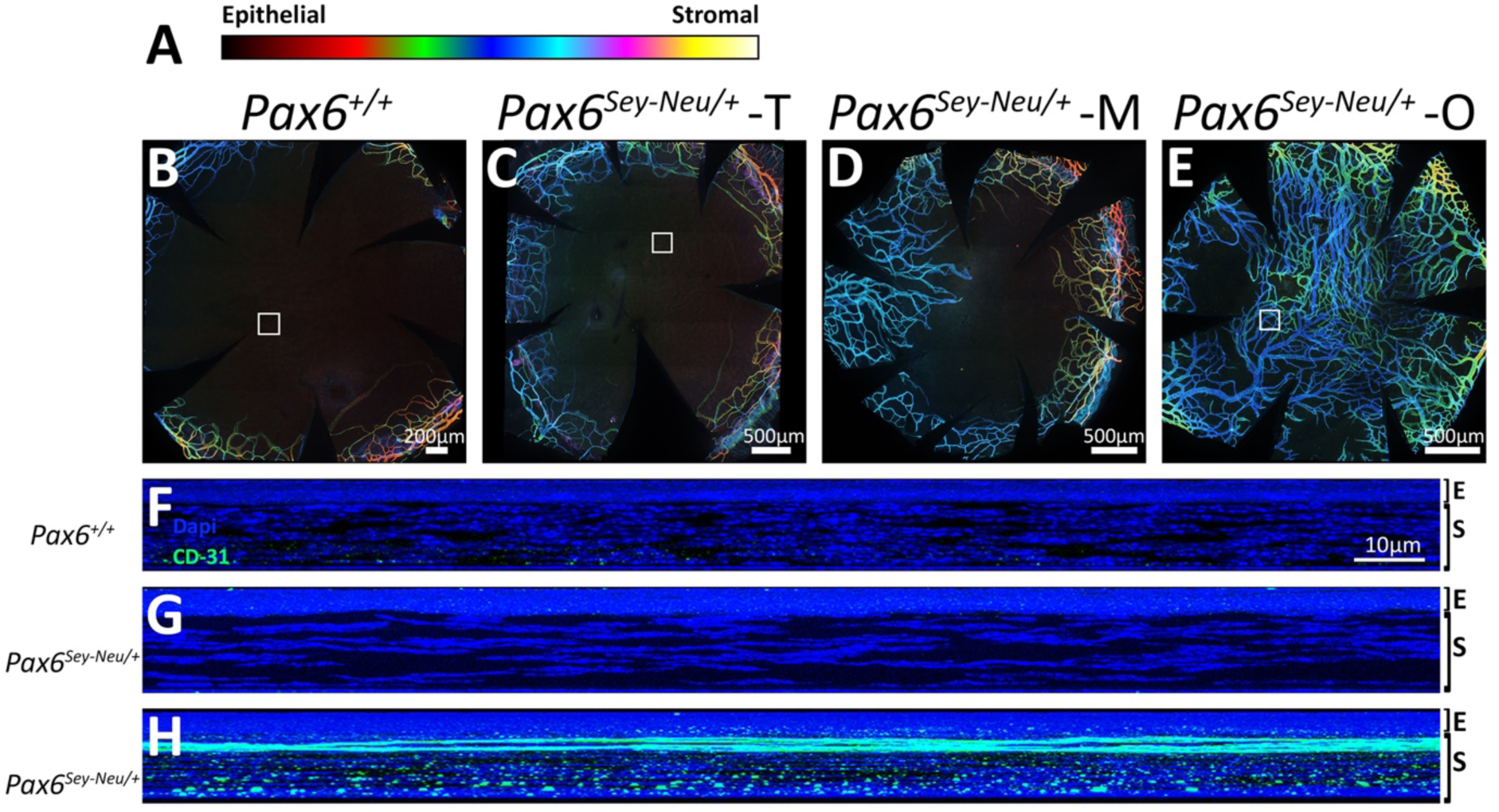
Vascular structures persist and extend into the cornea of *Pax6^Sey-Neu/+^* mice. (A) Depth-encoded color scale for CD31 immunolabeling, from superficial/epithelial (green) to deep/stromal (red). (B–E) Maximum-intensity projections of corneal flat mounts immunolabeled for CD31, showing *Pax6*^+/+^ (B) and *Pax6^Sey-Neu^*^/+^ corneas classified as transparent (–T; C), mixed (–M; D), and opaque (–O; E). In *Pax6*^+/+^ corneas, CD31 labeling is restricted to the limbal region and is not detected within the cornea proper (B). *Pax6^Sey-Neu^*^/+^ corneas exhibit variable patterns of intracorneal CD31 labeling, ranging from limbus-restricted labeling resembling wild type to CD31-positive structures extending into peripheral and central corneal regions (C–E). The cornea shown in C represents one of the minority of mutant corneas (4 of 23) that retained a limbus-restricted pattern. Scale bars: 200 μm (B) and 500 μm (C–E). Mouse ages are given in the Ages table (Supplementary Methods). Panels are representative of the group; see Results for n values. (F–H) XZ reslices showing the depth distribution of CD31 labeling in *Pax6*^+/+^ (F) and *Pax6^Sey-Neu^*^/+^ corneas (G, H). Panel G corresponds to the cornea shown in C, in which CD31 labeling remained limbus restricted and was not detected within the anterior corneal stroma. Panel H corresponds to the cornea shown in E and demonstrates CD31-positive structures within anterior stromal layers. DAPI, blue; CD31, green. E, epithelium; S, stroma. Scale bar in F applies to G and H. CD31 labels vascular endothelium but does not distinguish blood vessels from lymphatic vessels. Developmental frequencies of intracorneal CD31-positive structures are summarized in Supplementary Table 3.

Together, these findings demonstrate that reduced Pax6 dosage is associated with early and persistent vascular abnormalities in the cornea and that vascular ingrowth precedes the loss of transparency.

### Postnatal maturation of the wild-type cornea silences a largely vascular and lymphatic program, and the mutant cornea does not

Across postnatal development, comparing ages within genotype, 144 expression-status crossings were detected among 58 panel genes (Figure 7, Supplementary Tables 5 and 6). Development was predominantly a silencing process: 104 crossings represented genes turning off with age, whereas 40 represented genes turning on. Of the 144 crossings, 126 were also significant in the same direction when each interval was tested without the status gate by replicate-aware differential expression (Supplementary Figure 4; Supplementary Table 7).

**Figure 7.**
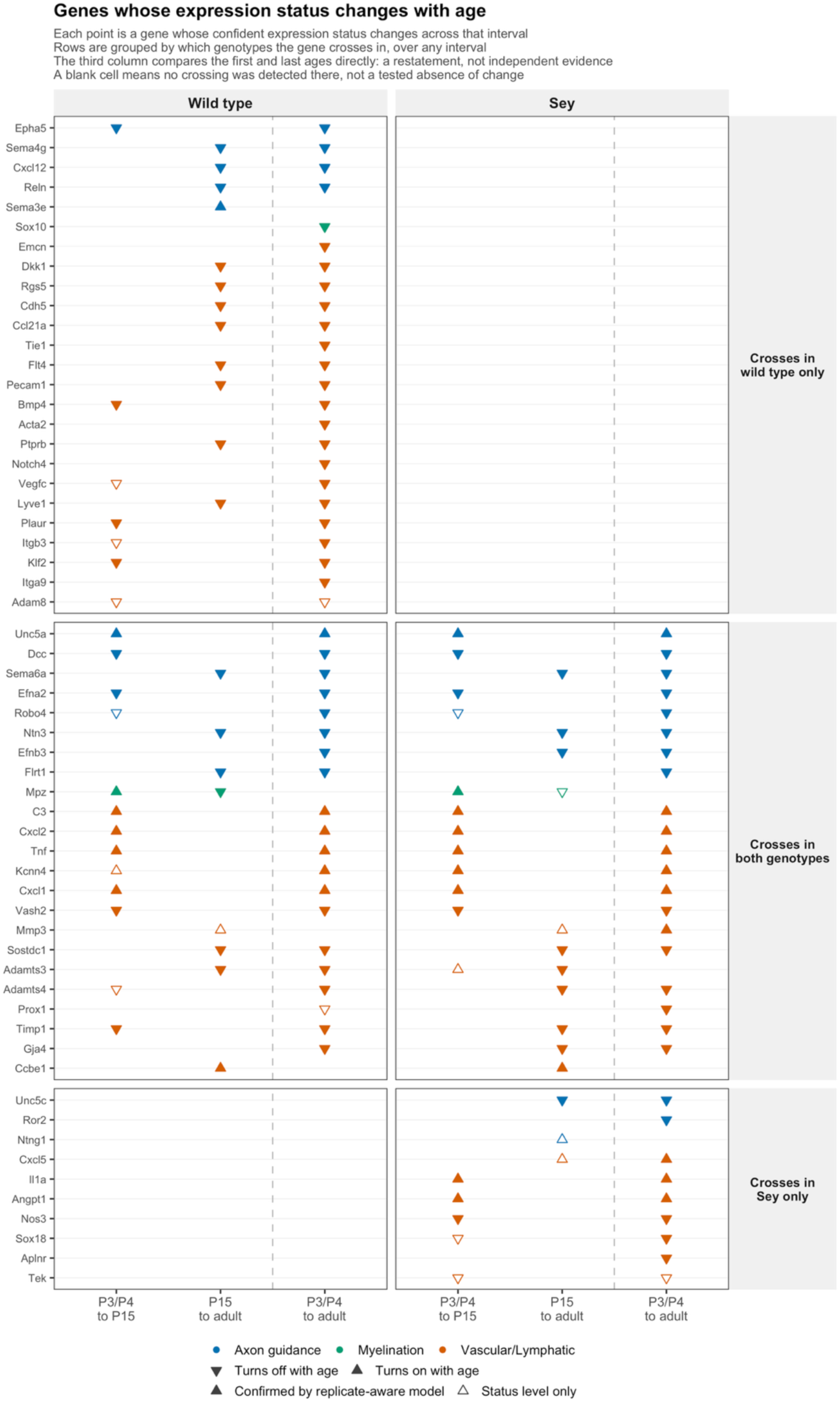
Postnatal maturation silences a vascular and lymphatic program in wild-type but not *Pax6^Sey-Neu^*^/+^. Expression-status crossings across corneal development in *Pax6*^+/+^ and *Pax6^Sey-Neu^*^/+^ (abbreviated *Sey*) corneas, based on RNA-seq of whole cornea collected at P3/P4, P15, and adulthood. Wild-type corneas: n = 4, 5, and 5 libraries; *Pax6^Sey-Neu^*^/+^ corneas: n = 5, 5, and 5 libraries. Adult mutant samples were restricted to transparent corneas so that developmental comparisons were not confounded by loss of transparency. Filled symbols denote crossings also confirmed by replicate-aware differential expression analysis (adjusted P < 0.05, same direction; Methods); 126 of 144 crossings are confirmed, and the 18 that are not are drawn open. A gene is plotted when it changes categorical expression status between ages, transitioning between confidently expressed and confidently silent states, or when the corresponding exact Poisson confidence intervals are nonoverlapping and differ by at least twofold. Crossings therefore represent changes in expression status rather than differential-expression model results. Downward triangles indicate genes turning off with age, whereas upward triangles indicate genes turning on. Colors denote the primary curated panel assignment used for multiple-testing correction; genes assigned to more than one panel are shown once under their primary panel. Rows are grouped according to the genotype(s) in which a gene crosses at any developmental interval. Twenty-five genes cross only in wild type (19 vascular/lymphatic, 5 axon-guidance, and 1 myelination gene), 23 cross in both genotypes (14, 8, and 1 genes, respectively), and 10 cross only in *Pax6^Sey-Neu^*^/+^ corneas (7 vascular/lymphatic and 3 axon-guidance genes). Across the full dataset, 144 crossings occur in 58 genes, with 104 representing genes turning off during maturation and 40 representing genes turning on. The first two columns show crossings occurring between adjacent developmental intervals. The third column summarizes the full developmental interval and is not an independent test. A blank cell indicates that no status crossing was detected and should not be interpreted as evidence that a difference was excluded. Whether genotype effects change with age was tested separately using genotype-by-age interaction models (Supplementary Table 8).

In wild-type cornea, these crossings occurred in two distinct phases. Between P3/P4 and P15, genes involved in guidance signaling, matrix regulation, and growth-factor pathways were preferentially silenced, including *Dcc* and *Epha5* and, at the status level only, *Robo4, Adamts4,* and *Vegfc*. Between P15 and adulthood, a second group of genes associated with endothelial and lymphatic identity was silenced, including *Pecam1, Cdh5, Flt4, Ptprb, Lyve1, Ccl21a,* and *Rgs5*. Thus, normal corneal maturation includes progressive withdrawal of a vascular and lymphatic transcriptional program.

The two genotypes differed primarily in execution of this second phase. Of the 25 genes that crossed in wild type only, 19 belonged to the vascular/lymphatic panel, compared with 5 axon-guidance genes and 1 myelination gene. Many endothelial and lymphatic markers that were silenced during wild-type maturation, including *Pecam1, Cdh5, Tie1, Flt4, Ptprb, Lyve1, Ccl21a,* and *Rgs5*, did not change status in the mutant cornea at any developmental interval. In contrast, genes crossing in both genotypes were distributed more evenly across the three panels. The myelination gene *Mpz* displayed a transient developmental peak in wild type, increasing between P3/P4 and P15 in both genotypes before declining by adulthood in wild type; the decline in the mutant is not confirmed by the replicate-aware model. These findings indicate that postnatal maturation of the mutant cornea specifically fails to execute much of the normal vascular and lymphatic silencing program.

Several developmental changes occurred similarly in both genotypes. The attractive netrin receptor *Dcc* declined with age whereas the repulsive receptor *Unc5a* increased, indicating that elements of developmental guidance-cue remodeling remain intact despite reduced Pax6 dosage. In contrast, *Unc5c* declined in both genotypes but fell below confident detection only in mutant corneas, producing a categorical adult genotype difference.

Two limitations should be noted. First, a blank cell in Figure 7 indicates that no status crossing was detected, not that a difference was excluded. Second, the developmental interval in which a crossing occurs is not itself compared between genotypes. Direct testing of genotype-by-age interactions detected significant effects for only 16 genes, whereas more than 93% of interaction tests remained unresolved (Supplementary Table 8). Accordingly, the Results below focus on comparisons that reached significance rather than on inferred times of onset. The assignment of genes to wild-type-only, shared and Sey-only groups in Figure 7 is specific to the configured expression thresholds and was not stable when those thresholds were varied (Supplementary Table 9).

### The vascular and lymphatic program is already expanded in the transparent mutant cornea and expands further with loss of transparency

To determine how reduced Pax6 dosage and loss of transparency affect corneal gene expression, every panel gene was classified as confidently expressed, confidently silent, or indeterminate in each comparison (Supplementary Table 10). Across the four comparisons, 831 panel-gene entries were confidently expressed in both groups and could be tested quantitatively, 50 differed categorically (45 distinct gene-by-comparison changes, because some genes belong to more than one panel) in expression status, 67 were confidently silent in both groups, and the remainder were unresolved because expression status could not be determined.

At the level of whole programs, the vascular/lymphatic panel showed the strongest and most consistent differences (Figure 8A). In transparent adult *Pax6^Sey-Neu^*^/+^ corneas, the vascular/lymphatic panel contained significantly more confidently expressed genes and fewer confidently silent genes than abundance-matched controls. A similar excess was detected at P15 and increased further in opaque relative to transparent mutant corneas. In contrast, the axon-guidance panel reached significance only in the opacity comparison, whereas the myelination panel showed no robust program-level differences. These findings indicate that vascular and lymphatic gene expression is already expanded in the transparent mutant cornea and undergoes further activation with loss of transparency.

**Figure 8.**
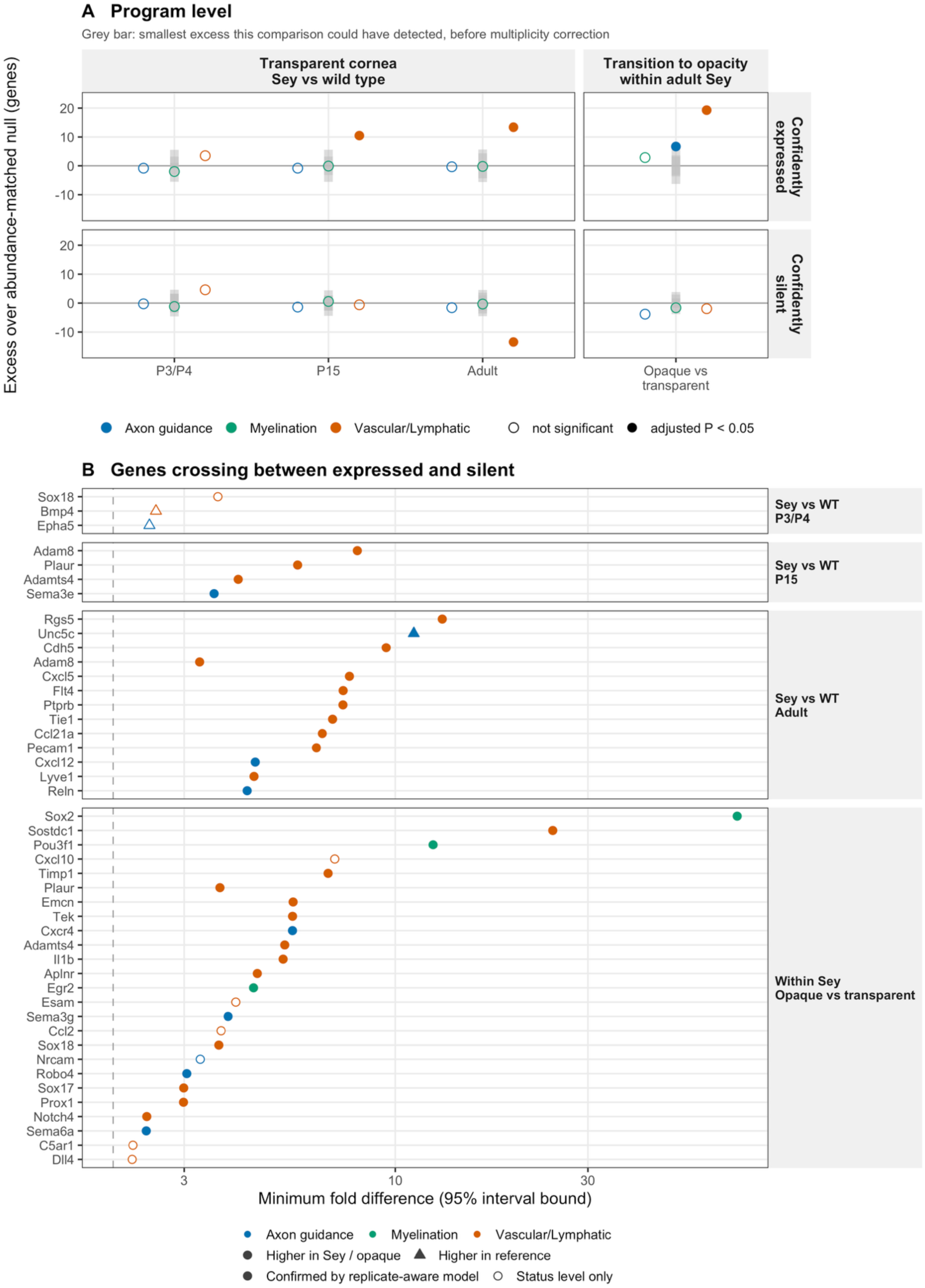
The vascular and lymphatic program is expanded before opacity and increases further with loss of transparency. Both panels separate two biological axes throughout. Left: *Pax6^Sey-Neu^*^/+^ versus *Pax6*^+/+^ at P3/P4, P15, and adulthood, with all corneas transparent at collection, isolating genotype effects independent of opacity. Right: opaque versus transparent adult *Pax6^Sey-Neu^*^/+^ corneas, isolating changes associated with loss of transparency within genotype. The comparison between opaque mutant and wild-type cornea combines both axes and is therefore excluded from this figure (shown in Supplementary Figure 7). (A) Program-level analysis of the curated axon-guidance, myelination, and vascular/lymphatic panels. Values represent the excess of confidently expressed or confidently silent genes relative to abundance-matched null gene sets drawn from non-panel genes. Twenty-four panel-by-comparison tests are shown; five remain significant following Benjamini-Hochberg correction across all tests. Grey bars indicate the unadjusted 95% interval of the matched null distribution for each comparison. Filled points denote significant results after correction. Significant expansion of the vascular/lymphatic program is detected at P15, in transparent adult mutant cornea, and in opaque relative to transparent mutant cornea. Axon-guidance reaches significance only in the opacity comparison, whereas the myelination panel shows no significant program-level expansion. (B) Individual genes exhibiting categorical expression-status differences between groups. Forty-five gene-by-comparison status changes are detected in 41 genes, comprising 20 changes in the genotype block and 25 changes in the opacity block. The x axis indicates the minimum fold difference compatible with the non-overlapping expression intervals; values are therefore lower bounds rather than point estimates. The dashed line indicates the twofold criterion used for categorical calls. Circles denote genes higher in *Pax6^Sey-Neu^*^/+^ or opaque cornea, whereas triangles denote genes higher in the reference group. Most categorical differences are increases in the test group, with 42 of 45 categorical changes occurring in mutant or opaque corneas. Filled symbols denote categorical differences also confirmed by replicate-aware differential expression analysis (adjusted P < 0.05, same direction; Methods); 36 of 45 are confirmed, and the nine that are not are drawn open.

Individual genes changing expression status are shown in Figure 8B (Supplementary Table 11). Between transparent adult mutant and wild-type corneas, 13 genes differed categorically, including multiple endothelial and lymphatic markers that are normally silenced during maturation (*Rgs5, Cdh5, Flt4, Ptprb, Tie1, Ccl21a, Pecam1,* and *Lyve1*). Thus, the abnormal developmental pattern identified in Figure 7 is retained in the adult transparent mutant cornea. Additional differences included *Cxcl5, Cxcl12, Reln, Adam8,* and *Unc5c*, the latter remaining higher in wild type.

Loss of transparency was associated with a second wave of transcriptional change. Opaque mutant corneas gained expression of additional endothelial and lymphatic genes, including *Emcn, Sox17, Sox18, Tek, Prox1, Notch4,* and *Aplnr* (and, at the status level only, *Esam* and *Dll4*), together with inflammatory and matrix-remodeling genes such as *Il1b, Adamts4,* and *Timp1* (and, at the status level only, *Ccl2* and *Cxcl10*). Notably, the only categorical changes involving myelination genes occurred in opacity-associated comparisons, where *Sox2, Pou3f1,* and *Egr2* became more highly expressed.

The mutant cornea also differed from wild type before adulthood. At P15, *Adam8, Plaur, Adamts4,* and *Sema3e* were confidently expressed in mutant but not confidently expressed in wild-type corneas, whereas at P3/P4 only three categorical differences were called, all at the status level only: *Sox18* was higher in the mutant and *Bmp4* and *Epha5* were higher in wild type. Among genes confidently expressed in both groups, quantitative changes followed the same pattern (Supplementary Figures 5 and 6): 99 gene-by-panel entries changed significantly in level in at least one comparison, 49 only while the cornea was transparent, 18 both before and after the transition to opacity, and 32 only with the transition. Increases predominated in the vascular/lymphatic panel (67 of 88 significant changes) more than in the axon-guidance and myelination panels (34 of 57). Across the four comparisons, categorical differences were overwhelmingly directional: 42 of 45 were higher in mutant or opaque corneas. Together, these findings indicate that reduced Pax6 dosage is associated primarily with persistence and expansion of a vascular and lymphatic expression program, whereas the transition to opacity is accompanied by broader vascular, inflammatory, matrix-remodeling, and myelination-related changes. The comparison of opaque mutant with wild-type cornea differs in genotype and in transparency at once, cannot be attributed to either, and is shown only in Supplementary Figure 7.

### Trigeminal ganglia showed no genotype-associated differences while the cornea remained transparent

To determine whether the corneal changes were accompanied by transcriptional alterations in the sensory neurons that innervate the tissue, trigeminal ganglia from a subset of the same animals were analyzed with the same curated gene panels. Fourteen libraries were available (Supplementary Table 12), two per group: wild-type and transparent-mutant ganglia at P3/P4, P15 and adult, and opaque-mutant adults. Two libraries per group are too few for expression-status calls, so the ganglia were analyzed by replicate-aware differential expression alone (Methods). No panel gene differed between genotypes at P3/P4, P15 or in adults with transparent corneas (smallest adjusted P = 0.30, 0.11 and 0.071, respectively), and 537 of the 745 gene-by-comparison tests excluded a change larger than 1.5-fold; the remaining 208 were undetermined. Differences appeared only in comparisons involving opaque animals, all in the same direction and modest in size. Eleven panel genes were 1.1- to 1.7-fold higher in opaque-mutant than in wild-type ganglia: the myelination genes *Egr2*, *Mag*, *Erbb3*, *Zeb2* and *Mbp* and the axon-guidance genes *Flrt3*, *Reln*, *Slit2*, *Sema5a*, *Adgrl2* and *Sema3b*. Three were higher in opaque than in transparent mutants: *Egr2* (1.8-fold), *Timp3* and *Mbp* (1.2-fold each). Markers of central nervous tissue showed that carry- over at dissection varied between libraries but was not specific to any group (Supplementary Methods). A wild-type age model fitted to the same libraries detected 11,468 of 20,287 tested genes as differentially expressed, showing that the libraries and analysis detect transcriptional differences where they exist.

Several limitations qualify this result. The small sample size permits detection only of relatively large genotype effects, and cornea-projecting neurons represent only a small fraction of trigeminal cells analyzed in whole-ganglion preparations. We therefore interpret these data as a pilot study. Within the limits of the present design, they do not detect a genotype-associated alteration of these programs in the trigeminal ganglion and are consistent with a predominantly target-derived mechanism.

### Expression-status classifications reproduce in an independent corneal RNA-seq dataset carrying a distinct Pax6 allele

To determine whether the expression-status classifications were reproducible across independent datasets, the identical framework was applied to an adult corneal RNA-seq dataset (GSE183742) generated by Melinda Duncan’s laboratory from three wild-type and three *Pax6tm1Pgr/+* corneas. Before comparison, the implementation was required to reproduce the adult wild-type calls from the present study exactly. Among the 263 panel genes, 155 were resolved in both datasets and 152 of those received the same classification, corresponding to 98.1% agreement (κ = 0.79). The three disagreements all involved genes classified as silent in the present dataset and expressed in GSE183742, the pattern expected if the GSE183742 dissections included more peripheral tissue, which markers of limbus, angle, pigmented tissue and retina confirm (Supplementary Methods), rather than opposite biological conclusions.

Genotype-associated categorical differences were also highly concordant. Of 47 panel genes exhibiting categorical differences in both datasets, 46 showed genotype effects in the same direction (κ = 0.85). The only disagreement involved *Dscam*, which was higher in mutant corneas in the present study and lower in the independent dataset. Two qualifications apply. Transparency is not reported for GSE183742, so the direction comparison used the opaque-mutant versus wild-type contrast of the present study, in which genotype and opacity cannot be separated; and neither comparison exceeded the agreement of abundance-matched non-panel genes (wild-type status, 98.1% against a null median of 96.8%, P = 0.19; direction, 97.9% against 97.9%, P = 0.53), so the panel calls transfer as well as comparable genes do, not better. These findings demonstrate that the expression-status classifications are reproducible across independent datasets and distinct *Pax6* haploinsufficient alleles. Although the analysis cannot establish identical behavior of individual genes between alleles, it indicates that the principal conclusions are unlikely to be artifacts of a particular dataset, library-preparation strategy, or computational implementation.

## Discussion

### Which models of aniridia-associated keratopathy do these data support?

Several, not mutually exclusive, explanations have been proposed for AAK: limbal stem cell deficiency; intrinsic failure of corneal epithelial differentiation and adhesion; a chronic wound-healing and inflammatory state; loss of corneal angiogenic and lymphatic privilege through arrested postnatal development; a neurotrophic deficit; and anterior-segment dysgenesis, including keratolenticular adhesion (Collinson et al., 2004; Davis et al., 2003; Javidjam et al., 2024; Lagali et al., 2020; Latta et al., 2021; Mort et al., 2011; Ramaesh et al., 2003; Ramaesh et al., 2005). Our measurements (nerve imaging, CD31 labeling, TEM and limbus-excluded whole-cornea RNA sequencing of three curated panels) can test models that make claims about nerves, vessels and timing relative to opacity, but not those centered on epithelial identity or lens–cornea relationships.

Loss of angiogenic and lymphatic privilege through arrested postnatal development is the model best supported by our data. In *Pax6^+/−^* (Sey) mice, Javidjam et al. found lymphatic vessels in the central cornea by one month, outnumbering blood vessels three- to fourfold, persisting without regression and accompanied by macrophages absent from wild type, and proposed a “frozen” developmental state (Javidjam et al., 2024). We show that peripheral CD31-positive structures are present in both genotypes at birth and largely cleared from wild-type corneas, so the regression the mutant fails to complete is a normal part of maturation (Supplementary Table 3); that the mutant defect is detectable by two weeks, before the first age examined in that study, with central extension in clear corneas; and that wild-type corneas silence the endothelial and lymphatic program between P15 and adulthood whereas mutant corneas do not. That *Prox1*, *Flt4* and *Figf* (a lymphatic transcription factor, its receptor VEGFR3 and its ligand VEGF-D) differ individually at P3/P4 fits the lymphatic predominance they report, but CD31 does not distinguish blood from lymphatic endothelium, and the studies differ in allele (Sey, an exon 8 nonsense mutation, versus the Sey-Neu splice-donor mutation), background (129S1 versus outbred Swiss) and ages examined, so we cannot say which vessel type persists in *Pax6^Sey-Neu/+^*corneas. The nerve phenotype shows a parallel delay: fine epithelial fibers that are readily apparent in wild-type corneas by one week were not prominent in mutant corneas until about four weeks (Figure 1; Supplementary Figure 1D–G), so both innervation and vascular regression appear to be arrested short of the mature state, as the “frozen” developmental state predicts.

Our nerve data agree with the published mouse phenotype but do not settle whether a neurotrophic deficit causes the keratopathy. Leiper et al. found that trigeminal neurites reach the periphery of the *Pax6^+/−^*cornea earlier than in wild type (E12.5), that stromal projections are disorganized by E16.5, and that adult epithelium is less densely innervated with a disrupted radial pattern; Javidjam et al. described central stromal nerves, loss of the subbasal vortex and microneuromas from grade 1, when the central cornea is still transparent (Javidjam et al., 2024; Leiper et al., 2009). We find the same central stromal fascicles and loss of the central whorl in postnatal corneas, with density reduced from one week, so the abnormal pattern is established well before opacity, and a pilot (two libraries per group) detected no genotype effect in the trigeminal ganglion while the cornea was transparent. Leiper et al. concluded that *Pax6* dosage controls nerve patterning non-cell-autonomously, with normal neurotrophic support and little evidence that AAK represents neurotrophic keratitis, whereas Javidjam et al. proposed that loss of epithelial basement membrane lets stromal nerves invade the epithelium and creates an early neurotrophic deficit. Our data are compatible with either; because we measured neither nerve function nor epithelial consequences, the neurotrophic model is supported here as an early phenotype and untested as a cause.

Two further points concern how the stromal fascicles arise. Stromal fascicles persisted in the central cornea of mutant corneas throughout postnatal development (Figure 1), yet transparent adult mutant corneas lacked central stromal fascicles whereas mixed and opaque corneas retained them (Figure 3). The latter differs from the central stromal nerves that Javidjam et al. described in transparent grade 1 corneas, and because our design is cross-sectional we cannot say whether fascicles are eventually cleared in corneas that stay clear or persist in those that become opaque. Stromal nerves are also not specific to opacity: after epithelial abrasion, regenerating fibers can form clustered subbasal patterns resembling those of our transparent mutants (Meneux et al., 2025), whereas removal of the central epithelium is followed by thick stromal nerves in the central cornea (Di et al., 2017). One possibility is that the integrity of the epithelial basement membrane determines whether an altered microenvironment disturbs only the subbasal plexus or also the stroma. This would be consistent with the basement-membrane loss reported in this model (Javidjam et al., 2024), but we did not examine the basement membrane in the animals reported here, so it remains a hypothesis.

The epithelial models cannot be tested here: corneas were dissected without limbus, and no panel contains epithelial identity, differentiation or stem-cell genes. The published mouse data argue against a single epithelial explanation. Collinson et al., using the Sey-Neu allele studied here, found fewer, larger limbal stem-cell clones, cell-autonomous depletion of mutant stem-cell progeny in chimeras, and diverted centripetal epithelial migration that wild-type cells restored, concluding that both a stem-cell niche deficiency and non-autonomous guidance cues contribute (Collinson et al., 2004). In the same allele Ramaesh et al. found a thinner epithelium with peripheral goblet cells, weak K12 in older mice, and corneas that were vascularized or infiltrated in many adults but clear in some (Ramaesh et al., 2003); in later work they found delayed and reduced K12 expression, peripheral goblet cells and increased proliferation in *Pax6^+/−^* corneal epithelium, and concluded that low *Pax6* may directly impair differentiation and that AAK is not entirely explained by limbal stem cell deficiency; Mort et al. found modest effects of *Pax6* dosage on stem-cell clone numbers but pronounced stromal and endothelial abnormalities; and Javidjam et al. reported a thin, fragile epithelium with loss of collagen IV, fewer cells expressing stem-cell markers, and later MUC5AC expression (Javidjam et al., 2024; Mort et al., 2011; Ramaesh et al., 2005). Because *Pax6* is expressed throughout the corneal epithelium (Koroma et al., 1997), the stromal, neural and vascular abnormalities we find in clear corneas by two weeks could be non-autonomous consequences of epithelial dysfunction, making the epithelial and the vascular and neural models successive rather than competing; testing this requires an epithelium-restricted manipulation.

A chronic wound-healing state describes the transition to opacity better than its origin. Opaque corneas show a proteolytic, myofibroblast and inflammatory program, but the earliest differences in clear corneas are vascular and neural, and inflammatory infiltrates were seen in only a minority of mutant eyes in the published model (Javidjam et al., 2024). Anterior-segment dysgenesis cannot be excluded: keratolenticular adhesion is present in all Sey eyes from before birth yet did not appear to affect vascular ingrowth (Javidjam et al., 2024), and we did not assess lens–cornea attachment in Sey-Neu eyes. A persistent vascular and lymphatic program could also compromise immune privilege and, because the CD31-positive structures we detected lie in the stroma, promote stromal remodeling through inflammatory signaling; we did not assess immune cells, so this link is a hypothesis.

In summary, the data support arrested postnatal maturation of vascular and lymphatic identity and abnormal nerve patterning as early, cornea-intrinsic features of the clear mutant cornea; place the wound-healing, myofibroblast and myelination phenotypes at the later transition to opacity; and are silent on limbal stem cell deficiency, epithelial maldifferentiation and anterior-segment dysgenesis, which remain candidates for the upstream defect.

### What the transcriptional data do and do not explain

The nerve phenotype is present before any loss of transparency (Figures 1–3), but its transcriptional counterpart is narrow. In the adult cornea, with both genotypes transparent, the axon-guidance panel shows no program-level change: an excess of −0.34 confidently expressed genes against a detectable limit of 4.34, and −1.59 confidently silent against 3.59 — a bounded negative in the same libraries in which the vascular and lymphatic panel gains 13.4 and loses 13.5. Thirteen of the 92 axon-guidance genes differ at all, and the median absolute fold change across the panel is 0.30 log2.

What does differ is coherent. *Ephb2* is higher in the mutant at every age (+0.77, +1.24 and +2.54 log2 at P3/P4, P15 and in the transparent adult), *Unc5c* is lower and no longer confidently detected in the adult mutant cornea, *Reln* is categorically higher and *Dab1* lower, and *Cxcl12*, *Sema3d* and *Epha4* complete the list. These fall within the window of nerve ingrowth and subbasal plexus formation, consistent with non-autonomous regulation of nerve patterning by epithelial *Pax6* dosage (Leiper et al., 2009) and with established roles for guidance cues in corneal nerve architecture (Müller et al., 2003; Spurlin and Lwigale, 2013), but they are candidate contributors to a misrouting phenotype, not an explanation of one.

A small transcriptional signal is what this design would recover from a substantial structural phenotype. The program-level test counts genes entering or leaving confident expression, so a bounded negative does not exclude changes in level (ten of the panel’s 61 testable genes changed); a haploinsufficiency model is expected to produce effect sizes of this magnitude; much of the panel is undetermined rather than unchanged; and whole-cornea RNA averages over a regionally heterogeneous innervation phenotype and cannot resolve spatial gradients.

The vascular phenotype is the one for which imaging and transcription agree on timing. Peripheral CD31-positive structures are present in both genotypes at birth, are subsequently not detected in wild type, and become more frequent in *Pax6^Sey-Neu/+^* corneas, extending into the central cornea in 3 of 7 and 3 of 5 corneas at 2 and 3 weeks, whereas no wild-type cornea showed central labeling (Supplementary Table 3; these are detection counts in small groups, not tested statistically, and the wild-type 1-week group is a single animal). The endothelial and lymphatic set is higher in the mutant cornea from P3/P4 onward for nine of its eleven members, although at that age neither the individual genes nor the panel reaches significance; by P15 the panel carries an excess of 10.5 confidently expressed genes, and in the transparent adult an excess of 13.4 expressed and 13.5 fewer silent genes.

### Reduced *Pax6* dosage establishes an early, cornea-intrinsic corneal defect

Three observations place the origin of these changes in the cornea and in the first two postnatal weeks. Fifty panel genes already differ between genotypes at P3/P4 or P15, ages at which every cornea examined was transparent, and 74 differ across the three comparisons made while both genotypes were clear. Both the nerve deficit and the first intracorneal vessels appear within the same window. And pilot sequencing of trigeminal ganglia from the same animals detected no genotype effect on any of the three panels at any age, while an internal age control run through identical code detected 11,468 of 20,287 tested genes differentially expressed with age.

A transparent mutant cornea is nonetheless not pre-pathologic: by two weeks it is already vascularized in most animals and its innervation is abnormal. The state is an established, cornea-intrinsic defect that persists in a clinically clear cornea, so the adult signature is partly a signature of the pathology itself, superimposed on differences established much earlier (Collinson et al., 2004; Javidjam et al., 2024; Mort et al., 2011; Ramaesh et al., 2003; Ramaesh et al., 2005; Sunny et al., 2022).

### A failure to withdraw the vascular and lymphatic program

Postnatal maturation of the wild-type cornea is predominantly a silencing process: of 144 status crossings among 58 panel genes, 104 are genes turning off with age. Those crossings fall into two phases that differ in kind: between P3/P4 and P15 the wild-type cornea silences guidance and matrix genes (including *Dcc* and *Epha5*), and between P15 and adulthood it silences endothelial and lymphatic identity (*Pecam1*, *Cdh5*, *Flt4*, *Ptprb*, *Lyve1*, *Ccl21a* and *Rgs5*). The mutant executes the first phase and not the second: of the 25 genes that cross in wild type only, 19 are vascular or lymphatic, and eleven of them, those above plus *Emcn*, *Tie1*, *Klf2* and *Notch4*, are not detected to change status in the mutant at any interval.

The pooled expression levels show that this is a failure of withdrawal rather than an ectopic activation. Nine of eleven endothelial and lymphatic genes are already higher in the mutant cornea at P3/P4; the wild-type cornea then falls by roughly an order of magnitude between P15 and adulthood, to under one count per million for most of the set, while the mutant remains between roughly 1 and 12. The mutant cornea does not switch this program on. It never turns it off.

Among the secreted regulators that could sustain it, the VEGF family is altered earliest and most consistently: *Figf*, encoding VEGF-D, is higher in the mutant at every age (+2.03, +1.90 and +1.01 log2) and *Vegfa* from P15 onward (+1.22 and +0.67), matching *Flt4* (VEGFR3) failing to be silenced, whereas *Kdr* (VEGFR2) is undetermined in every comparison. *Sema3A*, which maintains corneal avascularity by inhibiting VEGF-induced angioblast migration (McKenna et al., 2014), is confidently expressed in all three adult groups and is not detected to differ in any comparison.

Corneal avascularity is thus an actively consolidated state: both genotypes begin with a broadly expressed vascular and lymphatic signature (136 of 177 genes confidently expressed in the wild-type cornea at P3/P4), and only the wild type dismantles it.

### Segmental stromal myelination reflects an opacity-coupled change in Schwann cell state

In the normal cornea, stromal nerves are held in an unmyelinated state by coordinated interactions among axons, Schwann cells and the local epithelial–stromal environment (Bargagna-Mohan et al., 2021; Conrad et al., 2009; Courson et al., 2019; Frutos-Rincón et al., 2022; Mirmoeini et al., 2023; Müller et al., 2003; Stepp et al., 2017). In *Pax6^Sey-Neu/+^* corneas we find focal, segmental MBP-positive myelination that corresponds to regions of opacity, interspersed with unmyelinated fibers within the same fascicles.

The transcriptional data align with this at the level of Schwann cell state: Egr2/Krox20, the master pro-myelinating transcription factor, is confidently expressed only in opaque cornea; *Pou3f1* moves stepwise from silent in wild type through indeterminate in the transparent mutant to expressed in the opaque cornea; and *Sox2*, a marker of Schwann cell dedifferentiation, crosses from silent to expressed only with opacity, whereas *Sox10* is indeterminate in all three adult groups. These are the myelination panel’s only categorical calls.

Two features argue that this is a second, opacity-coupled event superimposed on an already-altered environment, not the continuation of a program running in the clear cornea. While the cornea is transparent the direction of change is away from myelin (*Id2* is 1.02 log2 and *Pmp22* 0.71 log2 lower in the transparent mutant, and *Pmp22* rises only with the transition), and *Id2* is reduced at P15 and in the adult irrespective of transparency, so it indexes genotype rather than opacity. Genotype-driven changes in the microenvironment therefore appear to create a permissive but not yet inductive state, with an opacity-associated second signal required to engage myelination. A further argument points the same way. Presumably the same population of sensory neurons innervates transparent and opaque corneas, so axonal signals alone would not readily explain why myelinated fibers are confined to opaque regions; a local change in the corneal environment is the more likely trigger. The canonical axonal signal for Schwann cell fate, *Nrg1*, was not detected in corneal tissue at any age, consistent with its axonal origin, and its receptor *Erbb3* was only modestly higher in the transparent mutant (+0.26 log2), so this pathway can be assessed here only from the glial side. Compact myelin around stromal fibers might also add to light scattering within affected regions, a possibility we did not test.

*Mbp*, the transcript encoding the protein whose focal labeling defines the phenotype, returns a bounded negative in all four comparisons, excluding any effect larger than 1.5-fold; a focal phenotype affecting a minority of fibers within a minority of the tissue is what whole-cornea RNA would fail to resolve, and this discrepancy illustrates what bulk measurement costs.

Two alternatives to ectopic myelination should be kept open. Regenerating corneal nerves can acquire myelin (Namavari et al., 2011), so the phenotype may reflect a response to chronic damage, and corneal fibers that are NF200-positive during early postnatal development may be transiently myelinated normally (Meneux et al., 2025), in which case we would be observing remyelination.

### Corneal nerve defects are most consistent with a target-derived mechanism

Sensory innervation could be disrupted by changes in the neurons, in the tissue they grow into, or both, and our data are most consistent with the second, in agreement with Leiper et al. (2009). No panel gene differed between genotypes in the trigeminal ganglion at any age while the cornea was transparent (smallest adjusted P = 0.071); genes differed only in comparisons involving opaque animals, modestly and led by *Egr2*, a pattern consistent with a ganglion response secondary to corneal disease rather than a cause of it.

This bounds a hypothesis rather than establishing one: two libraries per group resolve only changes larger than about 1.5-fold (and for 208 of 745 comparisons not even that), only three panels were examined, and cornea-projecting neurons are a small fraction of the ganglion. The pilot argues that the ganglion’s transcriptional state is not the driver, not that the neuron is unaffected. *Pax6* is itself expressed in neurons of the ophthalmic region of the trigeminal ganglion, so a neuron-intrinsic contribution cannot be excluded on first principles; preliminary immunolabeling showed Pax6 in mutant ganglia at a level comparable to wild type, but this was not quantified.

### Opacity is associated with a broader remodeling phase

Comparing opaque with transparent mutant cornea, within genotype, the changes are broader, larger (34 of 130 vascular entries exceed one log2 unit with the transition, against 7 of 112 while the cornea remains clear) and different in kind: the axon-guidance panel reaches program-level significance only here, the myelination panel makes its only categorical calls here, and Mmp3 (+4.17), Acta2 (+3.02), *Epha7* (+3.09), *Flt1* (+2.78) and *Adam8* (+2.65) describe a proteolytic, myofibroblast and inflammatory signature largely absent from the transparent comparisons (Acta2 is already higher at P3/P4, +3.55 log2, and Cxcl5 and *Adam8* differ categorically in the transparent adult). These parallel transcriptional programs implicated in corneal wound healing and pathologic remodeling (Bykhovskaya et al., 2016; Gibson et al., 2014; Hodgkinson et al., 2010; Kelwick et al., 2015; Liu et al., 2017; Rose et al., 2021; Wilson, 2020), and they correspond to clinical and experimental descriptions of aniridia-associated keratopathy, in which progressive vascularization and stromal fibrosis accompany loss of epithelial integrity and innervation (Eden et al., 2012; Ihnatko et al., 2016; Lagali et al., 2013; Lagali et al., 2018; Vicente et al., 2018).

This phase extends a process already under way rather than marking its onset, and it describes a state, not a demonstrated sequence: the tissue compared differs in fibrosis and inflammation as well as clarity, and transparent and opaque adults are different animals, so a transparent cornea here is not the precursor of an opaque one there. Within mixed corneas, stromal fascicles and MBP-positive fibers were confined to opaque regions (Figures 3–5), which points to events arising locally rather than to a uniform change across the cornea.

### An integrated model of *Pax6*-dependent corneal remodeling

Taken together, the imaging and transcriptional data support a staged account. At birth the two genotypes are alike: stromal fascicles run throughout the cornea, nerve density does not differ, and both carry a broad vascular and lymphatic transcriptional signature, with peripheral CD31-positive structures but no central vessels. In the first postnatal days the earliest molecular divergence appears (*Prox1*, *Flt4* and *Figf*). Between one and two weeks both phenotypes emerge together: nerve density diverges, stromal fascicles fail to clear, CD31-positive structures persist and extend centrally, and the vascular program-level excess reaches significance at P15. The wild-type cornea then completes a second silencing phase that the mutant does not, leaving a transparent adult cornea whose guidance and myelination programs are indistinguishable from wild type at panel level but whose vascular and lymphatic program carries an excess of roughly thirteen genes, whose innervation is reduced and disorganized, and which is vascularized in most animals. Loss of clarity marks a later, broader remodeling with proteolytic, myofibroblast and inflammatory programs, a change in Schwann cell state, and ectopic compact myelin.

The model asserts no causal arrow between vessels and nerves: the two diverge in the same window, nothing in these data orders them, and at one week the nerve deficit is already present while CD31 labeling is detected in only one of three mutant corneas.

### Implications for mechanisms and therapeutic intervention

These findings place the earliest measurable changes in the first two postnatal weeks, while the cornea is still transparent. Two consequences follow.

First, a clear cornea in *Pax6* haploinsufficiency is not an unaffected cornea: by the time transparency can be assessed the vascular program has failed to withdraw, vessels have entered the tissue in most animals, and innervation is reduced, so an intervention window is open while the cornea still appears clear, with a vascular readout as a candidate marker. Approaches that promote epithelial repair, target fibrotic remodeling or support regeneration (Gong et al., 2024; Setiawan and Kamarudin, 2024; Trujillo Cubillo et al., 2024; van Velthoven et al., 2023) would be candidates within it, consistent with evidence for an early therapeutic window in aniridia-associated keratopathy (Fries et al., 2023). The corresponding window in the mouse, the first two postnatal weeks, spans eye opening (about P12–P14), and the most effective time for intervention may be soon afterward, before opacity develops. If the one- to two-week mouse cornea is taken to correspond roughly to the human cornea around birth, this fits the clinical observation that aniridic corneas are usually clear at birth yet show subtle ocular-surface abnormalities during the first decade (Lagali et al., 2020; Latta et al., 2021).

Second, because the guidance environment is close to normal in the transparent mutant cornea, the nerve phenotype is unlikely to be corrected by manipulating guidance cues in the adult; if a target exists it is upstream, in whatever *Pax6*-dependent process fails to switch the vascular and lymphatic program off. Because corneal nerves and Schwann cells support epithelial maintenance and regeneration (Bigit et al., 2025; Mirmoeini et al., 2023), a nerve deficit established early could feed back on the epithelium. Nerve and epithelial dysfunction could then reinforce one another, since epithelial breakdown may in turn cause further nerve loss.

Three experiments would test this account: same-cornea CD31 and TuJ1 labeling at one and two weeks, to determine whether nerve disorganization is spatially coincident with vessels; localization of *Ephb2*, *Unc5c*, *Reln*, *Cxcl12* and *Sema3d* at P15, to address the principal limitation of whole-cornea measurement; and an epithelium-restricted, postnatal-onset manipulation of *Pax6*, to test whether the vascular and neural phenotypes are non-autonomous consequences of epithelial dosage and to separate cornea-intrinsic from ganglion-intrinsic contributions by design.

### Limitations

Several constraints apply. Timing: whether the genotype effect changes with age is undetermined for more than 93% of genotype-by-age tests, so these data neither distinguish onset from progression nor establish constancy. Confounding: opaque and transparent corneas differ in fibrosis and inflammation as well as transparency and come from different animals, and the comparison that would sum genotype and transparency is reported only in Supplementary Figure 7. Measurement: whole-cornea RNA of curated panels does not localize transcripts to cell types, retains a variable amount of limbus where CD31 labeling is present in both genotypes, and does not assess extracellular matrix or epithelial-identity programs (Fini and Stramer, 2005; Wilson, 2020), which bear on the epithelial and stem-cell models above; 28 of the 92 axon-guidance genes are also curated as vascular or lymphatic. Resolution: of 831 reportable tests, 504 (61%) are undetermined, and a typical undetermined test leaves open an effect close to two-fold, so similarity between wild type and the transparent mutant is demonstrated for the panel rather than for most individual genes (only nine of 61 axon-guidance tests returned a bounded negative). The immunolabeled corneas were not those sequenced, and the transparent adult sequencing group was classified by gross transparency rather than vascular status; given that 19 of 23 adult mutant corneas carried intracorneal vessels, it most likely included vascularized corneas. Finally, published mouse models differ in *Pax6* allele, background, ages and methods, and the AAK phenotype in humans depends on *PAX6* mutational status (Lagali et al., 2020), so differences in onset and severity from earlier studies may reflect allele or background.

## Methods

### Sex as a biological variable

Male and female mice were used in all experiments. Adult corneas analyzed by immunofluorescence comprised 5 female and 8 male *Pax6*^+/+^ and 14 female and 18 male *Pax6*^Sey-Neu/+^ corneas (Supplementary Table 1). Sequencing libraries were assembled without regard to sex; donor sex was recorded for adult pools and, for every library, pool composition was verified from *Xist* and Y-linked gene expression, which agreed with the recorded sex in all 15 adult corneal pools (Supplementary Table 13). The male fraction of each pool was carried as a covariate in every corneal model, so that a difference in sex composition between groups could not be read as a genotype or transparency effect. Sex was not recorded for the newborn, 1-week, 2-week and 4-week mice used for the developmental CD31 and nerve series.

### Animals

*Pax6*^Sey-Neu/+^ mice carry the Small eye Neuherberg allele, a G-to-T transversion at the +1 position of the intron 10 splice donor of *Pax6* (Hill et al., 1991). Founder mice, provided by Dr. Nadean Brown, arrived in 2000; beginning around 2002–2004 the allele was moved onto outbred Swiss backgrounds (ICR/CD-1, Charles River Laboratories; Swiss Webster, Taconic Biosciences), where it has been maintained since (Kim and Lauderdale, 2006); *Pax6*^+/+^ littermates served as controls. Mice were housed at 22 °C on a 12-hour light/12-hour dark cycle. Adults were euthanized by CO₂ followed by cervical dislocation, and pups by decapitation. Heterozygous mutants were identified by reduced eye size and confirmed by PCR genotyping. Genomic DNA was isolated from ear punches by the salting-out method (Miller et al., 1988). A 335-bp fragment spanning intron 10 and the mutation site was amplified with a forward primer in exon 10 (5′-GAGGAACCAGAGAAGACAGGC-3′) and a reverse primer in exon 11 (5′-GCATAGGCAGGTTGTTTGCC-3′), and digested with *Hin*cII (New England Biolabs R0103S). The mutation creates a *Hin*cII site, so the mutant amplicon is cleaved (about 107 and 228 bp) and the wild-type amplicon is not, as described for this allele (Quinn et al., 1996). Products were resolved on an agarose gel.

### Corneal whole-mount immunolabeling

Immunolabeling followed a published protocol (Yun et al., 2021) with modifications. Immediately after euthanasia, eyes were enucleated and placed in 1.3% paraformaldehyde (PFA) in 1× PBS on ice for 1 hour in total; after 30 minutes in fixative the corneas were dissected and returned to fixative for the remaining time. 1.3% PFA was made by diluting 4% PFA (FD NeuroTechnologies PF101). Corneas were washed in 1× PBS, permeabilized for 1 hour in 1% Triton X-100 in 1× PBS at room temperature, and blocked for 1 hour at room temperature in 5% serum (normal goat or normal donkey serum), 0.2% bovine serum albumin and 0.3% Triton X-100. Primary antibodies, rabbit anti-βIII-tubulin (TuJ1; 1:4,000) and goat anti-myelin basic protein (MBP; 1:500), were applied overnight at 4 °C. Corneas were washed 3 times in 1× PBS with 0.1% Tween 20 for 15 minutes each and incubated for 2 hours at room temperature with Alexa Fluor-conjugated secondary antibodies at 1:1,000: goat anti-rabbit 647 or donkey anti-rabbit 647 for TuJ1 (the two gave comparable patterns) and donkey anti-goat 488 for MBP. For TuJ1/MBP co-labeling, donkey secondary antibodies were used with donkey serum as the blocking serum. After washing, nuclei were counterstained with DAPI (300 nM, 10 minutes, room temperature). Corneas were washed, given relaxing cuts (at least 4 per cornea; the number varied) to allow flat mounting, and mounted on slides using 70% glycerol in 1× PBS and glass coverslips. TuJ1 and MBP labeling was performed in 1.3% PFA-fixed corneas, except for the TuJ1-labeled corneas shown in Supplementary Figure 3, which were fixed in 2% PFA. The higher PFA concentration made uniform labeling more difficult, but the corneas that did label showed patterns comparable with 1.3% PFA fixation; a subset with notable fiber morphology is shown in Supplementary Figure 3. Except at the newborn time point, corneas at each age came from several litters (Supplementary Table 1). Antibodies, kits and reagents are listed in Supplementary Table 4. No-primary/secondary-only controls were used, and MBP-positive limbal fibers served as an internal positive control.

### CD31 immunolabeling

Vascular structures were labeled with the same protocol with the following modifications: corneas were fixed in 4% PFA for 30 minutes or 2% PFA for 1 hour at room temperature (initially they were fixed with 4% PFA but changed to 2% PFA to perform dual labeling with TuJ1), the primary antibody was rat anti-CD31 (BD Pharmingen 557355; 1:2,000), and the secondary antibody was donkey anti-rat 488 (1:1,000). Only corneas blocked in normal goat serum were included in the CD31 analysis. Mouse ages are given in the Ages table (Supplementary Methods); corneal numbers are in Supplementary Table 3.

### Confocal and transparency imaging

Corneas were imaged on a Zeiss LSM 900 confocal microscope controlled by ZEN 3.6 (Blue) software, which was also used for image acquisition and processing. Objectives were 2.5×/0.085, 10×/0.3, 20×/0.8 (dry), 40×/1.4 (oil) and 63×/1.4 (oil). Fluorescence filter sets for direct viewing were DAPI (blue), FITC (green), TAMRA (orange), mCherry (red) and Cy5 (far-red). Imaging used diode lasers (405, 488, 561 and 640 nm), and emission was collected with two gallium arsenide phosphide photomultiplier tube (GaAsP-PMT) detectors. Z-step size was 3 µm for newborn, 1-week and 2-week 10× TuJ1 tile imaging and 4 µm for 4-week, 2-month and adult TuJ1 and TuJ1/MBP 10× tile imaging, except in Supplementary Figure 3, where panels A and C were imaged at 4.5 µm steps and panel B at 4 µm steps; CD31 imaging ranged from 3 µm to 7 µm depending on tissue thickness. Tile scans at 10× covered the entire corneal surface through its full thickness so that fibers in all layers were captured. Higher-magnification fields were acquired at 20× with a 2× crop at 1 µm step size and, for XZ reslices of CD31 labeling, at 40×. Acquisition settings were held constant across genotypes within an experiment. Images were color-coded by Z position in Fiji using the temporal color code and Z-stack depth color code plugin, from superficial/epithelial (green) to deep/stromal (red) for TuJ1; the color scale differed for CD31 and MBP. For TuJ1, maximum-intensity projections spanned the entire corneal thickness. For MBP and CD31, maximum-intensity projections were made from the selected optical sections that contained labeling, to avoid excess background; for the higher-magnification MBP fields in Figure 5, projections comprised 1-µm optical sections spanning the fibers of interest (selected by one investigator, S.K.M., to show the different kinds of MBP labeling). Transparency was then documented on a Zeiss SteREO Discovery.V12 stereomicroscope: after immunolabeling and confocal imaging, each cornea was removed from the slide, washed in 1× PBS, placed in PBS on a clean new slide, and photographed against a matte black background under incident illumination (ring light) at fixed magnification (20×), lamp settings and camera exposure; all images for a comparison were collected together using an AxioCam camera. Because the background is black and the light is incident rather than transmitted, the image records light backscattered by the tissue: a transparent cornea shows the black background and appears dark, whereas regions that scatter light appear bright.

### Transparency assessment

Corneal transparency was assessed *ex vivo* on the isolated cornea. The clinical staging system for aniridia-associated keratopathy (Lagali et al., 2020) is defined by slit-lamp features of the living eye that cannot be evaluated in a dissected cornea and was not applied; the categories used here are therefore not equivalent to clinical grades and are not a severity series. For corneas used for immunofluorescence, the classification was made from the standardized backscatter images described above and scored as transparent (no opaque region), mixed (one or more discrete opaque regions within an otherwise transparent cornea) or opaque (opacity across most of the corneal surface); the classes were treated as unordered categories. The impression at dissection guided tissue allocation only and was superseded by the image-based classification. A wild-type cornea imaged under the same illumination served as the reference for a fully transparent cornea. Classification was performed by a single investigator (S.K.M.), who was not masked to genotype. Corneas for RNA sequencing and electron microscopy were not imaged after labeling; their assignment was made at dissection on fresh tissue (see Transmission electron microscopy and RNA sequencing, below).

### Nerve fiber density quantification

Nerve fiber density was quantified in Fiji (ImageJ 1.54; Schindelin et al., 2012; Schneider et al., 2012) by a single investigator (S.K.M.). Because the relaxing cuts leave empty space at the periphery of a flat-mounted cornea, density was measured within regions of interest (ROIs) rather than over the whole image. The mean diameter of each cornea was calculated from two perpendicular diameter measurements and used to define the central 50% of corneal area (a circle of about 71% of the corneal diameter). Three circular ROIs of equal area were placed on each cornea: one ROI inside the central 50% of corneal area and two ROIs outside it. Peripheral ROIs were placed so that no more than 20% of each ROI fell within the central region and so that limbal fibers were excluded. Regional ROIs were used to capture regional differences, which matter particularly in mutant corneas, where opacity was not confined to one region. ROI area was held constant between *Pax6*^+/+^ and mutant corneas at each age and was scaled with corneal size across ages. “Entire cornea” density (Fig 1R, Fig 2J, Fig 3N, Supplementary Figure 1D, G, H, I) is shown as the central ROI and the two peripheral ROIs of each cornea plotted as individual points (three per cornea), and includes both epithelial and stromal fibers. The three ROIs were placed to sample different parts of the cornea so as to assess the cornea as a whole, and no ROI overlapped with another. Pixels outside the ROI were cleared (Fiji “Clear Outside”). Within the ROI the image was contrast-enhanced, thresholded and skeletonized, all using default settings, and the skeleton was analyzed with the Analyze Particles command (Ferreira and Rasband, 2012). The number of particles was normalized to ROI area to give fiber density, which is expressed in arbitrary units throughout; this is a count-based index, not a measure of total fiber length or of the thresholded pixel fraction. In a test cornea, density in the central ROI was comparable to density across the entire central 50% of the cornea. Z-stacks were also divided manually into epithelial and stromal compartments, the boundary being the subbasal fiber plane as a morphological landmark, so that epithelial/subbasal and stromal fibers could be quantified separately; “total” density is from the projection across the full thickness. Each data point on the graph is from 1 ROI measurement; numbers of corneas and images per age, genotype and class are given in Supplementary Table 1.

### Scoring of CD31-positive structures

CD31-labeled corneas were scored for the location of intracorneal CD31-positive structures: limbus-restricted or extending into peripheral and/or central cornea (Figure 6; Supplementary Table 3). Scoring was performed by one investigator (S.K.M.) using the same central and peripheral regions as for the nerve fiber analysis. CD31 labels vascular endothelium and does not distinguish blood from lymphatic vessels. Results are reported as frequencies and were not tested statistically.

### Transmission electron microscopy

Freshly isolated corneas were fixed in 2.5% glutaraldehyde in 0.1 M sodium cacodylate buffer (pH 7.4) for 2–7 days at room temperature, rinsed in 0.1 M sodium cacodylate, post-fixed in 1% aqueous OsO₄ for 1–2 hours at room temperature, and stained *en bloc* overnight in 2% uranyl acetate at room temperature. Tissue was dehydrated through a graded ethanol series and propylene oxide and embedded in Eponate 12 resin. Sections of about 70 nm were cut on an ultramicrotome, stained with 2% uranyl acetate followed by Reynolds’ lead citrate (Reynolds, 1963), and examined on a JEOL 1400EX electron microscope at 80 kV. Electron microscopy was performed on separate animals from those used for immunolabeling (samples in Figure 4S–Z and Supplementary Table 2). Stromal nerve profiles were sampled from the central cornea or, where stated in the legend of Figure 4, the peripheral cornea. Axons were classified as unmyelinated Remak-type bundles (axons ensheathed by Schwann cell cytoplasm without a compact sheath) or as myelinated (axons surrounded by a compact multilamellar sheath) (Whitear, 1960; Müller et al., 2003). One cornea per condition was used for TEM. The age and sex of the animals used for TEM are given in the Ages table (Supplementary Methods).

### RNA sequencing

Corneas (limbus excluded) and the matching trigeminal ganglia were dissected from *Pax6*^+/+^ and *Pax6*^Sey-Neu/+^ mice at P3/P4, P15 and adulthood (the age of each library is given in Supplementary Table 13), snap-frozen in liquid nitrogen and stored at −80 °C. Adult corneas were assigned to the transparent or opaque group at dissection, by gross appearance on the stereomicroscope under incident illumination against a black background and by comparison with wild-type corneas dissected in the same session, before freezing; because these corneas were not fixed or imaged, no image record or second-scorer confirmation exists for this assignment. Corneas with focal opacity were not used. Each corneal library pooled 10 corneas from 5–6 mice (P3/P4) or 3 corneas from 3 mice (P15) or 2–3 mice (adult); each trigeminal library pooled the ganglia of the same animals as the matching corneal pool (Supplementary Table 13). Five corneal and two trigeminal libraries were prepared per group; one P3/P4 *Pax6*+/+ corneal sample failed the sequencing provider’s RNA integrity threshold (RIN 1.8) and was not sequenced, leaving four, while the trigeminal library from the same animals was retained. For two corneal replicates per group (2025 batch), RNA was isolated in-house (RNAqueous Micro Kit for P3/P4 corneas, RNeasy Mini Kit otherwise) and shipped to Novogene; the other three (2026 batch) were shipped as frozen tissue for isolation there. Libraries were prepared from poly-A-enriched mRNA with a non-directional protocol and sequenced on an Illumina NovaSeq X Plus (2 × 150 bp). All 48 libraries were processed together from the raw reads in a single run of a version-pinned pipeline (see Data availability). Reads were trimmed (Trimmomatic 0.39), aligned with HISAT2 2.2.1 to GRCm38 with GENCODE vM25 annotation, and counted with featureCounts (Subread 2.0.6; fragments, unstranded, all 48 libraries in one invocation). Strandedness was measured rather than assumed: assignment was equal at the two directional settings and highest unstranded. Processing details, quality metrics and tool versions are in Supplementary Methods.

### RNA-seq analysis

Genes with ten or more counts in at least two libraries were retained. Three curated panels (axon guidance, 92 genes; myelination, 25; vascular/lymphatic, 177) were analyzed; a gene on more than one panel was assigned a primary panel for multiplicity correction. All parameters are held in a single configuration file, and the full analysis is described in Supplementary Methods.

### Expression status

Counts were pooled across the libraries of a group and exact Poisson 95% limits computed on the pooled rate: a gene was expressed when the lower limit exceeded 1 CPM, not expressed when the upper limit fell below 0.2 CPM, and indeterminate otherwise; a call that changed when any one library was removed was set to indeterminate. Between groups, a gene differed categorically when expressed in one and not expressed in the other, or when its intervals were separated by at least twofold; it was testable when expressed in both, and otherwise silent in both or not assessable. No library was excluded; no library’s detection sensitivity fell more than three median absolute deviations below its stratum median, although one trigeminal library lay at that threshold (Supplementary Figure 8; Supplementary Methods).

### Differential expression

Testable genes were analyzed with DESeq2 in two models, each with a single factor combining age, genotype and transparency, plus sequencing batch and male fraction as covariates: a developmental model (transparent corneas at P3/P4, P15 and adult; n = 29) comparing genotypes within each age, and an opacity model (adult only; n = 15) comparing wild-type, transparent-mutant and opaque-mutant corneas pairwise. The transparent-mutant versus wild-type adult contrast is reported from the opacity model. Log₂ fold changes are unshrunk, with 95% confidence intervals. Genes not significant on the primary test were tested for equivalence to a 1.5-fold bound (altHypothesis = “lessAbs”) and reported as no change detected when significant, otherwise as undetermined. P values were Benjamini–Hochberg corrected within primary panel and contrast, with no independent filtering or fold-change threshold.

### Program-level and developmental comparisons

Differences between groups in the number of expressed and silent panel genes were tested against abundance-matched sets of non-panel genes from the same libraries (20,000 permutations), corrected across all panel-by-contrast tests. Within each genotype, transparent libraries were fitted across the three ages; the status rule identified genes whose status changed between ages, and each interval was also tested for quantitative change as above.

### Concordance and sensitivity

Every status crossing and categorical difference was re-tested without the status gate by standard replicate-aware DESeq2 (all panel genes, corrected over the panel) and by edgeR quasi-likelihood; calls not confirmed at adjusted P < 0.05 in the same direction are drawn as open symbols in Figures 7 and 8 (Supplementary Table 7). The status thresholds and categorical fold bound were varied over a grid (0.5/0.1 to 2/0.4 CPM; 1.5- to 3-fold; Supplementary Table 9). Of the 58 genes with a crossing at the configured setting, 38 to 58 still had a crossing, and 25 to 58 kept their wild-type-only, shared or Sey-only group, across the nine settings; at the configured ON/OFF thresholds (1/0.2 CPM) 53 to 58 kept their group as the fold bound varied, whereas moving the thresholds themselves changed the group of roughly half of the genes.

### Cross-dataset replication

GSE183742 (three wild-type and three *Pax6*^tm1Pgr/+^ adult corneas) was processed through the same pipeline and reference (strandedness measured: reverse-stranded) and analyzed as a separate study (Supplementary Table 14). The status rule was required to reproduce this study’s adult wild-type calls exactly before comparison. Agreement of wild-type status, and of the direction of categorical genotype differences, was compared with abundance-matched non-panel genes, and a fixed set of limbal, conjunctival, angle and retinal markers was compared between datasets to identify disagreements attributable to dissection margins (Supplementary Methods).

### Trigeminal ganglia

With two libraries per group, too few for expression-status calls, trigeminal libraries were analyzed with DESeq2 alone, using the corneal pre-filter, equivalence bound and correction within primary panel and contrast, in a developmental model (wild-type and transparent-mutant ganglia at the three ages; n = 12) comparing genotypes within each age, and an adult model (wild-type, transparent-mutant and opaque-mutant ganglia; n = 6) comparing the three groups pairwise and by an omnibus likelihood-ratio test. All trigeminal libraries were sequenced in one batch, so no batch term was fitted, and male fraction was not included as a covariate. A likelihood-ratio test for age in wild-type ganglia (n = 6) served as a positive control. Central-nervous-system and Schwann-cell markers were reported for every library to assess carry-over of central tissue at dissection (Supplementary Methods).

### Statistics

Nerve fiber density measurements from Fiji were analyzed and graphed in GraphPad Prism 11.0.2 (GraphPad Software, Boston, Massachusetts, USA; www.graphpad.com). Because group sizes were small and normality could not be assumed, nonparametric tests were used throughout. Two independent groups were compared with two-tailed Mann–Whitney tests (Mann and Whitney, 1947). More than two independent groups (the age series within a genotype and the adult transparency classes) were compared with Kruskal–Wallis tests followed by Dunn’s multiple-comparisons procedure (Dunn, 1964; Kruskal and Wallis, 1952). Genotype comparisons repeated at each developmental age were performed as multiple Mann–Whitney tests with Holm–Šídák adjustment to control the family-wise error rate (Holm, 1979; Šidák, 1967). All tests were two-sided with α = 0.05, and the test used for each analysis is stated in the corresponding figure legend. In box plots the box shows the median and interquartile range and the whiskers the minimum and maximum; each point is one ROI measurement (three ROIs per cornea, placed as described above; ROI measurements were the unit of analysis), and the numbers of corneas, mice and ROIs per group are given in Supplementary Table 1. No outliers were excluded. RNA-seq statistics are described above and in Supplementary Methods.

### Study approval

All animal procedures were approved by the University of Georgia Institutional Animal Care and Use Committee (protocols A2022 10-003-Y3-A0 and A2026 03-008-Y1-A0) and conducted in accordance with the NIH Guide for the Care and Use of Laboratory Animals and the Association for Research in Vision and Ophthalmology (ARVO) Statement for the Use of Animals in Ophthalmic and Vision Research.

### Data availability

Values underlying all graphs and reported means are provided in the Supporting Data Values file. Raw reads and the fragment count matrix for all 48 libraries are deposited in NCBI GEO (GSE348661); the re-analyzed public dataset is GSE183742. Analysis code and the run-provenance record are available on GitHub (https://github.com/Lauderdale-Lab/pax6-cornea-nerve-vascular-transparency; release v1.0.4, commit 43597c4) and archived at Zenodo (v1.0.4, https://doi.org/10.5281/zenodo.23347047; all versions, https://doi.org/10.5281/zenodo.22965379) under the MIT license. The upstream processing scripts that produced the count matrices from the raw reads (processing run reprocess_2026-09, commit 9182fd5) are available at https://github.com/Lauderdale-Lab/pax6-cornea-trigeminal-rnaseq-upstream (release v1.0.0, commit 938bb61) and archived at Zenodo (https://doi.org/10.5281/zenodo.22998976), together with the provenance record associated with the run; the release adds that record to commit 9182fd5 and changes no processing step. The values plotted in Figures 7–10 are in Supplementary Table 19. The complete output files of the reported run (https://doi.org/10.5281/zenodo.23349739) and the library metadata and processing quality-control tables (Supplementary Tables 13 and 18; https://doi.org/10.5281/zenodo.23350548) are archived at Zenodo. Original image stacks are available from the corresponding author on request.

## Author contributions

SM performed conceptualization, study design, coordination of experiments, performed experiments, tissue processing, data acquisition, analysis, interpretation, and drafted the manuscript. JL performed conceptualization, study design, funding acquisition, data analysis and interpretation, and assisted with manuscript writing, critical manuscript review, and revisions.

## AI disclosure

Microsoft Copilot (Microsoft Corporation, Redmond, WA, USA) and Claude (Anthropic; Fable 5.1) were used between August to October 2026 to draft and review the R and shell code for the RNA-seq analysis pipeline, to check the agreement between analysis outputs and the numbers quoted in the text and figure legends, and to edit the manuscript text. AI tools were not used to generate or modify data or images. The authors assume full responsibility for ensuring the integrity and accuracy of AI output presented in this article.

## Acknowledgments

Confocal imaging was performed in the Biomedical Microscopy Core at the University of Georgia using Zeiss LSM 900 confocal microscope. Electron microscopy was supported by the Emory University Robert P. Apkarian Integrated Electron Microscopy Core Facility (RRID: SCR_023537). We thank Dr. Sondip K. Biswas for his assistance with Electron microscopy. This study was supported in part by resources and technical expertise from the Georgia Advanced Computing Resource Center, a partnership between the University of Georgia’s Office of Research and Office of the Vice President for Information Technology. We thank Woodward Young for assisting with trigeminal ganglia dissections for RNA sequencing.

