## Supplementary for "Nerve and vascular abnormalities precede loss of corneal transparency in *Pax6*-haploinsufficient mice"

**Conflict of interest:** The authors have declared that no conflict of interest exists.

Supplementary Table 4. Antibodies, kits and reagents (in Supplementary Methods) ........ 9

**Supplementary Tables (provided as separate Excel files; source files also archived at Zenodo)**

Supplementary Table 5. Developmental status crossings ........ xlsx
Supplementary Table 6. Developmental trajectories ........ xlsx
Supplementary Table 7. Concordance of status calls with replicate-aware models ........ xlsx
Supplementary Table 8. Genotype-by-age interaction tests ........ xlsx
Supplementary Table 9. Threshold and stability sensitivity ........ xlsx
Supplementary Table 10. Expression status and assessability ........ xlsx
Supplementary Table 11. Categorical differences in expression status ........ xlsx
Supplementary Table 12. Trigeminal ganglion pilot ........ xlsx

Supplementary Table 13. Sequencing library inventory and quality metrics (48 libraries) ...................... xlsx
Supplementary Table 14. Cross-dataset replication against GSE183742 .............................................. xlsx
Supplementary Table 15. Curated gene panels and primary-panel assignment .................................... xlsx
Supplementary Table 16. Per-gene differential expression outcomes ................................................... xlsx
Supplementary Table 17. Program-level tests ......................................................................................... xlsx
Supplementary Table 18. Read processing and quantification (54 libraries)......................................... xlsx
Supplementary Table 19. Source data for Figures 7–10 and supplementary figures ............................. xlsx
Supplementary Table 20. Library depth and off-target tissue quality control .......................................... xlsx

Supporting Data Values for all graphed data and reported means are provided as a separate Excel file; the values plotted in Figures 7–10 are in Supplementary Table 19. Raw sequencing reads and the processed count matrix are deposited in NCBI GEO under accession GSE348661. Analysis code is archived at Zenodo (analysis pipeline v1.0.4, https://doi.org/10.5281/zenodo.23347047; all versions, https://doi.org/10.5281/zenodo.22965379; FASTQ-to-counts pipeline v1.0.0, https://doi.org/10.5281/zenodo.22998976). The complete output files of the reported run, from which Supplementary Tables 5–12, 14–17, 19 and 20 are drawn, are archived at Zenodo (https://doi.org/10.5281/zenodo.23349739), and the library metadata and processing quality-control tables (Supplementary Tables 13 and 18) at https://doi.org/10.5281/zenodo.23350548.

### Supplementary Methods: Animals, Imaging and Nerve Quantification

#### Study approval

All animal procedures were approved by the University of Georgia Institutional Animal Care and Use Committee (protocols A2022 10-003-Y3-A0 and A2026 03-008-Y1-A0) and conducted in accordance with the NIH Guide for the Care and Use of Laboratory Animals and the Association for Research in Vision and Ophthalmology (ARVO) Statement for the Use of Animals in Ophthalmic and Vision Research.

#### Sex as a biological variable

Male and female mice were included in all experiments. Adult corneas analyzed by immunofluorescence comprised 5 female and 8 male *Pax6*$^{+/+}$ corneas and 14 female and 18 male *Pax6*$^{Sey\text{-}Neu/+}$ corneas (Supplementary Table 1). Sequencing libraries were assembled without regard to sex; donor sex was recorded for adult pools and, for every library, pool composition was verified from *Xist* and Y-linked gene expression, which agreed with the recorded sex in all 15 adult corneal pools (Supplementary Table 13). The male fraction of each pool was carried as a covariate in every corneal model, so that a difference in sex composition between groups could not be read as a genotype or transparency effect. Sex was not recorded for the newborn, 1-week, 2-week and 4-week mice used for the developmental CD31 and nerve series.

#### Animals, husbandry and genotyping

*Pax6*$^{Sey\text{-}Neu/+}$ mice carry the Small eye Neuherberg allele, a G-to-T transversion at the +1 position of the intron 10 splice donor of *Pax6* (Hill et al., 1991). Founder mice, provided by Dr. Nadean Brown, arrived in 2000; beginning around 2002–2004 the allele was moved onto outbred Swiss backgrounds (ICR/CD-1, Charles River Laboratories; Swiss Webster, Taconic Biosciences), where it has been maintained since (Kim and Lauderdale, 2006). *Pax6*$^{+/+}$ littermates served as controls. Mice were housed at 22°C on a 12-hour light/12-hour dark cycle.

Ages of the animals used for each histological figure are listed in the table below, which is the single source for histology ages. The age of each RNA-seq library is given in Supplementary Table 13, which is the single source for library ages.

| Figure / item | Experiment | Timepoint | Age |
|---|---|---|---|
| Fig 1 (B–T) | TuJ1, postnatal series | Newborn | P0 |
| Fig 1 | TuJ1 | 1 week | P6, P7 |
| Fig 1 | TuJ1 | 2 weeks | P14, P16 |
| Fig 1 | TuJ1 | 4 weeks | P28–P33 |
| Fig 2 | TuJ1, adult | Adult | 4–7 months |
| Fig 3 | TuJ1 by transparency class, adult | Adult | 4–7 months |

| Figure / item | Experiment | Timepoint | Age |
|---|---|---|---|
| Fig 4 (A–R) | TuJ1 + MBP, adult | Adult | 5–7 months |
| Fig 4 (S–Z), Supplementary Table 2 | TEM, separate animals | Adult | 7 months (WT, Sey-T, Sey-M); 8.5 months (Sey-O); all female |
| Fig 5 | MBP high magnification | Adult | 5–7 months |
| Fig 6 (B–H) | CD31, adult | Adult (from P26) | P26 to 7 months; panels representative (Fig 6 legend) |
| Fig 6 data, Supplementary Table 3 | CD31, postnatal series | Newborn | P0/P1 |
| Supplementary Table 3 | CD31 | 1 week | P8 |
| Supplementary Table 3 | CD31 | 2 weeks | P15, P16 |
| Supplementary Table 3 | CD31 | 3 weeks | P22 |
| Supplementary Figure 1A–G | TuJ1 density, postnatal series | Newborn, 1, 2, 4 weeks | As Fig 1 |
| Supplementary Figure 1B | TuJ1, total fiber density in entire cornea, wild type | Newborn, 1, 2, 4 weeks; adult | As Figs 1, 2 |
| Supplementary Figure 1H–K | TuJ1 density, adult | Adult | As Figs 2, 3 |
| Supplementary Figure 2* | TuJ1 + MBP, 2-month cornea | 2 months | 2 months (P59) |
| Supplementary Figure 3 | TuJ1 whorl-like organization, adult | Adult | 4–8 months |
| Figs 7, 8; Supplementary Figures 4–8 | RNA-seq (cornea and trigeminal libraries) | Per library | See Supplementary Table 13 |

*Mice in Supplementary Figure 2 were examined at 2 months to determine whether the changes observed at 4 months or later were already evident at that age.

Heterozygous mutants were identified by reduced eye size and confirmed by PCR genotyping. Genomic DNA was isolated from ear punches by the salting-out method (Miller et al., 1988). A 335-bp fragment spanning intron 10 and the mutation site was amplified with a forward primer in exon 10 (5′-GAGGAACCAGAGAAGACAGGC-3′) and a reverse primer in exon 11 (5′-GCATAGGCAGGTTGTTTGCC-3′) and digested with HincII (New England Biolabs R0103S). The mutation creates a *Hin*cII (*Hin*dII) site, so the mutant amplicon is cleaved (about 107 and 228 bp) and the wild-type amplicon is not, as described for this allele (Quinn et al., 1996). Products were resolved on an agarose gel.

#### Whole-mount immunofluorescence for TuJ1 and MBP

Immunofluorescence followed a published whole-mount protocol (Yun et al., 2021) with modifications. Immediately after euthanasia, eyes were enucleated and placed in 1.3%

paraformaldehyde (PFA) in 1× PBS on ice for 1 hour in total; after 30 minutes in fixative the corneas were dissected and returned to fixative for the remaining time. 1.3% PFA was made by diluting 4% PFA (FD NeuroTechnologies PF101). Corneas were washed in 1× PBS, permeabilized for 1 hour in 1% Triton X-100 in 1× PBS at room temperature, and blocked for 1 hour at room temperature in 5% serum (normal goat or normal donkey serum), 0.2% bovine serum albumin and 0.3% Triton X-100. Primary antibodies, rabbit anti-βIII-tubulin (TuJ1; 1:4,000) and goat anti-myelin basic protein (MBP; 1:500), were applied overnight at 4 °C. Corneas were washed 3 times in 1× PBS with 0.1% Tween 20 for 15 minutes each and incubated for 2 hours at room temperature with Alexa Fluor-conjugated secondary antibodies at 1:1,000: goat anti-rabbit 647 or donkey anti-rabbit 647 for TuJ1 (the two gave comparable patterns) and donkey anti-goat 488 for MBP. For TuJ1/MBP co-labeling, donkey secondary antibodies were used with donkey serum as the blocking serum. After washing, nuclei were counterstained with DAPI (300 nM, 10 minutes, room temperature). Corneas were washed, given relaxing cuts (at least 4 per cornea, actual number is variable) to allow flat mounting, and mounted on slides using 70% glycerol in 1X PBS and glass coverslips. Antibodies, kits and reagents are listed in Supplementary Table 4. No-primary/secondary-only controls; MBP-positive limbal fibers serve as an internal positive control.

TuJ1 and MBP labeling was performed in 1.3% PFA-fixed corneas, except for the TuJ1-labeled corneas shown in Supplementary Figure 3, which were fixed in 2% PFA. The higher PFA concentration made uniform labeling more difficult, but the corneas that did label showed patterns comparable with 1.3% PFA fixation; a subset with notable fiber morphology is shown in Supplementary Figure 3. Except at the newborn time point, corneas at each age came from several litters (Supplementary Table 1).

#### Whole-mount immunofluorescence for CD31

Vascular structures were labeled with the same protocol with the following modifications: corneas were fixed in 4% PFA for 30 minutes or 2% PFA for 1 hour at room temperature (initially they were fixed with 4% PFA but changed to 2% PFA to perform dual labeling with Tuj1), the primary antibody was rat anti-CD31 (BD Pharmingen 557355; 1:2,000), and the secondary antibody was donkey anti-rat 488 (1:1,000). Only corneas blocked in normal goat serum were included in the CD31 analysis. Ages of the mice used for CD31 labeling are given in the Ages table; numbers of corneas are in Supplementary Table 3.

#### Confocal imaging

Corneas were imaged on a Zeiss LSM 900 confocal microscope controlled by ZEN 3.6 (Blue) software, which was also used for image acquisition and processing. Objectives were 2.5×/0.085, 10×/0.3, 20×/0.8 (dry), 40×/1.4 (oil) and 63×/1.4 (oil). Fluorescence filter sets for direct viewing were DAPI (blue), FITC (green), TAMRA (orange), mCherry (red) and Cy5 (far-red). Imaging used diode lasers (405, 488, 561 and 640 nm), and emission was collected with two gallium arsenide phosphide photomultiplier tube (GaAsP-PMT) detectors.

For 10× tile imaging, Z-step size was 3 µm for newborn, 1-week and 2-week TuJ1 and 4 µm for 4-week, 2-month and adult TuJ1 and TuJ1/MBP, except in Supplementary Figure 3, where panels A and C were imaged at 4.5 µm steps and panel B at 4 µm steps. CD31 imaging ranged from 3 µm to 7 µm depending on the tissue thickness. Tile scans at 10× covered the entire corneal surface through its full thickness so that fibers in all layers were captured. Higher-magnification fields were acquired at 20× with a 2× crop at 1 µm step size and, for XZ reslices of CD31 labeling, at 40×. Acquisition settings were held constant across genotypes within an experiment. Images were color-coded by Z position in Fiji (Schindelin et al., 2012) using temporal color code and Z-stack depth color code plugin, from superficial/epithelial (green) to deep/stromal (red) for TuJ1; the color scale differed for CD31 and MBP. For TuJ1, maximum-intensity projections spanned the entire corneal thickness. For MBP and CD31, maximum-intensity projections were made from the selected optical sections that contained labeling, to avoid excess background; for the higher-magnification MBP fields in Figure 5, projections comprised 1-µm optical sections spanning the fibers of interest (selected by one investigator, S.K.M., to show the different kinds of MBP labeling).

#### Backscatter imaging and assessment of corneal transparency

After immunolabeling and confocal imaging, each cornea was removed from the slide, washed in 1× PBS, placed in PBS on a clean new slide, and photographed on a Zeiss SteREO Discovery.V12 stereomicroscope against a matte black background under incident illumination, provided by a ring light, at fixed magnification, lamp settings and camera exposure; all images for a comparison were collected together using AxioCam camera at 20x magnification. Because the background is black and the light incident rather than transmitted, the image records light backscattered by the tissue: a transparent cornea shows the black background and appears dark, whereas regions that scatter light appear bright.

Transparency was assessed ex vivo on the isolated cornea. The clinical staging system for aniridia-associated keratopathy (Lagali et al., 2020) is defined by slit-lamp features of the living eye that cannot be evaluated in a dissected cornea and was not applied; the categories used here are therefore not equivalent to clinical grades and are not a severity series. For corneas used for immunofluorescence, the classification was made from the standardized backscatter images above and scored as transparent (no opaque region), mixed (one or more discrete opaque regions within an otherwise transparent cornea) or opaque (opacity across most of the corneal surface); the classes were treated as unordered categories. The impression at dissection guided tissue allocation only and was superseded by the image-based classification. A wild-type cornea imaged under the same illumination served as the reference for a fully transparent cornea. Classification was performed by a single investigator (S.K.M.), who was not masked to genotype. Corneas for RNA sequencing and electron microscopy were not imaged after labeling; their assignment was made at dissection on fresh tissue (see Transmission electron microscopy and Supplementary Methods: RNA-seq analysis).

#### Quantification of nerve fiber density

Nerve fiber density was quantified in Fiji (ImageJ 1.54; Schindelin et al., 2012; Schneider et al., 2012) by a single investigator (S.K.M.). Because the relaxing cuts leave empty space at the periphery of a flat-mounted cornea, density was measured within regions of interest (ROIs) rather than over the whole image. The mean diameter of each cornea was calculated from two perpendicular diameter measurements and used to define the central 50% of corneal area (a circle of about 71% of the corneal diameter). Three circular ROIs of equal area were placed on each cornea: one ROI inside the central 50% of corneal area and two ROIs outside it. Peripheral ROIs were placed so that no more than 20% of each ROI fell within the central region and so that limbal fibers were excluded. Regional ROIs were used to capture regional differences, which matter particularly in mutant corneas, where opacity was not confined to one region. ROI area was held constant between *Pax6*$^{+/+}$ and mutant corneas at each age and was scaled with corneal size across ages. “Entire cornea” density (Fig 1R, Fig 2J, Fig 3N, Supplementary Figure 1D, G, H, I) is shown as the central ROI and the two peripheral ROIs of each cornea plotted as individual points (three per cornea), and includes both epithelial and stromal fibers. The three ROIs were placed to sample different parts of the cornea so as to assess the cornea as a whole, and no ROI overlapped with another.

Pixels outside the ROI were cleared (Fiji “Clear Outside”). Within the ROI the image was contrast-enhanced, thresholded and skeletonized, all using default settings, and the skeleton was analyzed with the Analyze Particles command (Ferreira and Rasband, 2012). The number of particles was normalized to ROI area to give fiber density, which is expressed in arbitrary units throughout; this is a count-based index, not a measure of total fiber length or of the thresholded pixel fraction. In a test cornea, density in the central ROI was comparable to density across the entire central 50% of the cornea. Z-stacks were also divided manually into epithelial and stromal compartments, the boundary being the subbasal fiber plane as a morphological landmark, so that epithelial/subbasal and stromal fibers could be quantified separately; “total” density is from the projection across the full thickness. Each data point on the graph is from 1 ROI measurement; numbers of corneas and images per age, genotype and class are given in Supplementary Table 1.

#### Scoring of CD31-positive structures

CD31-labeled corneas were scored for the location of intracorneal CD31-positive structures: limbus-restricted or extending into peripheral and/or central cornea (Figure 6; Supplementary Table 3). Scoring was performed by one investigator (S.K.M.) using the same central and peripheral regions as for the nerve fiber analysis. CD31 labels vascular endothelium and does not distinguish blood from lymphatic vessels. Results are reported as frequencies and were not tested statistically.

#### Transmission electron microscopy

Freshly isolated corneas were fixed in 2.5% glutaraldehyde in 0.1 M sodium cacodylate buffer (pH 7.4) for 2–7 days at room temperature, rinsed in 0.1 M sodium cacodylate, post-fixed in 1% aqueous $OsO_4$ for 1–2 hours at room temperature, and stained en bloc overnight in 2% uranyl acetate at room temperature. Tissue was dehydrated through a graded ethanol series and propylene oxide and embedded in Eponate 12 resin. Sections of about 70 nm were cut on an ultramicrotome, stained with 2% uranyl acetate followed by Reynolds' lead citrate (Reynolds, 1963), and examined on a JEOL 1400EX electron microscope at 80 kV.

Electron microscopy was performed on separate animals from those used for immunolabeling. Stromal nerve profiles were sampled from the central cornea or, where stated in the legend of Figure 4, the peripheral cornea. Axons were classified as unmyelinated Remak-type bundles (axons ensheathed by Schwann cell cytoplasm without a compact sheath) or as myelinated (axons surrounded by a compact multilamellar sheath) (Whitear, 1960; Müller et al., 2003). Samples used in Figure 4S -Z and supplementary table 2 are different from the corneas used for immunolabeling. One cornea per condition was used for TEM. The age and sex of the animals used for TEM are given in the Ages table.

#### Statistics

Nerve fiber density measurements from Fiji were analyzed and graphed in GraphPad Prism 11.0.2 (GraphPad Software, Boston, Massachusetts, USA; www.graphpad.com). Because group sizes were small and normality could not be assumed, nonparametric tests were used throughout. Two independent groups were compared with two-tailed Mann–Whitney tests (Mann and Whitney, 1947). More than two independent groups (the age series within a genotype and the adult transparency classes) were compared with Kruskal–Wallis tests followed by Dunn's multiple-comparisons procedure (Dunn, 1964; Kruskal and Wallis, 1952). Genotype comparisons repeated at each developmental age were performed as multiple Mann–Whitney tests with Holm–Šídák adjustment to control the family-wise error rate (Holm, 1979; Šidák, 1967). All tests were two-sided with $\alpha = 0.05$, and the test used for each analysis is stated in the corresponding figure legend. In box plots the box shows the median and interquartile range and the whiskers the minimum and maximum; each point is one ROI measurement (three ROIs per cornea, placed as described above; ROI measurements were the unit of analysis), and the numbers of corneas, mice and ROIs per group are given in Supplementary Table 1. No outliers were excluded. RNA-seq statistics are described in Supplementary Methods: RNA-seq analysis.

#### Supplementary Table 4. Antibodies, kits and reagents

Source, catalog number and working dilution for the antibodies, enzyme, kits and reagents used. Anti-rat secondary antibody: donkey anti-rat Alexa Fluor 488 was used for CD31 labeling.

| Reagent | Supplier | Catalog no. | Working dilution |
|---|---|---|---|
| **Primary antibodies** | | | |
| Rabbit anti-βIII-tubulin (TuJ1) | BioLegend | 802001 | 1:4,000 |
| Goat anti-myelin basic protein (MBP) | Invitrogen | PA5-143589 | 1:500 |
| Rat anti-CD31 | BD Pharmingen | 557355 | 1:2,000 |
| **Secondary antibodies and counterstain** | | | |
| Goat anti-rabbit IgG, Alexa Fluor 647 | Invitrogen | A21244 | 1:1,000 |
| Donkey anti-rabbit IgG, Alexa Fluor 647 | Invitrogen | A31573 | 1:1,000 |
| Donkey anti-goat IgG, Alexa Fluor 488 | Invitrogen | A11055 | 1:1,000 |
| Donkey anti-rat IgG, Alexa Fluor 488 | Invitrogen | A21208 | 1:1,000 |
| DAPI | Sigma | D8417 | 300 nM |
| **Fixation, permeabilization and blocking** | | | |
| 4% paraformaldehyde (PFA) | FD NeuroTechnologies | PF101 | – |
| Triton X-100 | Sigma | X100 | – |
| Normal goat serum | Sigma | G9023 | – |
| Donkey serum | Sigma | D9663 | – |
| Bovine serum albumin | Fisher Scientific | BP1600-100 | – |
| Tween 20 | Sigma | P9416 | – |
| Glutaraldehyde | Sigma | G5882 | – |
| **Genotyping and RNA isolation** | | | |
| HincII restriction enzyme | New England Biolabs | R0103S | – |
| RNAqueous-Micro Kit | Thermo Fisher Scientific | AM1931 | – |
| RNeasy Mini Kit | Qiagen | 74104 | – |

#### Supplementary Methods references

Dunn, O.J., 1964. Multiple comparisons using rank sums. Technometrics 6, 241–252.

Ferreira, T., Rasband, W., 2012. ImageJ User Guide: IJ 1.46r. National Institutes of Health, Bethesda, MD.

Hill, R.E., Favor, J., Hogan, B.L., Ton, C.C., Saunders, G.F., Hanson, I.M., Prosser, J., Jordan, T., Hastie, N.D., van Heyningen, V., 1991. Mouse small eye results from mutations in a paired-like homeobox-containing gene. Nature 354, 522–525.

Holm, S., 1979. A simple sequentially rejective multiple test procedure. Scand J Stat 6, 65–70.

Kim, J., Lauderdale, J.D., 2006. Analysis of Pax6 expression using a BAC transgene reveals the presence of a paired-less isoform of Pax6 in the eye and olfactory bulb. Dev Biol 292, 486–505.

Kruskal, W.H., Wallis, W.A., 1952. Use of ranks in one-criterion variance analysis. J Am Stat Assoc 47, 583–621.

Lagali, N., Wowra, B., Fries, F.N., Latta, L., Moslemani, K., Utheim, T.P., Wylegala, E., Seitz, B., Kasmann-Kellner, B., 2020. PAX6 mutational status determines aniridia-associated keratopathy phenotype. Ophthalmology 127, 273–275.

Mann, H.B., Whitney, D.R., 1947. On a test of whether one of two random variables is stochastically larger than the other. Ann Math Stat 18, 50–60.

Miller, S.A., Dykes, D.D., Polesky, H.F., 1988. A simple salting out procedure for extracting DNA from human nucleated cells. Nucleic Acids Res 16, 1215.

Müller, L.J., Marfurt, C.F., Kruse, F., Tervo, T.M., 2003. Corneal nerves: structure, contents and function. Exp Eye Res 76, 521–542.

Quinn, J.C., West, J.D., Hill, R.E., 1996. Multiple functions for Pax6 in mouse eye and nasal development. Genes Dev 10, 435–446.

Reynolds, E.S., 1963. The use of lead citrate at high pH as an electron-opaque stain in electron microscopy. J Cell Biol 17, 208–212.

Schindelin, J., Arganda-Carreras, I., Frise, E., et al., 2012. Fiji: an open-source platform for biological-image analysis. Nat Methods 9, 676–682.

Schneider, C.A., Rasband, W.S., Eliceiri, K.W., 2012. NIH Image to ImageJ: 25 years of image analysis. Nat Methods 9, 671–675.

Šidák, Z., 1967. Rectangular confidence regions for the means of multivariate normal distributions. J Am Stat Assoc 62, 626–633.

Whitear, M., 1960. An electron microscope study of the cornea in mice, with special reference to the innervation. J Anat 94, 387–409.

Yun, H., Lathrop, K.L., St Leger, A.J., 2021. A whole-mount immunohistochemistry protocol for detection of mouse corneal nerves. STAR Protoc 2, 100734.

### Supplementary Methods: RNA-seq analysis

#### S1. Overview and design principles

The RNA-seq analysis was built around three questions that the imaging raised: which genes of three curated programs (axon guidance, myelination, vascular/lymphatic) the normal cornea expresses and silences as it matures; whether reduced *Pax6* dosage changes that; and which changes are associated with loss of transparency rather than with genotype alone. Each is a question about *expression status* as much as about fold change, because the imaging phenotypes are categorical (vessels present or regressed; nerves present or absent), and because the corneas were pooled, mixed-sex tissue from several animals, sequenced in two batches, with 4–5 libraries per group.

The analysis therefore separates two kinds of claim and tests each with a method suited to it. A *status* claim (a gene is confidently expressed, or confidently not expressed, in a group) is made from pooled counts with exact Poisson limits and a stability check. A *quantitative* claim (a gene is higher or lower between two groups in which it is expressed) is made with a negative-binomial model on replicate libraries. Genes that cannot support either claim are reported as such rather than forced into one. Two further layers test the framework itself: every status-level result is re-examined under a standard replicate-aware model without the status gate, and the status calls are recomputed in an independent dataset from a different laboratory and allele.

The guiding rule throughout was to prefer reporting no effect and being wrong to reporting an effect and being wrong: negatives are split into “no change detected, with an effect of 1.5-fold or larger excluded” and “undetermined”, and a status call that depends on which single library was retained is not reported as a call.

The complete pipeline is fourteen R scripts run in sequence from one driver (`run_all.R`). Every threshold, path and design choice is held in one configuration file (`00_config.R`), can be overridden by an environment variable for sensitivity runs, and is printed at the start of every run. Each script carries a numbered *Statistical basis* block citing the method it implements. No number in the manuscript was computed outside this pipeline. The released code reproduces every reported number from a fresh clone of its repository (Section S15).

#### S2. Samples, batches and covariates

Corneal libraries (n = 34) comprise wild-type (*Pax6*$^{+/+}$) and *Pax6*$^{Sey\text{-}Neu/+}$ corneas at P3/P4, P15 and adult (3–8 months), with adult mutant corneas classified as transparent or opaque at dissection, by gross appearance on the stereomicroscope (main Methods). Group sizes are P3/P4 wild type 4, mutant 5; P15 5 and 5; adult wild type 5, transparent mutant 5, opaque mutant 5. Fourteen trigeminal-ganglion libraries from a subset of the same animals were analyzed as a pilot (Section S14). Each corneal library pools 10 corneas from 5–6 mice (P3/P4), 3 corneas from 3 mice (P15) or 3 corneas from 2–3 mice (adult), of both sexes. The pool composition (animals, litters, and corneas or ganglia per pool), recorded donor sex and age of every library are given in Supplementary Table 13, which is the authoritative record; the values summarised here are for orientation.

Libraries were sequenced in two batches (2025: 13 corneal and all 14 trigeminal libraries; 2026: 21 corneal). Every corneal group contains libraries from both batches (two from 2025 and three from 2026, except P3/P4 wild type with one and three), so batch is estimable in every model with no empty batch-by-group cell. Batch enters every model as a covariate; it is never removed from the counts beforehand, because batch-corrected counts overstate the confidence of downstream tests (Nygaard et al., 2016). The two batches were sequenced under separate orders (Novogene X202SC24111367, 2025; X202SC26055473, 2026) with target yields of 12 and 15 Gb per library, and differ systematically in FastQC metrics (per-tile quality, per-base sequence content and GC content, 50–51% versus 44–49%), which is the physical basis for the covariate. A genome-wide check found a detectable batch effect in 13–50% of expressed genes depending on age stratum, so batch adjustment is not optional in this dataset (Leek et al., 2010); the P3/P4 stratum showed the weakest batch signal and the lowest correlation of $\log_2$ fold changes fitted with and without the term, and is treated as the least secure context for batch adjustment.

Because each library pools animals of both sexes, the male contribution to each pool was estimated from the summed counts-per-million of four Y-linked genes (*Ddx3y*, *Eif2s3y*, *Uty*, *Kdm5d*), scaled to the maximum within each tissue and age group. The result is a relative composition index on [0, 1], not an absolute male proportion, and is used only as a covariate (`male_frac`) so that a difference in sex composition between groups cannot be read as a genotype or transparency effect.

#### S3. Read processing and quantification

**Library chemistry.** Libraries were prepared by Novogene from poly-T-purified mRNA with random-hexamer priming and conventional second-strand synthesis (non-directional protocol, as stated in both batch reports) and sequenced on an Illumina NovaSeq X Plus in 2 × 150 bp paired-end mode. All read processing was performed together, from the raw reads, in a single run of the upstream pipeline (run `reprocess_2026-09`, commit `9182fd5`) on the University of Georgia Sapelo2 cluster under SLURM with Lmod modules; both batches were therefore processed with the same program versions and settings. Module versions are pinned in the pipeline (`config/modules.tsv`), recorded as reported by each tool at run time, and listed in Section S15.

**Trimming and QC.** Raw reads were assessed with FastQC 0.12.1 (Andrews, 2010). Adapter sequences were determined per library by parsing the FastQC over-represented-sequences module for sequences ≥ 12 nt annotated to a known adapter family (homopolymers excluded); none was detected in any library of either batch, so all were trimmed against the Trimmomatic `TruSeq3-PE` adapter file. Reads were trimmed with Trimmomatic 0.39 (Bolger et al., 2014; `PE`, `phred33`; `ILLUMINACLIP:TruSeq3-PE.fa:2:30:10:2:True LEADING:3 TRAILING:3 SLIDINGWINDOW:4:20 MINLEN:50`), with identical parameters for both batches. Only pairs in which both mates survived were carried forward (98.1–98.5% of pairs in 2026; 96.7–97.3% in 2025); orphaned reads were discarded. Yields were 49.2–73.2 million pairs per library in 2026 (median 54.4) and 39.2–63.5 million in 2025 (median 50.4); trimmed mean read length was 147–150 bp. Both mates of the trimmed reads were re-assessed with FastQC, and reports were aggregated per dataset with MultiQC 1.28 (Ewels et al., 2016). Per-base and per-sequence quality passed in every file of both batches; per-base sequence content warned or failed in every file, the expected signature of random-hexamer priming (Hansen et al., 2010) and deliberately not trimmed. Sequence duplication is high in all libraries, as expected for deep RNA-seq, and duplicates were not removed (Parekh et al., 2016); duplication rank within batch coincides exactly with the library-complexity ranking from the detection-sensitivity survey (Section S5), so the two independent measurements identify the same libraries. Vendor MD5 checksums were verified for every raw file (2025: 27 of 27; 2026: 21 of 21).

**Alignment.** Reads were aligned with HISAT2 2.2.1 (Kim et al., 2019; `--dta`) to the GRCm38 primary assembly with GENCODE vM25 annotation (Frankish et al., 2021), the last GENCODE release on GRCm38, supplying the annotation's 284,754 splice junctions with `--known-splicesite-infile`. The HISAT2 index has the annotation's splice sites and exons built in and no SNPs, as checked with `hisat2-inspect` (0 SNPs, 274,187 splice sites); its directory name, `genome_snp_tran`, is historical. Alignments were coordinate-sorted and indexed with SAMtools 1.18 (Danecek et al., 2021). Overall alignment was 93.8–97.8% in 2026 (median 97.1%; 84.5–92.4% of pairs aligning concordantly exactly once) and 86.7–91.9% in 2025 (median 89.3%; 72.7–83.9%). Checksums of the genome, annotation and index files are recorded with the count matrix; the MD5 checksums of the raw FASTQ files are listed for each library in Supplementary Table 13.

**Counting.** All 48 libraries were counted in a single featureCounts invocation (Liao et al., 2014; Subread 2.0.6; `-p --countReadPairs -B -C -t exon -g gene_id -O -M --primary -Q 10 -s 0`, GENCODE vM25 GTF), so that both batches were quantified by one program version with identical settings, yielding a 55,401-gene matrix of fragment counts. The matrix and the raw reads are deposited in the NCBI Gene Expression Omnibus under accession GSE348661. Because `-M` admits multi-mapping fragments and `-Q 10` then removes most of them (HISAT2 assigns low MAPQ to multi-mappers), the matrix in practice contains near-uniquely aligned fragments. Because the libraries are unstranded, a fragment in an antisense-overlapping gene pair is counted for both partners under `-O`, which is the appropriate behaviour when the originating strand is unknown but means counts at such loci are shared rather than resolved.

**Strandedness was measured rather than assumed.** In the processing run, three libraries from each batch were counted at each of the three strand settings with fragment counting enabled, and counting proceeds only if the measured setting agrees with the one recorded for the dataset. Assignment was 65.5% (2025) and 73.4% (2026) unstranded against 34–38% at either directional setting, with the two directional settings indistinguishable, which is the signature of a non-directional library. The libraries were therefore counted as unstranded. Assigned fragments were 65.1–78.4% of primary fragments per library (median 69.2%; Supplementary Table 18); the ratio assigned / (assigned + no feature) was 81–97%. An earlier processing of the same raw reads, trimmed and aligned separately for each batch and counted together in August 2026 with the same settings, gave matrices within 31 fragments per library of these (at most 26 fragments for any gene; Pearson r of $\log_2$ counts > 0.99999; GSE183742 identical), and the full analysis gives the same value for every reported result from either matrix; the only difference, a library-QC flag, is noted in Section S5. The comparison is deposited with the upstream pipeline (`docs/validation/reprocess_2026-09`).

**Gene universe and pre-filter.** Genes with a count of ten or more in at least two libraries of the fitted subset were retained for model fitting. All-zero genes were retained in the expression-status step, because a gene with no counts in any library is a valid "not expressed" call and must stay in the panel denominator; they were excluded only from the permutation control universe (Section S9).

#### S4. Gene panels and the primary-panel assignment

Three curated panels were assembled from the literature and from Gene Ontology and pathway annotation (Gene Ontology Consortium, 2023): axon guidance (92 genes), myelination (25) and vascular/lymphatic (177), 294 panel entries over 263 distinct genes. Panel lists carry Ensembl release 112 (Harrison et al., 2024) identifiers; ENSMUSG identifiers are stable between GRCm38 and GRCm39, and the pipeline verifies every panel gene against the counted gene universe on each run and asserts the expected panel sizes. Gene symbols and chromosome assignments are from Bioconductor `org.Mm.eg.db` (Carlson, 2026). The three built panels are committed with the code as master lists; every run rebuilds the panels from the curated lists and stops if the result differs from the committed lists by a single gene.

Thirty genes belong to more than one panel (27 axon guidance + vascular/lymphatic; *Gas1* axon guidance + myelination; *Notch1* myelination + vascular/lymphatic; *Sema4d* all three). Each is tested once, within one *primary* panel, so that no gene receives two adjusted P values. Assignment follows a stated precedence, axon guidance > myelination > vascular/lymphatic, on the grounds that the first two are narrow mechanism-specific curations and the third a broad annotated survey; one biological override is recorded in the configuration (*Notch1* → vascular/lymphatic, because Dll4–Notch1 signalling specifies tip and stalk identity during sprouting and its myelination role is secondary; this moves the gene into the largest, most conservative correction family). The resulting multiplicity families are 92 axon guidance, 22 myelination and 149 vascular/lymphatic genes. Per-panel tables report each gene under every panel it belongs to, carrying the same adjusted P value; totals in the text are over distinct genes.

#### S5. Library quality control; no library was excluded

Library complexity was assessed by rarefaction (multinomial subsampling to the smallest library, three replicates) and summarised as the number of detected genes and the effective number of expressed genes (exp of Shannon entropy; Hill, 1973) at matched depth. Rarefaction was used only for this comparison, never for any differential test (McMurdie & Holmes, 2014). Each library was compared with the median of its own tissue-by-age stratum, since detection differs by both; a library more than three median absolute deviations below its stratum median is flagged as a hard outlier (Leys et al., 2013), and one below 90% of the stratum median as a soft flag. No corneal library received a hard flag; six corneal libraries received soft flags, and these coincide exactly with the libraries of highest raw-read duplication within their batch. One trigeminal library lies at the hard-outlier threshold (robust z −2.96 with the reported matrix, −3.17 with the separately processed matrix of Section S3); it was not excluded.

#### S6. Expression status: the three-state rule

Each panel gene was resolved, in each group of libraries, into one of three states rather than treated as present or absent.

**Rule.** Counts were pooled across the libraries of a group (k = total fragments for the gene, L = total library size), and exact Poisson 95% limits on the pooled rate were computed from the gamma quantile function, lower = `qgamma(0.025, k)` / L × $10^6$ and upper = `qgamma(0.975, k + 1)` / L × $10^6$ (Garwood, 1936; the gamma form of Ulm, 1990). A gene is **expressed** when the lower limit exceeds $T_{HI}$ = 1 CPM, **not expressed** when the upper limit falls below $T_{LO}$ = 0.2 CPM, and **indeterminate** otherwise.

**Why pooled counts.** A replicate-based detection rule (for example, “counted in every library”) lets the least complex library govern the whole group, which is what the earlier exclusion in Section S5 had been written around. Pooling makes the criterion depend on total counts and total depth rather than on how those were divided among libraries, so groups of four and five are judged on the same footing, and the interval width follows depth in the way Poisson sampling implies. The Poisson interval describes counting uncertainty only; between-library biological variance is handled by the stability gate below and, for quantitative claims, by the negative-binomial models.

**Stability gate.** Any call that changed when any single library was removed from the pool (a leave-one-out check in the jackknife sense; Efron, 1979) was reclassified as indeterminate, so no reported status depends on which library happened to be retained. The gate only ever demotes; it moved 1.9–6.1% of panel-gene calls per group to indeterminate, most in the adult mutant groups (6.1%; adult wild type 4.9%), and its cost per group is reported (`GateCost_ByCell.csv`). Because removing one library of four perturbs the pooled depth more than one of five, the gate is slightly stricter in smaller groups; with six of seven groups at n = 5 this had no measurable effect on the results.

**Thresholds.** The band 1 / 0.2 CPM is fixed in the configuration and was not tuned to the results; it was chosen so that an “expressed” call requires the pooled rate to exceed

1 CPM with 95% confidence and a "not expressed" call requires it to fall below one-fifth of that. Its consequences are that expressed calls begin at roughly 1.3 CPM pooled and not-expressed calls end at roughly 0.1 CPM, with everything between indeterminate by construction. Sensitivity to the band and to the categorical fold threshold is reported in Section S12.

#### S7. Assessability and categorical differences

For each pairwise comparison, the two status calls place every panel gene in one of five classes, recorded in `Assessability.csv`:

1. **Test quantitatively** — expressed in both groups; the gene enters the differential model (Section S8).
2. **Categorical (absent in one group)** — expressed in one group, not expressed in the other.
3. **Categorical (interval separated)** — expressed in at least one group, and the two Poisson intervals do not overlap by at least the configured fold: bounded fold = $\max(a_{lower} / b_{upper}, b_{lower} / a_{upper}) \geq 2$. Because this quantity exceeds 1 exactly when the intervals are disjoint, non-overlap and the fold threshold are a single criterion. Non-overlap of two intervals is a conservative criterion for a difference, stricter than a direct test at the same level (Schenker & Gentleman, 2001), which suits a class that asserts a difference without fitting a model.
4. **Off in both** — not expressed in either group.
5. **Cannot assess** — the remainder, with the reason named (one group expressed but intervals overlap; or neither group confidently expressed).

Classes 2 and 3 together are the "categorical differences" of Figure 8B. Class 3 was added because the operating band leaves a wide indeterminate zone: a gene at 0.3 CPM in wild type and 20 CPM in mutant cannot be called "absent" in wild type, yet the difference is unambiguous. The class draws only from genes that could never enter the differential model, so it changes no fold change or P value anywhere; the script stops if the "test quantitatively" count ever varies with the fold threshold. Counts at minimum folds of 1, 1.5, 2, 3 and 5 are printed every run and given in `Categorical_FoldThresholdSensitivity.csv`; there is no elbow, and 2 was chosen on interpretive grounds. Genes that are categorical only at folds between 2 and 2.5 are identified in the supplementary tables so that no argument rests on them.

The comparison of transparent adult mutant with adult wild type appears in both models of Section S8 with the same libraries; categorical calls are counted once for it, giving 97 categorical calls over 55 genes across the five distinct comparisons.

#### S8. Differential expression on the testable set

**Two models, not one.** Genes expressed in both groups of a comparison were tested with DESeq2 (Love et al., 2014; median-of-ratios normalization [Anders & Huber, 2010], default dispersion estimation, Wald tests). Two models were fitted rather than one, because the developing cornea and the adult cornea undergoing opacification do not share a variance structure: the **developmental model** comprises the 29 transparent corneal libraries (both genotypes at P3/P4, P15 and adult), and the **opacity model** the 15 adult libraries (wild type, transparent mutant, opaque mutant). A single combined fit gave near-identical effect estimates (Pearson r 0.97–1.00 across contrasts) but tested adult contrasts against neonatal variability. Counts of differing genes are therefore compared within a model, never between models.

**Design.** Each model is `~ batch + male_frac + cell_group`, where `cell_group` is a single factor whose levels combine age, genotype and transparency (`P3_P4_WT`, `P3_P4_Sey`, `P15_WT`, `P15_Sey`, `Adult_WT`, `Adult_Sey_T` in the developmental model; `Adult_WT`, `Adult_Sey_T`, `Adult_Sey_O` in the opacity model). Each cell has its own mean, no additivity is assumed, an interaction is implicit, and every contrast is extracted from one fit and shares one dispersion estimate. Batch was included after fitting each model with and without the term: a genome-wide comparison found a detectable batch effect in 13–50% of expressed genes depending on age stratum (Section S2), and the two batches differ measurably in library properties, so the term is retained in every model; batch was estimable in every model with no empty batch-by-group cell. The developmental model yields *Pax6*$^{Sey\text{-}Neu/+}$ versus wild type at each age; the opacity model yields the three pairwise adult contrasts. The transparent-mutant versus wild-type adult contrast is estimable in both; it is reported from the opacity model throughout, and the developmental-model estimate is written to `HingeCheck.csv` as a consistency check (agreement on outcome 49/61 axon guidance, 14/17 myelination and 94/112 vascular/lymphatic panel entries; r of $\log_2$ fold change 0.97–1.00).

**Effect sizes.** $\text{Log}_2$ fold changes are the unshrunk maximum-likelihood estimates, with standard errors and 95% confidence intervals computed from them. Shrinkage was not applied because it would narrow the intervals on which the equivalence tests rest and make “no change” claims look stronger than the data support; `betaPrior = FALSE` is asserted at fit time.

**Three outcomes per gene.** Each gene was tested twice. The primary Wald test gives *changed* at adjusted $P < 0.05$, with no fold-change threshold. Genes not significant on the primary test were tested for *equivalence* against the composite null $|\log_2$ fold change$| \geq 0.585$ (1.5-fold) using DESeq2’s `lfcThreshold` with `altHypothesis = "lessAbs"`, whose P value is the larger of two one-sided tests (Schuirmann, 1987); genes significant on this test are reported as *no change detected*, with an effect of 1.5-fold or larger excluded. Genes significant on neither are *undetermined*, which does not distinguish absence of an effect from an effect the experiment lacked power to detect. The bound of 1.5-fold was chosen as the smallest effect the study would regard as biologically relevant; at 2-fold most non-significant genes become bounded, at 1.5-fold most remain undetermined, and the more conservative claim was preferred.

**Multiplicity.** P values were corrected by the Benjamini–Hochberg procedure (1995) within primary panel and contrast, so that each gene is corrected exactly once and that value is reported wherever the gene appears; the correction panel and the size of its family are given per gene in the supplementary tables. Independent filtering was not applied, because the tested set is already an expression-based prefilter chosen before any test was run, and filtering again on mean count would select on the data twice (Bourgon et al., 2010). Cook's-distance (Cook, 1977) outlier flagging was left at DESeq2's default; with 4–5 libraries per group DESeq2 does not replace outlying counts, so an affected gene would be reported as not testable rather than tested. No gene was set aside on this criterion in any contrast, and no gene was lost to the pre-filter after passing the status gate (`CooksFiltering.csv`, `TestabilityAudit.csv`).

**Adult omnibus and interaction tests.** For the opacity model, a likelihood-ratio test of the full model against one lacking `cell_group` identifies genes differing among the three adult groups (`AdultOmnibusLRT.csv`). In the developmental model, whether the genotype effect itself differs between two ages was tested directly as a difference of differences from the cell-group parameterisation (three interaction contrasts: P15 vs P3/P4, adult vs P15, adult vs P3/P4), because comparing two separately significant pairwise contrasts is not that test (Gelman & Stern, 2006). An interaction was evaluated only where both component effects were estimable, and the same three-outcome scheme was applied. About 7% of interaction tests (33 of 483: 26 significant and 7 bounded as equivalent) were resolved in either direction (`InteractionResults.csv`), so the manuscript describes which contrasts reached significance rather than inferring times of onset, and does not claim constancy of a genotype effect across ages.

#### S9. Program-level test

Differences between two groups in the number of confidently expressed, and the number of confidently not-expressed, panel genes were tested against an abundance-matched null. The null is competitive (Goeman & Bühlmann, 2007): the panel is compared with sets of non-panel genes drawn from the same libraries. The control universe comprises autosomal, non-panel genes detected in at least one library (sex-linked and mitochondrial genes excluded because pool sex composition varies). Genes were stratified into 20 abundance bins by quantiles of mean CPM over the two groups; ties at the detection floor collapse bins as needed and the number used is recorded. For each of 20,000 permutations a control set of the panel's size and abundance profile was drawn and the same two status counts computed, giving the null distribution of the observed difference; matching on abundance is necessary because detection probability depends on abundance and an unmatched control would measure that instead of the effect (Young et al., 2010). P values are (1 + r) / (n + 1) (Phipson & Smyth, 2010), so a value of $5 \times 10^{-5}$ means only "smaller than the resolution of 20,000 permutations" and such values are not ranked against one another. Both metrics for all five comparisons and three panels (30 tests) were corrected jointly by Benjamini–Hochberg. Because panel genes are co-regulated, competitive tests are inflated relative to their nominal level (Wu & Smyth, 2012); the program-level P values order the panels and contrasts rather than being exact, and this is stated in Limitations. The null's 2.5–97.5% interval gives each test's detection limit (the smallest excess that would have reached significance), which is reported alongside every negative result so that a negative can be read as bounded rather than as underpowered.

#### S10. Developmental analysis and Figure 7

Within each genotype separately, the transparent corneal libraries at P3/P4, P15 and adult were analyzed on two levels.

**Status crossings.** The three-state rule of Section S6 was applied to each age group, and a gene was recorded as turning *off* between two ages when it was expressed at the earlier and not expressed at the later age, and *on* for the reverse, for the two adjacent intervals (P3/P4→P15, P15→adult) and for the whole span (P3/P4→adult). The whole-span column is a direct comparison of the first and last ages, not a union of the adjacent intervals: a gene can cross there without crossing in either adjacent interval (when it moves through the indeterminate zone in two steps) or cross in an adjacent interval without crossing there (a transient change, as for *Mpz*). Figure 7 shows every gene with at least one crossing in either genotype, grouped into three bands by whether it crosses in wild type only, in both genotypes, or in mutant only, ordered within band by panel and by the earliest wild-type crossing.

**Why status level.** A gene falling from 50 to 10 CPM changes quantitatively but not in state; a gene falling from 3 CPM to below detection changes state. The imaging questions are state questions (are vessels present; are nerves present), and the paper's central developmental claim — that the normal cornea silences the vascular/lymphatic identity program between P15 and adulthood and the mutant cornea does not — is a claim about state. Figure 7 is the developmental account; it is not the statistical test of the genotype difference. That test is the adult genotype contrast (Section S8) and the program-level comparison (Section S9), reported in Figure 8, and the Results cite those wherever a genotype difference is asserted.

**What Figure 7 cannot show.** A blank cell means no crossing was detected, not that a difference was excluded. The interval in which a crossing occurs is not compared between genotypes (Section S8, interaction tests). Because the tissue is whole cornea, a gene "expressed" may reflect the presence of a cell type rather than regulation within corneal cells. Row-band membership is specific to the 1 / 0.2 CPM band (Section S12). The ungated within-genotype age contrasts for every panel gene, with the Figure 7 crossings marked, are shown as heatmaps in Supplementary Figure 4 (Section S12).

**Quantitative change within genotype.** In parallel, within each genotype the transparent libraries were modelled across the three ages (`~ batch + male_frac + age_group`) with pairwise contrasts between adjacent ages and between the first and last. A gene entered a comparison only where confidently expressed at both ages and was classified as changed, no change detected or undetermined exactly as in Section S8, with Benjamini–Hochberg correction within primary panel, genotype and interval. A shape over the three ages was then derived mechanically from the two adjacent-interval outcomes rather than imposed on them (`Developmental_Trajectories.csv`): rises or falls across both intervals; rises then falls (peak at P15) or falls then rises (trough at P15); up or down to P15 only, or after P15 only, when the other interval is bounded; no change detected across either interval; not assessable when the gene was not expressed at both ages of an interval; and not resolved otherwise. The whole-span contrast is reported but never used to infer a shape. Shapes are threshold summaries and are not compared between genotypes. Positive evidence of developmental stability at the 1.5-fold level was obtainable for very few genes, because a typical undetermined interval leaves 0.84–1.01 $\log_2$ units open against a bound of 0.585.

#### S11. Figure 8

Figure 8A plots, for each panel and comparison, the excess of confidently expressed and confidently not-expressed genes over the abundance-matched null (Section S9), with its 95% null interval, and marks the tests that survive joint correction. Figure 8B lists every categorical difference (Section S7) for the four comparisons in which genotype and opacity are separable (45 status changes in 41 genes), with direction and bounded fold, arranged by panel and comparison; all five comparisons together (97 changes in 55 genes) are in Supplementary Figure 7. Unconfirmed calls are drawn open (Section S12). The confounded comparison (opaque mutant versus wild type, in which genotype and opacity are not separable) is shown as Supplementary Figure 7 rather than in the main figure.

#### S12. Concordance with a replicate-aware analysis, and threshold sensitivity

The three-state framework was chosen for interpretability, not because it was expected to find what a standard analysis would miss. To establish that its status-level results are a subset of what a standard analysis reports, and to identify any that are not, every status crossing and categorical difference was re-examined without the status gate (`07_concordance.R`).

**Ungated models.** The DESeq2 models of Sections S8 and S10 were applied to *all* panel genes passing the count pre-filter, not only those expressed in both groups, and Benjamini–Hochberg correction was applied over that larger family within primary panel and contrast. For each Figure 7 crossing, the corresponding within-genotype contrast was looked up and classed as *confirmed* (adjusted $P < 0.05$, same direction as the crossing), *not significant*, *opposite* (significant in the other direction) or *not in model*; for each categorical difference, the corresponding genotype contrast was classed the same way. A reverse check listed genes significant at ≥ 2-fold under the ungated model that received no status-level call, with the reason (`Concordance_AgeAxis_StandardSignificant.csv`; the “cannot assess” class of Section S7).

**Second engine.** To ensure the comparison did not rest on one implementation, the identical design was fitted with edgeR quasi-likelihood (TMM normalization, `estimateDisp`, `glmQLFit`, `glmQLFTest`; Robinson & Oshlack, 2010; Lund et al., 2012; Chen et al., 2016; Chen et al., 2025) on the same samples and pre-filtered genes, and each categorical difference was classed against it as well. No reported number derives from edgeR.

**Result and its use in the figures.** Of the 144 status crossings in Figure 7, 126 (88%) were significant in the same direction under the ungated DESeq2 model and none was significant in the opposite direction; of the 97 categorical differences, 86 (89%) were confirmed by DESeq2 and 91 by edgeR, with edgeR and DESeq2 agreeing on 92–97% of panel genes per contrast. Every "absent in one group" call was confirmed; all unconfirmed calls are interval-separated with bounded folds of 7.1 or less. Unconfirmed crossings and differences are drawn as open symbols in Figures 7 and 8B, named in the captions, and are not quoted individually in the text; the confirmed set carries every narrative claim. A diagnostic that accompanies each call, the share of the pooled count contributed by the single largest library (`top_library_share`), is enriched among unconfirmed calls (11 of 18 crossings above 0.5, against 48 of 126 confirmed) and identifies the cases in which one library carried a pooled rise that the replicate model does not support (for example *Mmp3* in adult wild type, share 0.90).

**Threshold sensitivity.** The status band and categorical fold were varied over a 3 × 3 grid ($T_{HI}$ / $T_{LO}$ = 0.5/0.1, 1/0.2, 2/0.4 CPM; minimum fold 1.5, 2, 3), recomputing the crossings and the Figure 7 row bands at each setting with a local implementation of the rule (`Concordance_ThresholdStability.csv`). Two acceptance tests precede the sweep: the local rule must reproduce the calls of `03_expression_status.R` exactly in every group, and the sweep at the configured settings must reproduce the crossings of `06_developmental.R` exactly; the script stops otherwise. At the 1 / 0.2 CPM band, 51–58 of the 58 genes in Figure 7 keep their row band across minimum folds 1.5–3; changing the band itself leaves only 25–35 of the 58 genes in their row band, as expected since the band defines which genes are near the detection limit. The captions therefore state that band membership is specific to the configured band.

**Sample-level diagnostics.** P-value histograms for every contrast of both engines (`QC_PValueHistograms.png`), an inter-library Spearman correlation heatmap on log CPM (`QC_SampleCorrelation.png`), and the per-gene standard-model heatmaps for the developmental contrasts (Supplementary Figure 4A, axon guidance and myelination; 4B, vascular/lymphatic) are written by the same script.

#### S13. Cross-dataset replication (GSE183742)

**Dataset and processing.** GSE183742 (Duncan laboratory; Krishnan et al., 2024) comprises six adult corneal libraries, three wild-type and three *Pax6*$^{tm1Pgr/+}$, deposited as raw reads. It was trimmed and aligned in the same processing run as the present libraries, with the same reference (GRCm38, GENCODE vM25), index, program versions and settings (Section S3), and counted as a separate study. Strandedness was measured as in Section S3: assignment at the forward setting collapsed to about 2% while the reverse setting nearly matched unstranded, so these libraries are reverse-stranded and were counted at `-s 2`, the opposite of the present libraries. Overall alignment was 51–55%; the unaligned fraction is dominated by 45S pre-rRNA internal and external transcribed spacer sequence (BLAST; Altschul et al., 1990), which lies in the rDNA repeat array collapsed in the primary assembly and so has nowhere to map. Retention of this non-polyadenylated RNA indicates a ribosomal-depletion rather than poly-A library, a second chemistry difference from the present study. Read pairing, read quality and chemistry were ruled out as causes (R1 alone aligned at the same rate as the pair). After counting, 8.6–11.2 million fragments were assigned per library (assigned / (assigned + no feature) 73–80%), roughly a quarter of the present depth. Transparency of the mutant corneas is not reported in the deposit, so the mutant-versus-wild-type comparison in that dataset conflates genotype with any opacity present. Because the allele, laboratory, chemistry and depth all differ, the dataset was treated as a separate study and never pooled with the present one.

**Tier 1: wild-type expression status.** The three-state rule (Section S6, including the stability gate and the full-library denominator) was implemented once, in the replication script, and required to reproduce the present study's adult wild-type calls for all 263 panel genes exactly before any comparison (acceptance test; 0 disagreements). It was then applied to the GSE183742 wild-type libraries and the calls compared. Genes indeterminate in either dataset are reported separately and never counted as agreement or disagreement, so the unresolved fraction is set largely by the shallower dataset. Agreement is reported as the proportion and as Cohen's κ (Cohen, 1960), and is compared with an abundance-matched null: 2,000 draws of non-panel gene sets matched to the panel's abundance profile in 20 quantile bins (ties at the detection floor collapsed), scored identically. Two adult corneal datasets agree on most genes by construction, so the null, not the raw percentage, is the interpretable quantity.

**Tier 2: direction of the genotype effect.** For genes that were categorical (Section S7, bounded fold ≥ 2) in both datasets, the direction of the mutant-versus-wild-type difference was compared with the present opaque-mutant-versus-wild-type contrast, the comparison that matches the unknown opacity status. A binomial test against 0.5 is not reported: with both datasets dominated by genes higher in mutant, independent calls would agree on most genes by their margins alone (86% for the observed margins), and the genes are co-regulated members of three programs, not independent draws. The script instead reports the 2 × 2 direction table, agreement expected from the margins, Cohen's κ, a one-sided Fisher exact test with both margins fixed, and an abundance-matched permutation null in which non-panel genes are scored by the identical rule in both datasets; because "categorical in both" selects on effect size, the null draw is scaled (calibrated over multipliers 1–32) so that its median number of resolved genes matches the panel's. The permutation null is the quantity to interpret, because it is the only one of the three that places comparable co-regulation and the same selection in the reference.

**Results.** Tier 1: 155 of 263 panel genes resolved in both datasets, 152 agreeing (98.1%, κ = 0.79); null median 96.8% (2.5–97.5%: 93.9–99.4), P = 0.19. The three disagreements all run one way (not expressed here, expressed in GSE183742). Tier 2: 47 genes categorical in both, 46 agreeing in direction (κ = 0.85; Fisher $P = 2.5 \times 10^{-4}$); size-matched permutation null median 97.9% (92.6–100), P = 0.53. In neither tier does the panel exceed abundance-matched background: the calls transfer to an independent dataset with a different allele, laboratory and chemistry, and they transfer as well as comparable genes do. The four-orders-of-magnitude gap between the Fisher and permutation P values is the size of the dependence and selection effects, and is why the permutation value is the one reported. The single direction reversal, *Dscam*, is confident in both datasets (bounded fold 7.0 here, 6.1 there).

**Dissection-margin check.** The three Tier 1 disagreements (*Tek*, *Sema3g*, *Epha5*) and *Dscam* are genes expressed in limbal, conjunctival or angle tissue, and the dissection margin of GSE183742 is not reported. A fixed set of markers was therefore compared between datasets: central-cornea controls (*Krt12*, *Aldh3a1*, *Kera*, *Angptl7*); limbal and conjunctival epithelium (*Krt15*, *Krt19*, *Krt13*, *Krt4*, *Muc5ac*); trabecular meshwork and angle (*Myoc*, *Mgp*); pigmented tissue (*Tyr*, *Tyrp1*, *Dct*, *Mlana*); vascular endothelium (*Vwf*, *Cldn5*); lens (*Cryaa*, *Cryba1*, *Crybb2*, *Crygd*, *Mip*, *Bfsp2*); retina and RPE (*Rho*, *Gnat1*, *Sag*, *Pde6b*, *Rcvrn*, *Crx*, *Nrl*, *Rpe65*); and ciliary body (*Optc*). The list and its thresholds are shared with the off-target tissue screen of Supplementary Table 13; *Muc4*, absent from the count matrix, was dropped. Status and bounded fold were computed for each marker in each wild-type group. Eleven of the 28 peripheral markers were expressed in the GSE183742 wild-type corneas and not here at twofold or more, and none in the other direction, consistent with inclusion of peripheral tissue in that dissection; the same comparison shows the stromal controls *Kera* and *Angptl7* 104- and 37-fold lower there, consistent with an epithelium-enriched preparation, and *Rho* at 261 CPM, indicating retinal contamination. These observations explain the disagreements without invoking the allele and are stated as such; the dissection boundary itself is not reported and cannot be verified.

#### S14. Trigeminal ganglion pilot

Fourteen trigeminal-ganglion libraries, two per group (wild-type and transparent-mutant ganglia at P3/P4, P15 and adult; opaque-mutant adults), were analyzed by `10_trigeminal.R`. Two libraries per group are below the minimum of three at which the three-state rule makes calls (Section S6), so the ganglia were analyzed by DESeq2 alone, with the corneal conventions wherever they carry over: the same pre-filter; a single factor combining age, genotype and transparency; unshrunk $\log_2$ fold changes with 95% confidence intervals; the same equivalence bound (±0.585 $\log_2$); and Benjamini–Hochberg correction within primary panel and contrast, without independent filtering. A developmental model (wild-type and transparent-mutant ganglia at the three ages; n = 12) compared genotypes within each age, and an adult model (n = 6) compared the three adult groups pairwise and by an omnibus likelihood-ratio test. Two departures from the corneal models are forced by the design. All trigeminal libraries are from the 2025 batch, so no batch term can be fitted; and the male read fraction was not included as a covariate, because with two libraries per group it would consume most of the residual degrees of freedom. Each library's male fraction is reported instead (`Trigeminal_SampleTable.csv`). As a positive control for sensitivity, a likelihood-ratio test for age was fitted on the wild-type ganglia alone (n = 6) over all genes passing the pre-filter.

No panel gene differed between genotypes in transparent tissue at any age (smallest adjusted P: P3/P4 0.30, P15 0.11, adult 0.071). Of the 745 gene-by-comparison tests in these three comparisons, 537 excluded a change of 1.5-fold or more and 208 were undetermined. Differences arose only in comparisons involving opaque animals, and all were higher in opaque ganglia: eleven genes against wild type (*Flrt3* 1.66-fold, *Egr2* 1.40, *Reln* 1.36, *Slit2* 1.32, *Mag* 1.28, *Sema5a* 1.26, *Erbb3* 1.23, *Adgrl2* 1.20, *Zeb2* 1.17, *Sema3b* 1.14, *Mbp* 1.10) and three against transparent mutants (*Egr2* 1.83, *Timp3* 1.19, *Mbp* 1.16); the adult omnibus test identified ten. The positive control found 11,468 of 20,287 tested genes differentially expressed with age.

Because *Mbp* and *Mag* are made by central oligodendrocytes as well as by Schwann cells, carry-over of nerve root or brainstem at dissection could raise them for a reason unrelated to genotype. Every library's expression of central markers (*Mog*, *Mobp*, *Olig2*, *Gfap*) and of myelinating Schwann-cell markers (*Prx*, *Mpz*, *Pmp22*, *Egr2*) is therefore reported (`Trigeminal_TissueMarkers.csv`). Central carry-over varied between libraries, most pronounced in one P15 wild-type library (*Mog* 14, *Mobp* 81, *Gfap* 89 CPM) and present to a lesser degree in wild-type, transparent-mutant and opaque-mutant libraries alike. Because it was not specific to any group, and small relative to the expression of the myelination genes concerned, it does not account for the differences in opaque ganglia. *Egr2*, essentially specific to myelinating Schwann cells, was higher in both opaque adult libraries (35 and 45 CPM) than in all four other adult libraries (19–30 CPM).

#### S15. Reproducibility and software

The pipeline is fourteen numbered R scripts run in order by `run_all.R` in one R session (`00_config.R`, `00b_panel_universes.R`, `01_load.R`, `01b_qc_library_depth.R`, `01c_contamination_indices.R`, `02_shared_gene_assignment.R`, `03_expression_status.R`, `04_differential.R`, `05_program_level.R`, `06_developmental.R`, `07_concordance.R`, `08_figures.R`, `09_cross_dataset_replication.R`, `10_trigeminal.R`). Every parameter (thresholds, α, fold, equivalence bound, permutation count, seed, covariate switches, exclusion lists, contrast table) is defined once in `00_config.R`, overridable by a `PAX6_*` environment variable, and printed at the start of each run; the sensitivity runs described above were made this way, without editing code. The driver stops at the first failure, refuses to run if a numbered script is present but not listed, and writes `RunProvenance.txt` recording the input files, resolved settings, R and package versions and timings; a copy from the run that produced the submitted figures is deposited with the code. Each figure is written with a source-data CSV, and every caption number is written to `Figure_CaptionNumbers.txt` by the figure script rather than transcribed. Two acceptance tests are built into the pipeline (Sections S12 and S13) and the categorical class carries an invariant check (Section S7). The released code was verified by a complete run from a fresh clone of the repository, which reproduced the input checksums and every reported number.

The results reported were produced with R 4.6.1 (R Core Team, 2026), DESeq2 1.52.0 (Love et al., 2014), edgeR 4.10.5 (Chen et al., 2025), limma 3.68.5 (Ritchie et al., 2015), ggplot2 4.0.3 (Wickham, 2016), patchwork 1.3.2 (Pedersen, 2025), org.Mm.eg.db 3.23.0 (Carlson, 2026), AnnotationDbi 1.74.0 (Pagès et al., 2026; Bioconductor 3.23,

Huber et al., 2015) and data.table 1.18.6.1 (Barrett et al., 2026) on macOS 26.7 (run of 13 October 2026 with release v1.0.4 on the count matrix as deposited at GEO; `docs/RunProvenance_v1.0.4_2026-10-13.txt`). The full session information is deposited with the code. Upstream processing of all 54 libraries (run `reprocess_2026-09`, pipeline commit `9182fd5`, 26 September 2026) used FastQC 0.12.1, Trimmomatic 0.39, MultiQC 1.28, HISAT2 2.2.1, SAMtools 1.18 and Subread 2.0.6, the versions each tool reported at run time. The upstream SLURM scripts, the dataset, genome-build, module and parameter manifests (`config/`), and the run's provenance record (tool versions, reference and raw-FASTQ checksums, per-library QC and measured strandedness) are in a separate repository shared by all analyses of these libraries (https://github.com/Lauderdale-Lab/pax6-cornea-trigeminal-rnaseq-upstream; release v1.0.0, archived at Zenodo, https://doi.org/10.5281/zenodo.22998976). Repeated full runs reproduce every reported number exactly. Code is available at https://github.com/Lauderdale-Lab/pax6-cornea-nerve-vascular-transparency (release v1.0.4, commit `43597c4`) and archived at Zenodo (v1.0.4, https://doi.org/10.5281/zenodo.23347047; all versions, https://doi.org/10.5281/zenodo.22965379) under the MIT licence. The v1.0.4 run reproduced every file of the v1.0.3 run. The complete output files of the reported run are archived at Zenodo (https://doi.org/10.5281/zenodo.23349739), and the library metadata and processing quality-control tables (Supplementary Tables 13 and 18) at https://doi.org/10.5281/zenodo.23350548; raw reads and the count matrix are deposited at GEO (GSE348661); GSE183742 is cited by its accession and was not re-deposited.

Supplementary Tables. The pipeline output files behind Supplementary Tables 5–12, 14–17, 19 and 20 are named in the description of each table and archived at Zenodo (see above). By section: S3, Supplementary Tables 13 and 18; S4, Table 15; S5, Tables 13 and 20; S6 and S7, Tables 10 and 11; S8, Table 16; S9, Table 17; S10, Tables 5, 6, 8 and 19; S11, Tables 11 and 19; S12, Tables 7 and 9; S13, Table 14; S14, Table 12. Supplementary Table 13 is the authoritative record of pool composition, age, sex and raw-file checksums for the 48 libraries of this study.

**A**

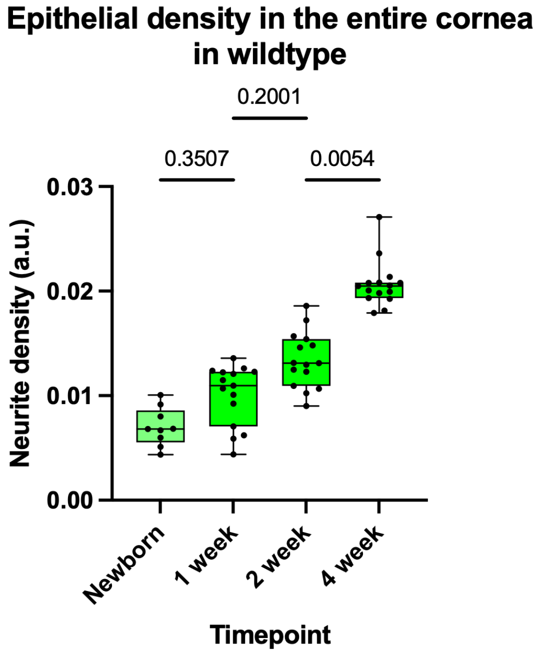
Epithelial density in the entire cornea
in wildtype
0.2001
0.3507
0.0054
0.03
0.02
0.01
0.00
Neurite density (a.u.)
Newborn
1 week
2 week
4 week
Timepoint

**B**

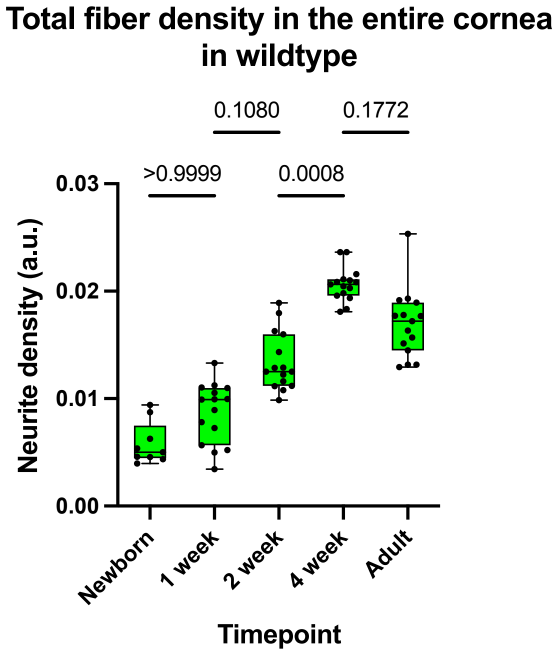
Total fiber density in the entire cornea
in wildtype
0.1080
0.1772
>0.9999
0.0008
0.03
0.02
0.01
0.00
Neurite density (a.u.)
Newborn
1 week
2 week
4 week
Adult
Timepoint

**C**

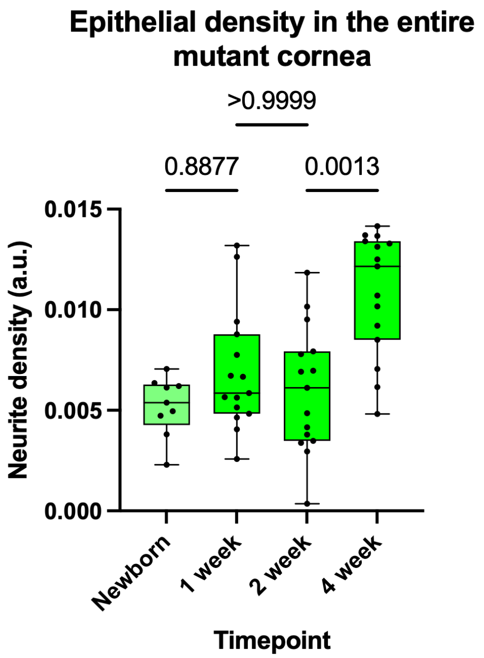

Epithelial density in the entire mutant cornea
>0.9999
0.8877
0.0013
0.015
0.010
0.005
0.000
Neurite density (a.u.)
Newborn
1 week
2 week
4 week
Timepoint

**D**

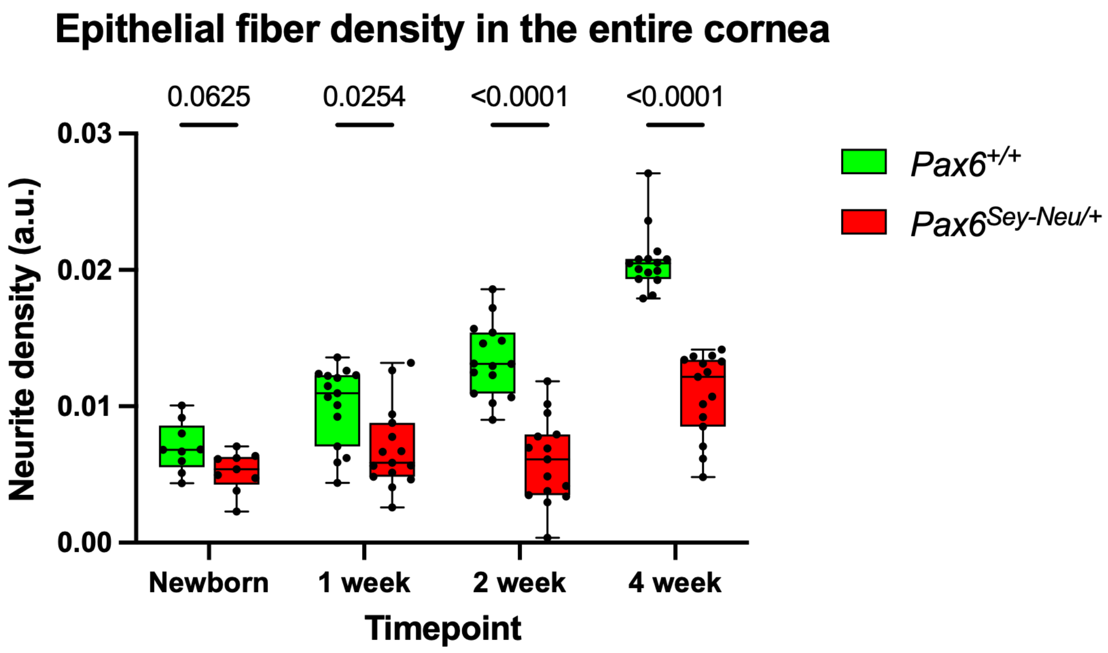

Epithelial fiber density in the entire cornea
0.0625
0.0254
<0.0001
<0.0001
0.03
0.02
0.01
0.00
Neurite density (a.u.)
Newborn
1 week
2 week
4 week
Timepoint
Pax6+/+
Pax6Sey-Neu/+

E

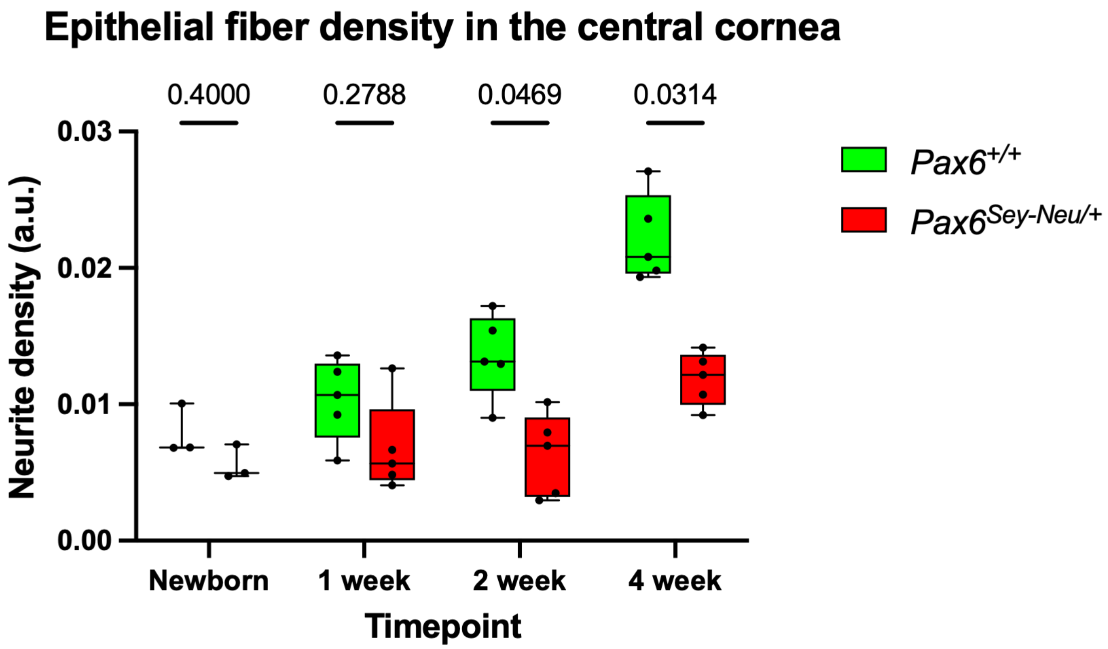
Epithelial fiber density in the central cornea
0.4000
0.2788
0.0469
0.0314
0.03
0.02
0.01
0.00
Neurite density (a.u.)
Newborn
1 week
2 week
4 week
Timepoint
Pax6+/+
Pax6Sey-Neu/+

F

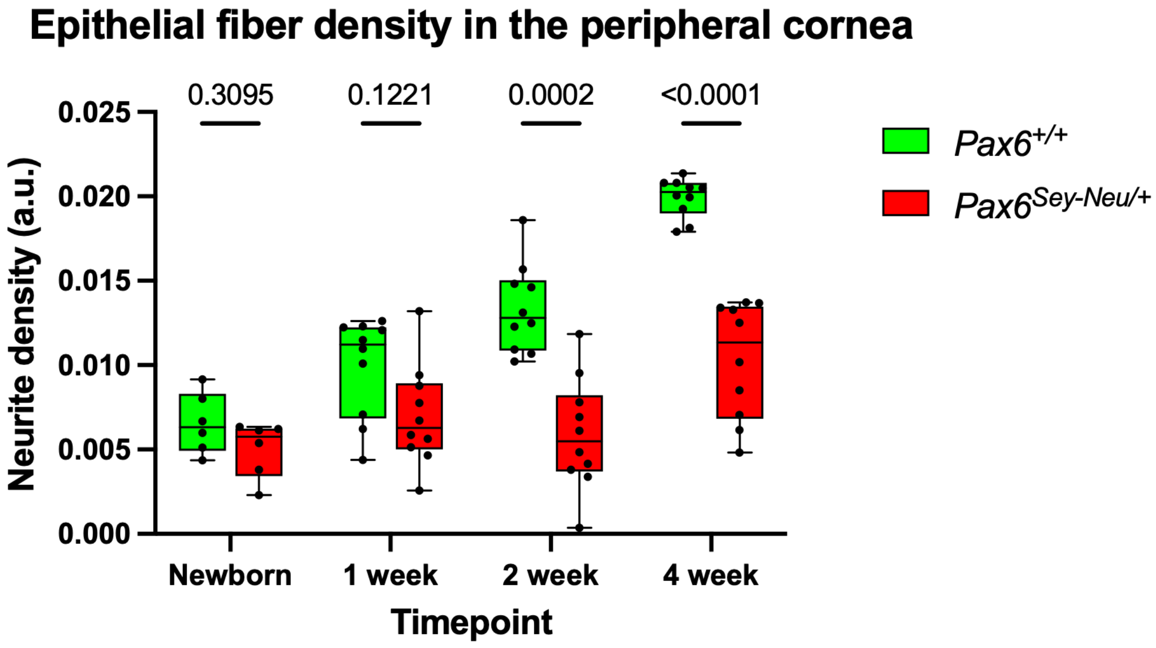
Epithelial fiber density in the peripheral cornea
0.3095
0.1221
0.0002
<0.0001
0.025
0.020
0.015
0.010
0.005
0.000
Neurite density (a.u.)
Newborn
1 week
2 week
4 week
Timepoint
Pax6+/+
Pax6Sey-Neu/+

**G**

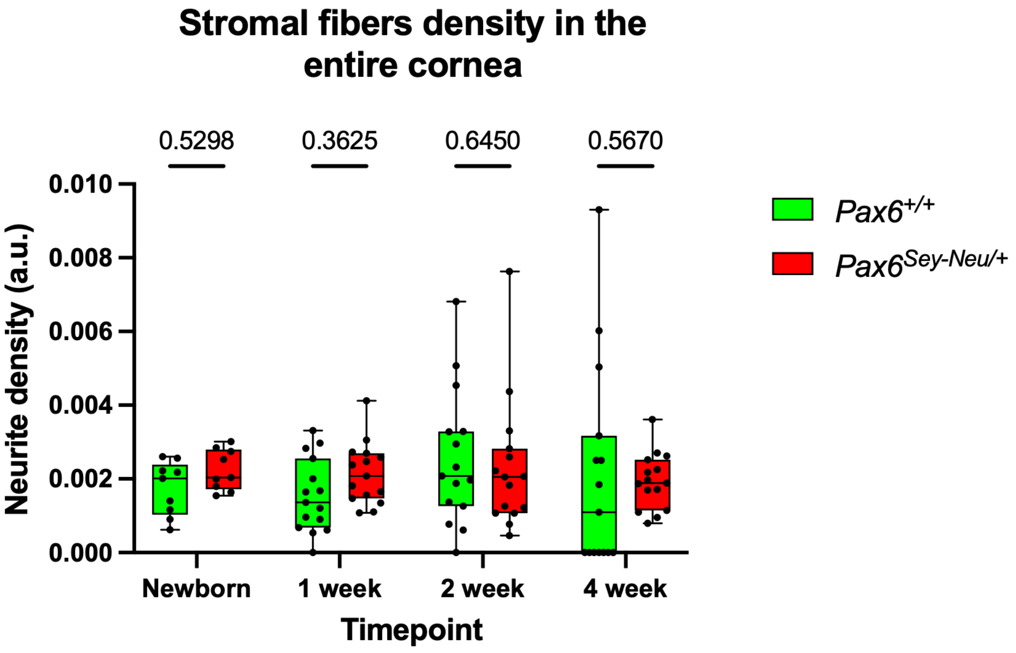
Stromal fibers density in the entire cornea
0.5298
0.3625
0.6450
0.5670
0.010
0.008
0.006
0.004
0.002
0.000
Neurite density (a.u.)
Newborn
1 week
2 week
4 week
Timepoint
Pax6+/+
Pax6Sey-Neu/+

**H**

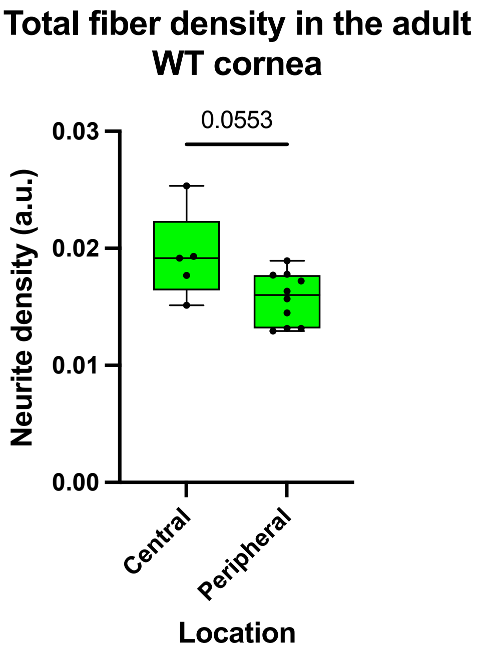
Total fiber density in the adult WT cornea
0.0553
0.03
0.02
0.01
0.00
Neurite density (a.u.)
Central
Peripheral
Location

I

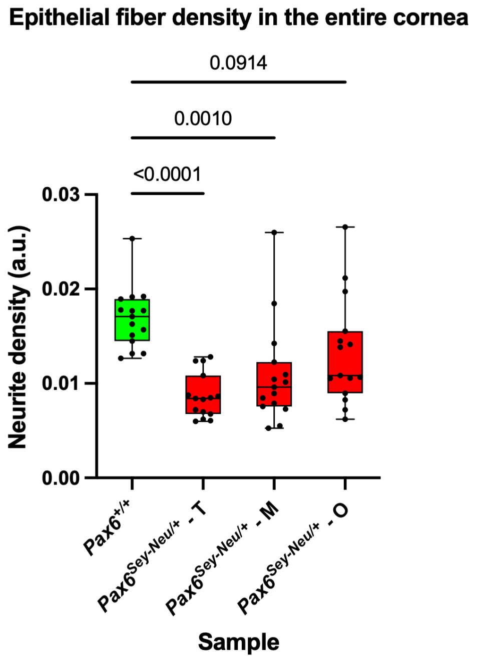
Epithelial fiber density in the entire cornea
0.0914
0.0010
<0.0001
0.03
0.02
0.01
0.00
Neurite density (a.u.)
Pax6+/+
Pax6Sey-Neu/+ - T
Pax6Sey-Neu/+ - M
Pax6Sey-Neu/+ - O
Sample

J

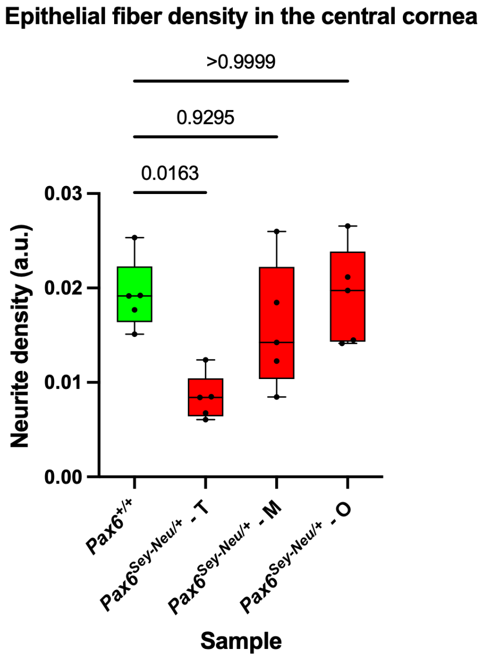
Epithelial fiber density in the central cornea
>0.9999
0.9295
0.0163
0.03
0.02
0.01
0.00
Neurite density (a.u.)
Pax6+/+
Pax6Sey-Neu/+ - T
Pax6Sey-Neu/+ - M
Pax6Sey-Neu/+ - O
Sample

K

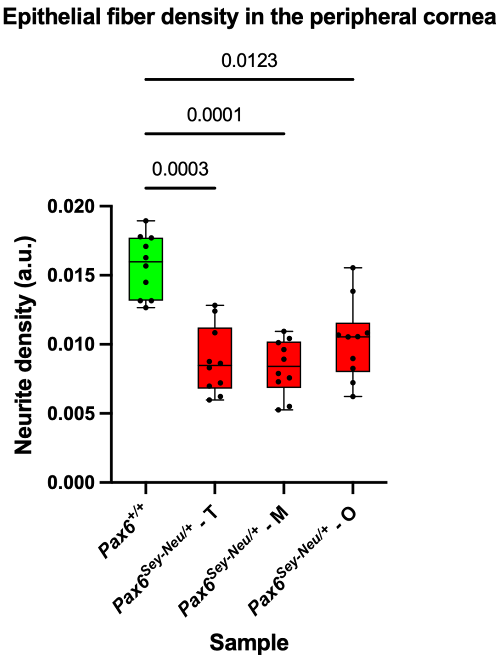

**Supplementary Figure 1. Quantification of TuJ1-labeled corneal nerve density during development and adulthood.** (A) Epithelial nerve-fiber density across postnatal development in *Pax6*$^{+/+}$ corneas. (B) Total nerve-fiber density across postnatal development in *Pax6*$^{+/+}$ corneas. (C) Epithelial nerve-fiber density across postnatal development in *Pax6*$^{Sey\text{-}Neu/+}$ corneas. (D–G) Genotype comparisons during postnatal development. Quantification of epithelial nerve-fiber density across the entire cornea (D), within a central ROI positioned inside the central 50% of corneal area (E), within two equal-area peripheral ROIs positioned outside the central zone (F), and stromal nerve-fiber density across the entire cornea (G). Peripheral ROIs were permitted up to 20% overlap with the central zone to avoid inclusion of limbal fibers. (H–K) Adult nerve-fiber density measurements. Quantification of total nerve-fiber density across the entire cornea (H), epithelial nerve-fiber density across the entire cornea (I), within a central ROI positioned inside the central 50% of corneal area (J), and within two equal-area peripheral ROIs positioned outside the central zone (K). Peripheral ROIs were permitted up to 20% overlap with the central zone to avoid inclusion of limbal fibers. For all panels, fiber density was quantified as thresholded particle counts normalized to ROI area and is expressed in arbitrary units (a.u.; see Methods). Each point represents one ROI measurement (three non-overlapping ROIs per cornea: one central and two peripheral; see Methods and Supplementary Table 1). Boxes indicate the median and interquartile range; whiskers indicate the minimum and maximum values.

Statistical analyses: (A–C) Kruskal-Wallis tests followed by Dunn's multiple-comparisons procedure. (D–G) Multiple two-tailed Mann-Whitney tests with Holm-Šídák correction for multiple comparisons. (H) Two-tailed Mann-Whitney test. (I–K) Kruskal-Wallis tests followed by Dunn's multiple-comparisons procedure. Exact or adjusted P values are shown in the graphs.

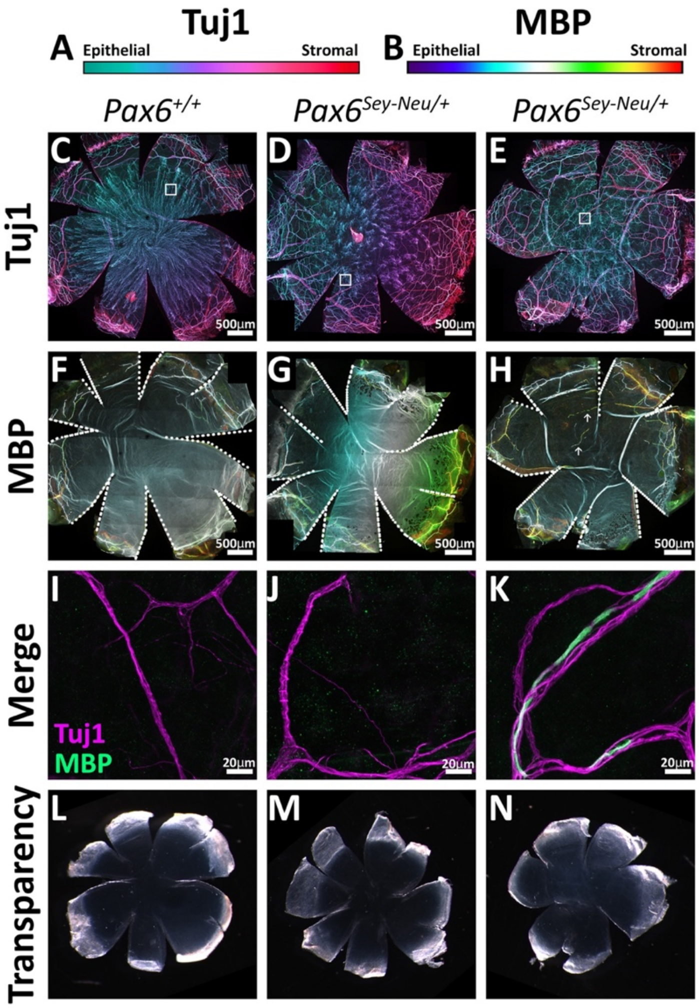

**Supplementary Figure 2. Altered nerve architecture and MBP labeling in a 2-month-old *Pax6$^{Sey\text{-}Neu/+}$* cornea.** (A) Depth-encoded color scale for TuJ1-labeled fibers, from superficial/epithelial (green) to deep/stromal (red). (B) Depth-encoded color scale for MBP immunolabeling, using the same depth convention. Columns show Pax6+/+ (C, F, I, L), a transparent Pax6Sey-Neu/+ cornea (D, G, J, M), and a Pax6Sey-Neu/+ cornea containing an opaque region (E, H, K, N). (C–E) Maximum-intensity projections of TuJ1-labeled nerve fibers across the entire cornea. White boxes indicate the location of the corresponding higher-magnification fields. (F–H) Maximum-intensity projections of selected optical sections containing MBP immunolabeling. (I–K) Higher-magnification merged images showing TuJ1 (magenta) and MBP (green) immunolabeling over the same Z range. (L–N) Backscatter images of the same corneas, illustrating transparency phenotypes. Relative scales apply. All fluorescence panels are maximum-intensity projections of confocal Z-stacks. Scale bars as indicated. Images are representative of the group.

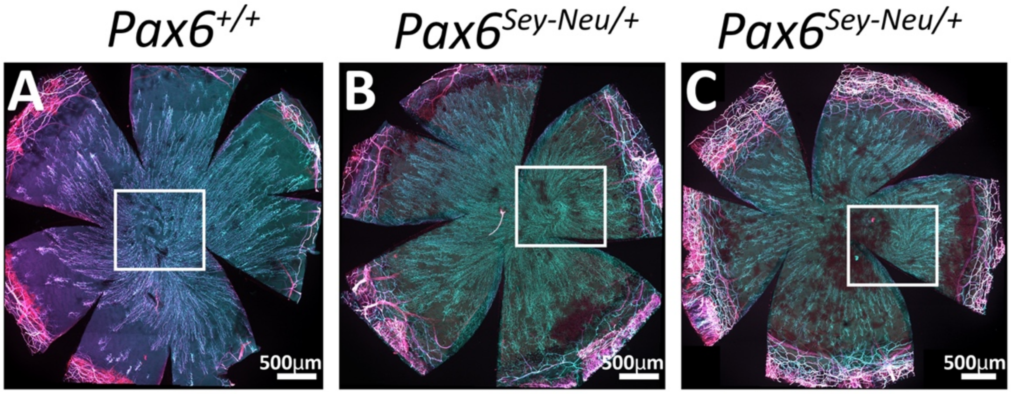

**Supplementary Figure 3. Residual subbasal whorl-like organization in a subset of adult *Pax6$^{Sey\text{-}Neu/+}$* corneas**. (A) *Pax6$^{+/+}$* cornea showing the characteristic centrally converging subbasal nerve whorl. B) *Pax6$^{Sey\text{-}Neu/+}$* cornea containing long subbasal nerve fibers with partial preservation of whorl-like organization. (C) *Pax6$^{Sey\text{-}Neu/+}$* cornea containing a localized cluster of subbasal fibers exhibiting whorl-like organization (boxed region). All panels are maximum-intensity projections of TuJ1-immunolabeled corneal whole mounts. Images illustrate the heterogeneity of nerve architecture observed among adult *Pax6$^{Sey\text{-}Neu/+}$* corneas and demonstrate that localized whorl-like organization can persist despite the broader disruption of corneal innervation described in the main figures. Scale bars as indicated. Corneas were fixed in 2% PFA prior to indirect immunolabeling (see Methods).

**A.**

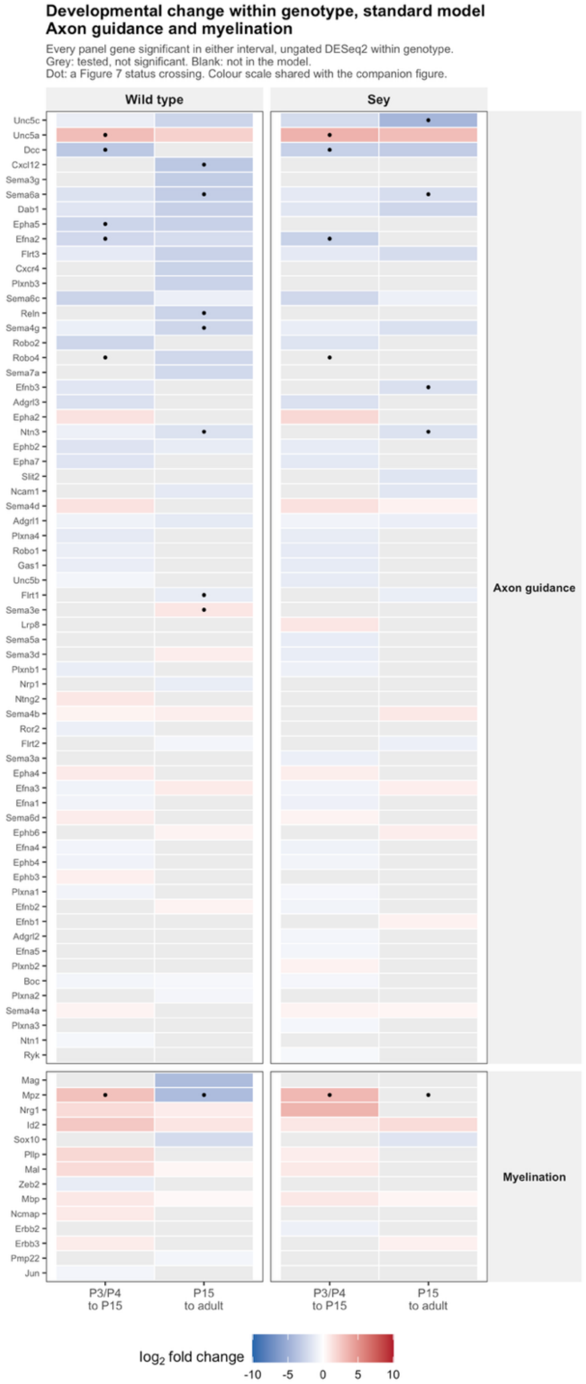
Developmental change within genotype, standard model
Axon guidance and myelination
Every panel gene significant in either interval, ungated DESeq2 within genotype.
Grey: tested, not significant. Blank: not in the model.
Dot: a Figure 7 status crossing. Colour scale shared with the companion figure.
Wild type
Sey
Unc5c
Unc5a
Dcc
Cxcl12
Sema3g
Sema6a
Dab1
Epha5
Efna2
Flrt3
Cxcr4
Plxnb3
Sema6c
Reln
Sema4g
Robo2
Robo4
Sema7a
Efnb3
Adgrl3
Epha2
Ntn3
Ephb2
Epha7
Slit2
Ncam1
Sema4d
Adgrl1
Plxna4
Robo1
Gas1
Unc5b
Flrt1
Sema3e
Lrp8
Sema5a
Sema3d
Plxnb1
Nrp1
Ntng2
Sema4b
Ror2
Flrt2
Sema3a
Epha4
Efna3
Efna1
Sema6d
Ephb6
Efna4
Ephb4
Ephb3
Plxna1
Efnb2
Efnb1
Adgrl2
Efna5
Plxnb2
Boc
Plxna2
Sema4a
Plxna3
Ntn1
Ryk
Axon guidance
Mag
Mpz
Nrg1
Id2
Sox10
Plp
Mal
Zeb2
Mbp
Ncmap
Erbb2
Erbb3
Pmp22
Jun
Myelination
P3/P4 to P15
P15 to adult
P3/P4 to P15
P15 to adult
log2 fold change
-10 -5 0 5 10

**B.**

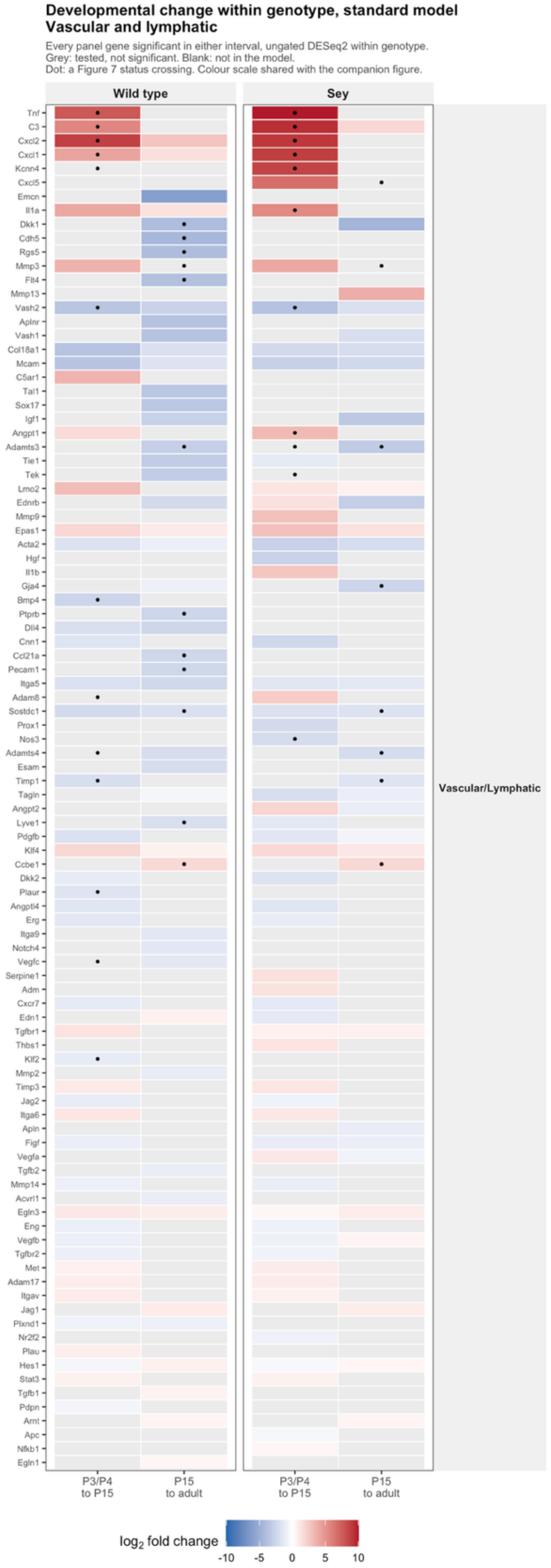
Developmental change within genotype, standard model
Vascular and lymphatic
Every panel gene significant in either interval, ungated DESeq2 within genotype.
Grey: tested, not significant. Blank: not in the model.
Dot: a Figure 7 status crossing. Colour scale shared with the companion figure.
Wild type
Sey
Tnf
C3
Cxcl2
Cxcl1
Kcnn4
Cxcl5
Emcn
Il1a
Dkk1
Cdh5
Rgs5
Mmp3
Flt4
Mmp13
Vash2
Aplnr
Vash1
Col18a1
Mcam
C5ar1
Tal1
Sox17
Igf1
Angpt1
Adamts3
Tie1
Tek
Lmo2
Ednrb
Mmp9
Epas1
Acta2
Hgf
Il1b
Gja4
Bmp4
Ptprb
Dll4
Cnn1
Ccl21a
Pecam1
Itga5
Adam8
Sostdc1
Prox1
Nos3
Adamts4
Esam
Timp1
Tagln
Angpt2
Lyve1
Pdgfb
Klf4
Ccbe1
Dkk2
Plaur
Angptl4
Erg
Itga9
Notch4
Vegfc
Serpine1
Adm
Cxcr7
Edn1
Tgfbr1
Thbs1
Klf2
Mmp2
Timp3
Jag2
Itga6
Apln
Figf
Vegfa
Tgfb2
Mmp14
Acvrl1
Egln3
Eng
Vegfb
Tgfbr2
Met
Adam17
Itgav
Jag1
Plxnd1
Nr2f2
Plau
Hes1
Stat3
Tgfb1
Pdpn
Arnt
Apc
Nfkb1
Egln1
Vascular/Lymphatic
P3/P4 to P15
P15 to adult
P3/P4 to P15
P15 to adult
log2 fold change
-10 -5 0 5 10

**Supplementary Figure 4. Developmental $\log_2$ fold changes under the replicate-aware model.** (A) Axon guidance and myelination panels; (B) vascular and lymphatic panel. Ungated within-genotype DESeq2 $\log_2$ fold changes for each developmental interval, with status crossings from Figure 7 marked.

Source data (Supplementary Table 19; the files below are archived at Zenodo, https://doi.org/10.5281/zenodo.23349739):

- Concordance/Figure7_DevelopmentalDE_Nerve_SourceData.csv
- Concordance/Figure7_DevelopmentalDE_Vascular_SourceData.csv

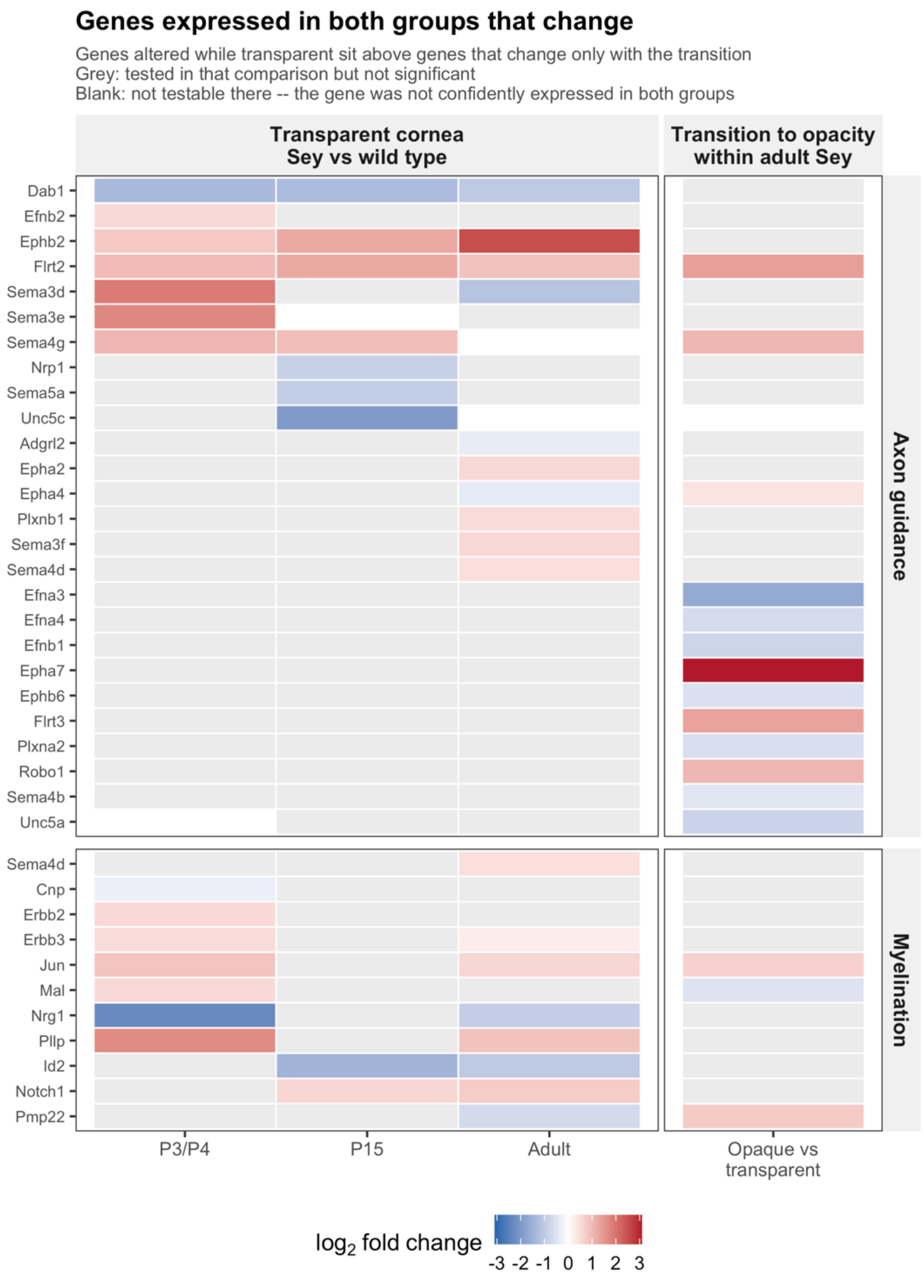

**Supplementary Figure 5**. **Quantitative change among expressed axon guidance and myelination panel genes.** Each tile is the unshrunk log2 fold change for a panel gene that was confidently expressed in both groups of the comparison and differed significantly (DESeq2; Benjamini–Hochberg adjusted P < 0.05, corrected within primary panel and comparison; no fold-change threshold). The first three columns compare transparent *Sey* with wild-type cornea at P3/P4 (n = 5 and 4), P15 (5 and 5) and adulthood (5 and 5); the fourth compares opaque with transparent adult *Sey* cornea (5 and 5). Red, higher in *Sey* or in the opaque cornea; blue, lower. Grey, tested but not significant; blank, not testable because the gene was not confidently expressed in both groups. A gene is shown if it changes significantly in at least one comparison: 37 gene-by-panel entries (26 of the 92 axon-guidance genes and 11 of the 25 myelination genes; Sema4d is on both panels). Genes altered while the cornea is transparent are placed above genes that change only with the transition to opacity (21 entries changed while transparent only, 6 in both, 10 with the transition only). Of the 57 significant changes, 34 are increases. *Ephb2* is higher in *Sey* at every age (+0.77, +1.24 and +2.54 log2) and *Flrt2* is higher in all four comparisons; the largest change is *Epha7* at +3.09 log2 in the opaque cornea, which sets the limit of the color scale. The color scale differs from that of Supplementary Figure 6. Quantitative changes in level are distinct from the categorical status differences of Figure 8B, which require no model. Pipeline Figure 9.

Source data (Supplementary Table 19; the files below are archived at Zenodo, https://doi.org/10.5281/zenodo.23349739):

- Figure9_SourceData.csv

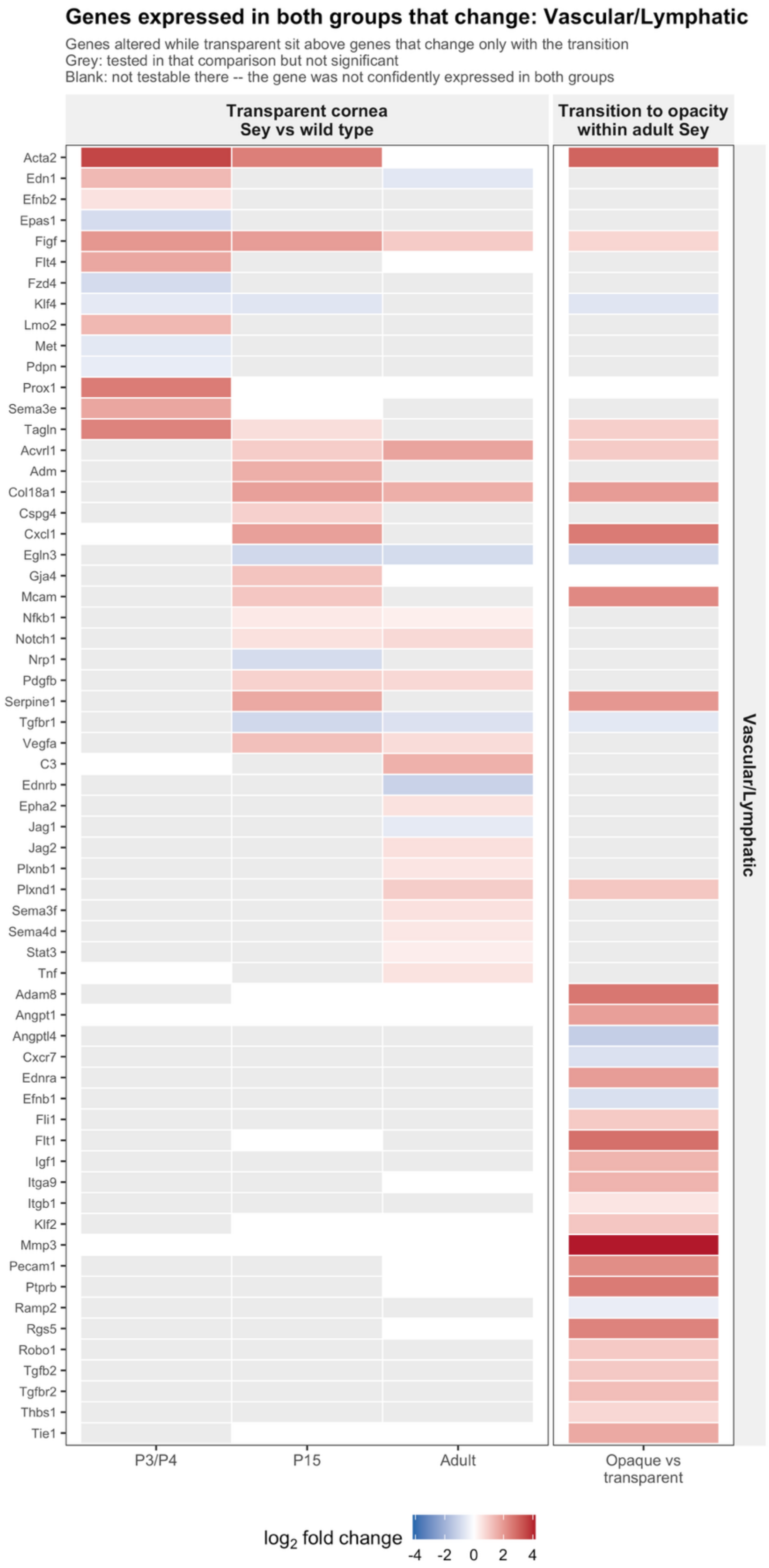

**Supplementary Figure 6. Quantitative change among expressed vascular and lymphatic panel genes.** Each tile is the unshrunk log2 fold change for a panel gene that was confidently expressed in both groups of the comparison and differed significantly (DESeq2; Benjamini–Hochberg adjusted P < 0.05, corrected within primary panel and comparison; no fold-change threshold). Columns compare transparent *Sey* with wild-type cornea at P3/P4, P15 and adulthood and, at right, opaque with transparent adult *Sey* cornea. Red, higher in *Sey* or in the opaque cornea; blue, lower. Grey, tested but not significant; blank, not testable because the gene was not confidently expressed in both groups. A gene is shown if it changes significantly in at least one comparison: 62 of the 177 panel genes. Genes altered while the cornea is transparent are placed above genes that change only with the transition to opacity (28 changed while transparent only, 12 in both, 22 with the transition only). Of the 88 significant changes, 67 are increases, including 27 of the 34 in the opacity comparison. *Figf* is changed in all four comparisons; the largest changes are *Mmp3* (+4.17 log2) and *Acta2* (+3.02) in the opaque cornea and *Acta2* at P3/P4 (+3.55), and the largest change sets the limit of the color scale, which differs from that of Supplementary Figure 5. Pipeline Figure 10

Source data (Supplementary Table 19; the files below are archived at Zenodo, https://doi.org/10.5281/zenodo.23349739):

Figure10_SourceData.csv

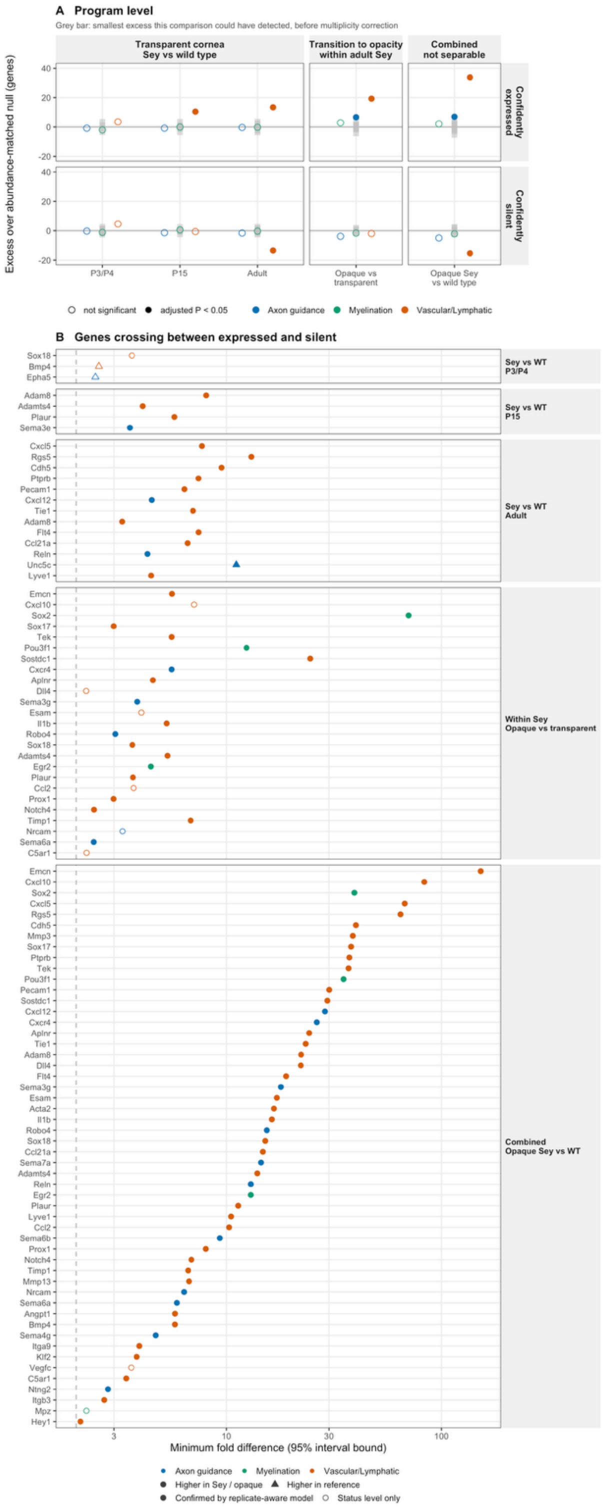

**Supplementary Figure 7. The combined comparison: opaque *Pax6*$^{Sey\text{-}Neu/+}$ cornea versus *Pax6*$^{+/+}$.** Constructed exactly as Figure 8, with a third block added at the right under a "Combined / not separable" strip. Opaque mutant and wild-type corneas differ in genotype and in transparency at once, so a difference between them cannot be attributed to either; the comparison is shown here for completeness and is excluded from Figure 8 for that reason. (A) Thirty panel-by-comparison tests are shown and eight reach significance. The combined block carries the largest excesses anywhere in the analysis; the vascular/lymphatic panel gains 33.7 confidently expressed genes and loses 15.4 confidently silent ones over its abundance-matched null, which is what summing two axes produces, not a stronger biological result.

(B) Ninety-seven gene-by-comparison status changes in 55 genes: 20 in the transparent block, 25 in the opacity block and 52 in the combined block, all of which are higher in the opaque mutant. Eighty-six of the 97 entries are also significant under a replicate-aware negative-binomial model (filled symbols); the other 11 (*Bmp4*, *C5ar1*, *Ccl2*, *Cxcl10*, *Dll4*, *Epha5*, *Esam*, *Mpz*, *Nrcam*, *Sox18* and *Vegfc*) are drawn open and are changes in status level only. Circles mark genes higher in *Sey* or in the opaque mutant and triangles genes higher in the reference group. Fourteen genes appear only in the combined comparison (*Acta2*, *Angpt1*, *Hey1, Itga9, Itgb3*, *Klf2*, *Mmp3*, *Mmp13*, *Mpz*, *Ntng2, Sema4g, Sema6b*, *Sema7a* and *Vegfc*) and the largest separation in the figure, *Emcn* at 152-fold, is a combined-block value. Nothing in this block can be assigned to genotype or to the loss of transparency, and none of it is quoted in the Results.

Point positions are not identical to Figure 8. Adding the combined comparison rescales the y axis of panel A and re-orders panel B, whose genes are ranked by the largest separation each achieves in any comparison; the underlying values are unchanged. The heatmaps are not reproduced for this comparison, because their color scale is taken from the largest effect present and would not be comparable with Supplementary Figures 5 and 6.

Source data (Supplementary Table 19; the files below are archived at Zenodo, https://doi.org/10.5281/zenodo.23349739):

- Generated by 08_figures.R with PAX6_FIG7_CONFOUNDED=TRUE
- Figure8_Confounded_CaptionNumbers.txt
- Figure8_Confounded_SourceData.csv.

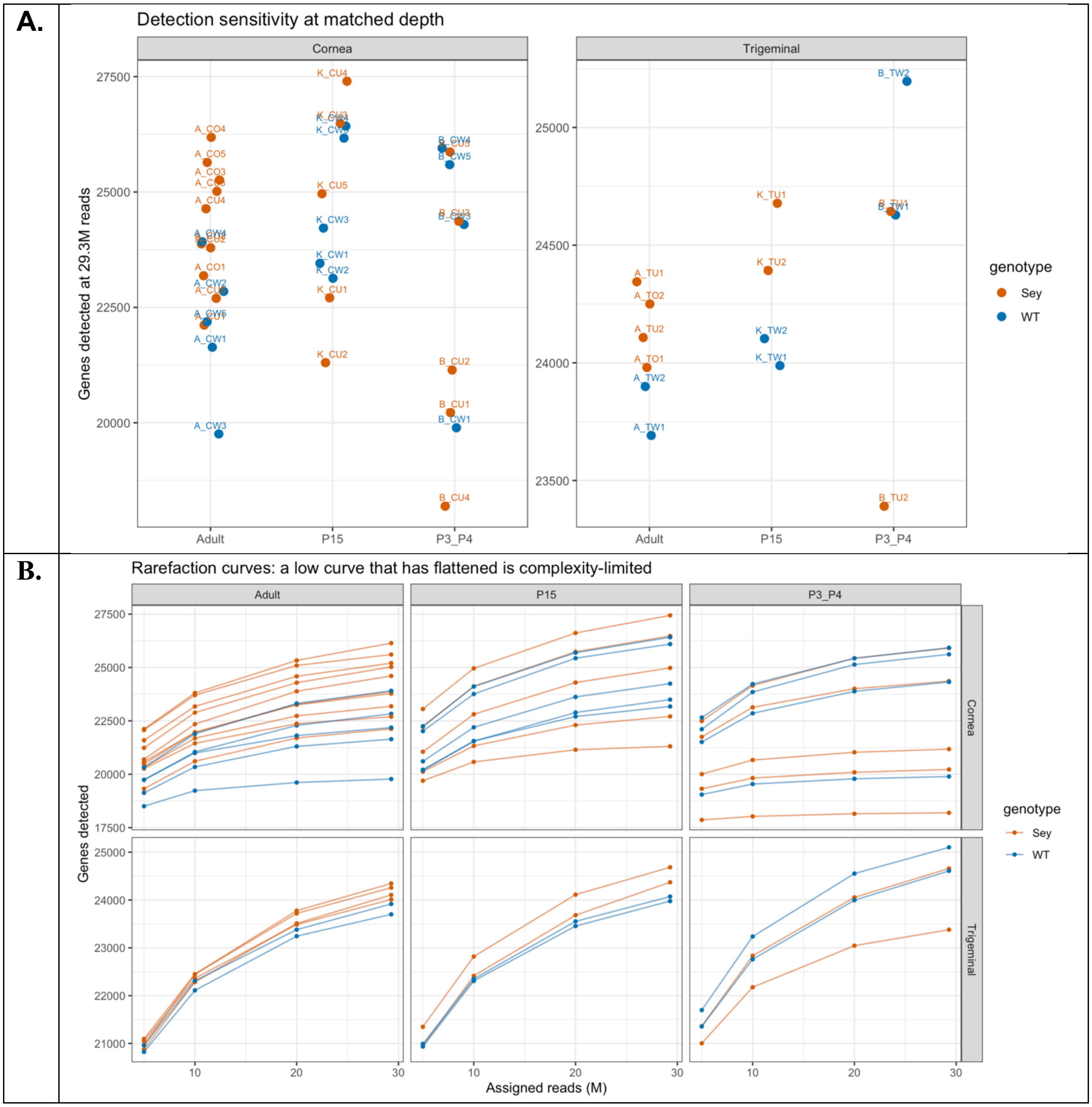

**Supplementary Figure 8. Library depth, rarefaction and detection sensitivity.** (A) Genes detected in each of the 48 libraries after every library was downsampled to the size of the smallest, 29.3 million assigned reads, so that detection can be compared at matched depth; cornea (left) and trigeminal ganglion (right), by age. Each point is one library, colored by genotype (Sey, orange; WT, blue) and labeled by library, with points offset horizontally for legibility. (B) Rarefaction curves for the same libraries, showing genes detected at 5, 10, 20 and 29.3 million assigned reads. A low curve that has flattened indicates a library limited by complexity, whereas a library that is merely shallow is still rising. Each library was judged against the median of its own tissue and age stratum (cornea, n = 9, 10 and 15 at P3/P4, P15 and adult; trigeminal, n = 4, 4 and 6). No library fell more than three median absolute deviations below its stratum median, and none was excluded. At matched depth, detection ranged from 18,192 to 27,399 genes in cornea and from 23,391 to 25,196 in trigeminal ganglion. Six corneal libraries (four Sey and two wild type) fell below 90% of their stratum median; the lowest, the P3/P4 Sey library B_CU4, detected 75% of its stratum median and its curve has flattened. One trigeminal library, B_TU2 (Sey, P3/P4), lay at the three-deviation threshold (robust $z = -2.96$) within a stratum of four, where the median absolute deviation is unstable. Rarefied counts were used only for this assessment and for no differential test.

Source data (Supplementary Table 20; the files below are archived at Zenodo, https://doi.org/10.5281/zenodo.23349739):

- QC_LibraryDepth_PerSample.csv
- QC_LibraryDepth_Series.csv.

### Supplementary Tables

Supplementary Tables 1–3 are in this file. Supplementary Table 4 (antibodies, kits and reagents) is in Supplementary Methods: Animals, Imaging and Nerve Quantification. Supplementary Tables 5–20 are provided as separate Excel files, each also as one CSV per sheet; their contents and source files are described below. WT, $Pax6^{+/+}$; Sey, $Pax6^{Sey\text{-}Neu/+}$; transparent, mixed and opaque are the corneal transparency classes (T, M, O) described in the Methods.

#### Overview of Supplementary Tables

| No. | Title | Format | Cited in |
|---|---|---|---|
| 1 | Corneas analyzed for TuJ1 labeling, by age, genotype and litter | This file | Results; Methods; Supplementary Methods (animals) |
| 2 | Transmission electron microscopy nerve fiber quantification | This file | Results; Supplementary Methods (animals) |
| 3 | Early postnatal CD31 immunolabeling | This file | Results; Discussion; Figure legends; Supplementary Methods (animals) |
| 4 | Antibodies, kits and reagents | Suppleme ntary Methods | Methods; Supplementary Methods (animals) |
| 5 | Developmental status crossings | Excel | Results; Supplementary Methods S10 |
| 6 | Developmental trajectories | Excel | Results; Supplementary Methods S10 |
| 7 | Concordance of status calls with replicate-aware models | Excel | Results; Methods; Supplementary Methods S12 |
| 8 | Genotype-by-age interaction tests | Excel | Results; Figure legends; Supplementary Methods S10 |
| 9 | Threshold and stability sensitivity | Excel | Results; Methods; Supplementary Methods S12 |
| 10 | Expression status and assessability | Excel | Results; Supplementary Methods S6, S7 |
| 11 | Categorical differences in expression status | Excel | Results; Supplementary Methods S7, S11 |
| 12 | Trigeminal ganglion pilot | Excel | Results; Supplementary Methods S14 |
| 13 | Sequencing library inventory and quality metrics | Excel | Methods; Supplementary Methods (animals); Supplementary Methods S2, S3, S5, S13 |
| 14 | Cross-dataset replication against GSE183742 | Excel | Methods; Supplementary Methods S13 |
| 15 | Curated gene panels and primary-panel assignment | Excel | Supplementary Methods S4 |
| 16 | Per-gene differential expression outcomes | Excel | Supplementary Methods S8 |
| 17 | Program-level tests | Excel | Supplementary Methods S9 |
| 18 | Read processing and quantification (54 libraries) | Excel | Supplementary Methods S3 |
| 19 | Source data for Figures 7–10 and supplementary figures | Excel | Supplementary Methods S10, S11 |
| 20 | Library depth and off-target tissue QC | Excel | Supplementary Methods S5 |

#### Supplementary Table 1. Corneas analyzed for TuJ1 labeling, by age, genotype and litter

Total mice and corneas used for TuJ1 immunolabeling and for TuJ1 quantification at each age, the number of litters from which they came, and the number of regions of interest (ROIs) quantified per cornea. Adult Sey corneas were classified as transparent, mixed or opaque. WT, $Pax6^{+/+}$; Sey, $Pax6^{Sey\text{-}Neu/+}$; transparent, mixed and opaque are the corneal transparency classes (T, M, O) described in the Methods.

| Age | Genotype | Mice labeled (n) | Litters (n) | Corneas labeled (n) | Mice quantified (n) | Corneas quantified (n) | ROIs per cornea (n) |
|---|---|---|---|---|---|---|---|
| **Newborn** | WT | 3 | 1 | 3 | 3 | 3 | 3 |
| | Sey | 4 | 1 | 4 | 3 | 3 | 3 |
| **1 week** | WT | 6 | 4 | 8 | 5 | 5 | 3 |
| | Sey | 8 | 4 | 10 | 5 | 5 | 3 |
| **2 weeks** | WT | 4 | 3 | 8 | 4 | 5 | 3 |
| | Sey | 4 | 3 | 8 | 4 | 5 | 3 |
| **4 weeks** | WT | 6 | 6 | 9 | 5 | 5 | 3 |
| | Sey | 8 | 6 | 12 | 5 | 5 | 3 |
| **Adult** | WT | 10 | 9 | 13 | 5 | 5 | 3 |
| | Sey, transparent | 10 | 6 | 14 | 5 | 5 | 3 |
| | Sey, mixed | 8 | 6 | 10 | 5 | 5 | 3 |
| | Sey, opaque | 5 | 4 | 8 | 5 | 5 | 3 |
| | Sey, not classified | 3 | 3 | 3 | 0 | 0 | 0 |

#### Supplementary Table 2. Transmission electron microscopy nerve fiber quantification

Number of myelinated nerve fibers among the nerve fiber clusters captured by transmission electron microscopy. Images from mixed and opaque Sey corneas are separated by the region imaged. WT, $Pax6^{+/+}$; Sey, $Pax6^{Sey\text{-}Neu/+}$; transparent, mixed and opaque are the corneal transparency classes (T, M, O) described in the Methods.

| Sample | Nerve fiber clusters captured (n) | Myelinated nerve fibers (n) |
|---|---|---|
| WT | 2 | 0 |
| Sey, transparent | 5 | 0 |
| Sey, mixed: transparent region | 2 | 0 |
| Sey, mixed: opaque region | 3 | 2 |
| Sey, opaque: central cornea | 10 | 4 |
| Sey, opaque: peripheral cornea | 19 | 11 |
| **Total** | **41** | **17** |

#### Supplementary Table 3. Early postnatal CD31 immunolabeling

Number of corneas used for CD31 immunolabeling and the number with vascular structures detected in the peripheral and central cornea (corneas with structures / corneas examined). WT, Pax6$^{+/+}$; Sey, Pax6$^{Sey\text{-}Neu/+}$; transparent, mixed and opaque are the corneal transparency classes (T, M, O) described in the Methods.

| Age | Genotype | Mice labeled (n) | Corneas labeled (n) | Peripheral cornea with vessels | Central cornea with vessels |
|---|---|---|---|---|---|
| **Newborn** | WT | 3 | 3 | 3/3 | 0/3 |
| | Sey | 3 | 3 | 2/3 | 0/3 |
| **1 week** | WT | 1 | 2 | 0/2 | 0/2 |
| | Sey | 3 | 3 | 1/3 | 0/3 |
| **2 weeks** | WT | 3 | 3 | 1/3 | 0/3 |
| | Sey | 7 | 7 | 7/7 | 3/7 |
| **3 weeks** | WT | 3 | 3 | 0/3 | 0/3 |
| | Sey | 5 | 5 | 5/5 | 3/5 |

#### Supplementary Tables provided as separate Excel files

Supplementary Tables 5–20 are provided as separate Excel files, each also as one CSV per sheet. Each description names the pipeline output file(s) that the table reproduces. The pipeline output files named in the descriptions of Tables 5–12, 14–17, 19 and 20 are archived at Zenodo (https://doi.org/10.5281/zenodo.23349739); Tables 13 and 18 (library metadata and processing quality control) are archived at https://doi.org/10.5281/zenodo.23350548.

##### Supplementary Table 5. Developmental status crossings.

Every gene whose expression status crossed between ages within a genotype, with the interval, direction, calls at both ages and bounded fold (sheet Status_crossings; Developmental_StatusCrossings.csv); the basis of Figure 7. Genes on more than one panel appear once per panel.

##### Supplementary Table 6. Developmental trajectories.

Sheet Trajectories: outcome and trajectory shape for every panel gene across the three ages, within each genotype (Developmental_Trajectories.csv). Sheet Steps_per_gene: differential expression for every panel gene at each adjacent age step within each genotype, with its interval, equivalence test and outcome (Developmental_Steps.csv). Sheet Steps_summary: counts of outcomes by panel, genotype and step (Developmental_Summary.csv).

##### Supplementary Table 7. Concordance of status calls with replicate-aware models.

Each status crossing and categorical call re-tested under an ungated DESeq2 model and edgeR quasi-likelihood, with the confirmation used to draw open symbols in Figures 7 and 8, and the “cannot assess” calls that reach significance under the ungated model. Sheet Age_axis_crossings: the 144 Figure 7 crossings against the ungated within-genotype model, with a verdict and the share of the pooled count supplied by the largest library (Concordance_AgeAxis.csv). Sheet Age_axis_cannot_assess: ungated calls at the categorical fold that are not status crossings, with the route the pipeline sent each down (Concordance_AgeAxis_StandardSignificant.csv). Sheet Genotype_axis_calls: every panel gene in every genotype comparison, with assessability, ungated verdict and largest-library share (Concordance_GenotypeAxis.csv), and Genotype_axis_summary: counts by comparison, route and verdict (Concordance_GenotypeAxis_Summary.csv). Sheet edgeR: the second engine’s results (Concordance_edgeR.csv). Sheet Flagged_library_calls: status calls whose higher side is led by a library flagged for off-target tissue in Supplementary Table 13, with that library’s share (Concordance_FlaggedLibraryCalls.csv).

##### Supplementary Table 8. Genotype-by-age interaction tests.

For each panel gene and pair of ages, the difference between genotype effects with its interval and outcome (sheet Interaction_tests; InteractionResults.csv): 483 gene-by-interval tests, of which 26 show a genotype effect that differs between ages and 7 are bounded as equivalent. The file has 564 rows because a gene on more than one panel appears once per panel.

##### Supplementary Table 9. Threshold and stability sensitivity.

Three sheets. Threshold_grid: the 3 × 3 grid of ON/OFF thresholds (0.5/0.1, 1/0.2 and 2/0.4 CPM) and minimum folds (1.5, 2 and 3) used to re-derive the status crossings, with the number of crossings and genes at each setting and the number of the 58 genes of Figure 7 that retain a crossing and keep their row group (Concordance_ThresholdStability.csv). Fold_sweep: the number of categorical calls at minimum bounded folds of 1, 1.5, 2, 3 and 5 (Categorical_FoldThresholdSensitivity.csv). Gate_cost: the number of calls reclassified by the leave-one-out stability gate in each group (GateCost_ByCell.csv); the Adult:Sey and Adult:Sey_T rows have identical values.

##### Supplementary Table 10. Expression status and assessability.

Sheet Expression_calls: expression status for every panel gene in every group (pooled CPM, pooled count, library size, exact Poisson limits, gated and ungated call; ExpressionCalls.csv). Sheet Assessability: assessability of every gene-by-comparison test, with the named reason where a gene could not be assessed (Assessability.csv). Sheets Expression_summary and Assessability_summary: counts by panel, group and call, and by panel, comparison and reason (ExpressionSummary.csv, Assessability_Summary.csv).

##### Supplementary Table 11. Categorical differences in expression status.

Sheet Categorical_genes: every gene called expressed in one group and not expressed in the other, or with Poisson intervals separated by at least twofold, with the bounded fold and its direction (CategoricalGenes.csv). A gene on more than one panel appears once per panel. The calls for the four comparisons of Figure 8B are separated from the combined comparison shown in Supplementary Figure 7. The axis column (opacity or age) labels the comparison family; the adult transparent-mutant versus wild-type comparison appears under both axes with identical rows, so counts should be taken from one axis.

##### Supplementary Table 12. Trigeminal ganglion pilot.

Five sheets. Panel_results: replicate-aware differential expression (DESeq2) for the three panels in the 14 trigeminal-ganglion libraries, two per group (Trigeminal_PanelResults.csv), with Panel_summary: outcomes by panel and comparison (Trigeminal_PanelSummary.csv). Sample_table: the per-library male fraction (Trigeminal_SampleTable.csv). Tissue_markers: expression of central and myelinating Schwann-cell marker genes (Trigeminal_TissueMarkers.csv). Genome_wide: the number of genes tested and significant genome-wide for each comparison, including the wild-type age-effect positive control (Trigeminal_GenomeWide.csv). Two libraries per group are below the minimum of three at which the three-state rule makes calls, so no expression-status calls were made for the ganglia (Supplementary Methods S14).

##### Supplementary Table 13. Sequencing library inventory and quality metrics.

All 48 libraries (34 cornea and 14 trigeminal ganglion), one row per library: sample ID, tissue, age, genotype, corneal transparency, sequencing batch and project, library and sequencing parameters, read pairs, trimming survival, alignment and assignment rates, duplication and GC content, library size (fragments assigned to genes in the count matrix deposited at GEO, GSE348661), genes detected, detection-sensitivity flag, inferred sex composition, the sequencing provider's RNA quality values, and the pigment-cell and lymphoid contamination indices used in the sensitivity analysis. Each library is a pool, and the table also gives the animals, litters (Same, all animals from one litter; Different / n, animals from n litters), and corneas or ganglia per pool and the recorded donor sex (corneas from male : female donors, not mice; not recorded at P3/P4 and P15). Trigeminal ganglia came from the same animals as the corresponding cornea pool. Sex composition of every pool was also inferred from Xist and Y-linked gene expression: sex_inferred_male_frac is a relative index from 0 to 1 scaled within tissue and age group, not an absolute male proportion, and pool_sex_class_inferred is all female (Y-gene CPM < 1), all male (Xist CPM < 2) or mixed; the inferred class agreed with the recorded sex in all 15 adult cornea pools. Raw FASTQ file names and MD5 checksums and the deposited processed count matrix are listed for each library. Replicate 2 of the P3/P4 wild-type cornea (6 mice, same litter) has no library: its total RNA did not meet the sequencing provider's integrity threshold (RIN 1.8) and the trigeminal ganglion library from the same animals was sequenced; 28 RNA samples were submitted in 2025, and 27 passed the provider's RNA QC and were sequenced. Differential expression models normalised counts by the DESeq2 median-of-ratios method, with size factors estimated within each model, so size factors are not tabulated here. This table is the authoritative record of the pool composition, age, recorded donor sex and raw-file MD5 checksums of the 48 libraries.

**Library quality and off-target tissue.** Read pairs, trimming survival, alignment and assignment rates are from Trimmomatic, HISAT2 and featureCounts (fragment counting, unstranded); duplication and GC are FastQC values for raw read 1. RIN and RNA input are from the sequencing provider's sample QC (2025: submitted RNA, Agilent 5400; 2026: RNA extracted by the provider from frozen tissue). Detection-sensitivity flags are from the rarefied depth survey (soft, <90% of stratum median; hard, robust $z < -3$). Off-target tissue was scored per library from the marker genes used for the GSE183742 comparison: melanocyte-lineage genes (Tyr, Tyrp1, Dct, Mlana; reported as a summed index), Rho (retina), Cryaa (lens) and Myoc (iridocorneal angle), with Kera and Krt12 as stromal and epithelial controls, all as CPM of assigned fragments. Lymphoid cell content is reported as a summed index (Cd3e, Cd19, Cd79a, Cd79b, Ms4a1), with Ptprc (pan-leukocyte) given separately. A library is flagged when an index exceeds ten times the median of same-age cornea libraries (opaque corneas excluded from the reference) with at least two constituent genes elevated, and noted as "trace" at five- to ten-fold. Retinal (K_CU4, A_CW1; Rho ≤ 46 CPM) and lens (A_CO1, B_CU2; Cryaa ≤ 144 CPM) signals correspond to well under 1% of library RNA and no panel gene is retina- or lens-restricted; the pigment-cell and Myoc signals in the 2026 opaque adult corneas (A_CO3–A_CO5) reflect iris adherent to opaque regions at dissection and are retained and taken into account in the opacity-axis comparisons. The 2025 opaque libraries carry smaller signals of the same kind: A_CO1 (iris pigment trace, 8-fold; iridocorneal angle, Myoc 16-fold) and A_CO2 (angle trace, 5-fold). A_CO4 and A_CO5 are also epithelium-poor (Krt12 and Aldh3a1 at least 5- and 9-fold below the reference, respectively).

Flags are listed per library in the contamination_flags column; K_CW4, A_CU1 and A_CO4 are also stroma-poor by the criterion applied to GSE183742. Trigeminal libraries were not scored; their Tyrp1 signal is developmental, not melanocytic. K_CU3 (transparent Sey cornea at P15) was flagged for a pigment-cell index 12-fold above the median of same-age corneas (Tyr, Tyrp1, Dct; consistent with iris) and was retained; it is not part of the adult opacity-axis comparisons. No library was excluded.

##### Supplementary Table 14. Cross-dataset replication against GSE183742.

Eight sheets (CrossDataset_*.csv). WT_status_concordance and WT_crosstab: wild-type status concordance. WT_null_distribution: its abundance-matched null (2,000 draws). Direction and Direction_crosstab: the direction comparison. Direction_null_distribution: its size-matched null (2,000 draws). Dissection_margin: the dissection-margin marker check. GSE183742_off_target: the six GSE183742 libraries judged by the off-target rule applied to this study (Supplementary Table 13).

##### Supplementary Table 15. Curated gene panels and primary-panel assignment.

The axon guidance (92), myelination (25) and vascular/lymphatic (177) panels (263 unique genes) with Ensembl IDs, source of curation, membership of more than one panel, and the primary panel used for multiplicity correction (sheets Axon_guidance_panel, Myelination_panel and Vascular_lymphatic_panel: AxonGuidance_Master_List.csv, Myelination_Master_List.csv, VascularLymphatic_Master_List.csv; sheet Primary_panel_assignment: SharedGenePrimaryPanel.csv). Sheet Shared_gene_assignment gives the derivation of the primary panel for each gene on more than one panel (SharedGeneAssignment.csv).

##### Supplementary Table 16. Per-gene differential expression outcomes.

Sheet DE_results: unshrunk $\log_2$ fold change, standard error, 95% confidence interval, raw and adjusted P value, equivalence-test P value, outcome (changed, no change detected, undetermined), correction panel and family size for every testable gene in every comparison (DE_Results.csv). The transparent-mutant versus wild-type adult comparison appears under both models; the developmental-model row is a consistency check and is not reportable. Further sheets: DE_summary (outcome counts by panel and comparison; DE_Summary.csv), Adult_omnibus_LRT (likelihood-ratio test across the adult groups; AdultOmnibusLRT.csv), Hinge_check (transparent Sey and all Sey against wild type in adults; HingeCheck.csv), Testability_audit (genes testable, tested and lost to pre-filtering for each comparison; TestabilityAudit.csv) and Cooks_filtering (genes flagged by Cook's distance per model and library; CooksFiltering.csv).

##### Supplementary Table 17. Program-level tests.

For each panel and comparison, the observed difference in confidently expressed and confidently silent genes, the abundance-matched permutation null (20,000 permutations), the detectable limit, and raw and adjusted P values (ProgramLevel.csv). Tests for the four comparisons of Figure 8A (24 tests) are distinguished from the combined comparison shown in Supplementary Figure 7 (30 tests in all).

##### Supplementary Table 18. Read processing and quantification (54 libraries).

Four sheets. Libraries: read pairs, trimming survival, alignment and assignment rates, and fragments in the count matrix for the 54 libraries processed together (48 from this study and 6 from GSE183742), from reprocessing the raw reads with the released upstream pipeline (run reprocess_2026-09, pipeline commit 9182fd5). The reprocessed matrix is the matrix of record deposited in GEO (GSE348661) and used for every result; the sheet compares it with the earlier August 2026 matrix, which differs by at most 31 fragments per library. Strandedness: measured library strandedness for each dataset. Software and reference: tool versions and reference-file checksums. Raw file checksums: raw FASTQ file names and MD5 checksums. For the 48 libraries of this study, the read pairs, trimming, alignment, assignment, matrix-size and checksum values here are the same as those in Supplementary Table 13, which remains the authoritative record for this study's libraries; this table adds the GSE183742 libraries and the processing detail.

##### Supplementary Table 19. Source data for Figures 7–10 and the supplementary figures.

The values plotted in each figure panel, one sheet per figure: Figure7, Figure8, Figure9 and Figure10 (Figure7_SourceData.csv, Figure8_SourceData.csv, Figure9_SourceData.csv, Figure10_SourceData.csv); Figure8_Confounded, the combined comparison shown in Supplementary Figure 7 (Figure8_Confounded_SourceData.csv); and Figure7_DevDE_Nerve and Figure7_DevDE_Vascular, the supplementary developmental heatmaps (Figure7_DevelopmentalDE_Nerve_SourceData.csv, Figure7_DevelopmentalDE_Vascular_SourceData.csv).

##### Supplementary Table 20. Library depth and off-target tissue quality control.

Pipeline outputs behind the detection-sensitivity and off-target flags in Supplementary Table 13, one row per library: Contamination_indices, the off-target tissue and stromal and epithelial indices judged against same-age corneal libraries (Contamination_Indices.csv); Depth_per_sample, detected and effective genes per library (QC_LibraryDepth_PerSample.csv); Depth_pool_share, each library's share of its pooled reads (QC_LibraryDepth_PoolShare.csv); and Depth_rarefaction_series, genes detected at each rarefied depth (QC_LibraryDepth_Series.csv).
